# *C. elegans* toxicant responses vary among genetically diverse individuals

**DOI:** 10.1101/2022.07.19.500602

**Authors:** Samuel J. Widmayer, Timothy A. Crombie, Joy N. Nyaanga, Kathryn S. Evans, Erik C. Andersen

**Author notes:** Corresponding author: Erik C. Andersen Department of Molecular Biosciences Northwestern University 4619 Silverman Hall 2205 Tech Drive Evanston, IL 60208. Emails: Sam Tim Joy Katie Erik.

## Abstract

Comprehensive chemical hazard risk evaluations require reproducible, efficient, and informative experimental workflows in tractable model systems that allow for high replication within exposure cohorts. Additionally, the genetic variability of toxicant responses among individuals in humans and mammalian models requires practically untenable sample sizes. *Caenorhabditis elegans* is a premier toxicology model that has revolutionized our understanding of cellular responses to environmental pollutants and boasts robust genomic resources and high levels of genetic variation across the species. In this study, we performed dose-response analysis across 23 environmental toxicants using eight *C. elegans* strains representative of species-wide genetic diversity. We observed substantial variation in EC10 estimates and slope parameter estimates of dose-response curves of different strains, demonstrating that genetic background is a significant driver of differential toxicant susceptibility. We also showed that, across all toxicants, at least one *C. elegans* strain exhibited a significantly different EC10 or slope estimate compared to the reference strain, N2 (PD1074), indicating that population-wide differences among strains are necessary to understand responses to toxicants. Moreover, we quantified the heritability of responses to each toxicant dose and observed a correlation between the dose closest to the species-agnostic EC10 estimate and the dose that exhibited the most heritable response. Taken together, these results provide robust evidence that heritable genetic variation explains differential susceptibility across an array of environmental pollutants and that genetically diverse *C. elegans* strains should be deployed to aid high-throughput toxicological screening efforts.

## INTRODUCTION

Hazard risk assessment of environmental chemicals is a top priority of toxicological research. Over 350,000 chemicals are currently registered for use and production globally, of which tens of thousands are either confidential or ambiguously described (Wang et al., 2020). This staggering rate of production, paired with traditional means of hazard safety testing, which typically uses mammalian or cell-based methods of response evaluation, means that human populations are exposed to a complex array of xenobiotic compounds with virtually unknown risk levels. Although approaches to hazard risk assessments using mammalian systems have translational appeal, they often suffer from low statistical power because of necessarily limited sample sizes. These approaches are also time-consuming and economically costly (Tralau et al., 2012), drastically reducing their potential for thorough risk assessment of a growing, sometimes multifactorial, collection of chemical exposures (Brooks et al., 2020). Most importantly, meta-analyses estimate that rodent systems predict human toxic effects approximately 50% of the time (Hartung, 2009; Knight et al., 2009), suggesting that chemical risk assessment requires a more integrative approach.

*Caenorhabditis elegans* is a free-living nematode roundworm that can be cheaply reared in large samples in a matter of days, vastly accelerating the pace and scale at which hazard risk evaluations can be performed compared to most vertebrate models. Furthermore, studies using *C. elegans* provide data from whole animals with intact neuromuscular, endocrine, digestive, and sensory systems unlike popular *in vitro* systems. *C. elegans* is a powerful toxicology model that unites toxicologists with molecular geneticists so that expertise in routes of chemical exposure, internal dosage-specific effects, tissue distribution, and chemical metabolism is combined with expertise in DNA damage, oxidative and osmotic stress, and regulation of apoptosis and necrosis (Boyd et al., 2012; Hartman et al., 2021). All three phases of xenobiotic metabolism are present in *C. elegans*, though the conservation of specific gene families within each phase, such as the cytochromes P450, UDP-glucuronosyltransferases (UGTs), sulfotransferase enzymes (SULTs), and ATP-binding cassette (ABC) transporters (Hartman et al., 2021) have important differences. In addition to being inexpensive and easy to use, *C. elegans* responses to dozens of chemicals more accurately predict responses in rabbits and rats compared to zebrafish models (Boyd et al., 2016). Furthermore, meta-analyses indicate that rank-ordered toxicant sensitivity in several rodent models correlates with responses in *C. elegans* (Hunt, 2017). Therefore, toxicity assessments in *C. elegans* provide an alternative to vertebrate models with significantly greater scalability and potential to accelerate the characterization of molecular targets of chemical exposures.

One approach to account for intra- and inter-species variation in toxicant responses is to use uncertainty factors (UFs) to translate a hazard’s point of departure (POD) between species with distinct exposure routes and pharmacokinetic and pharmacodynamic capacities (Piersma et al., 2011). Although this approach has limited biases based on the compound class or populations being compared, POD calculations alone fail to directly account for heritable genetic variation between individuals - variance in susceptibility that can be explained by genetic differences that segregate among individuals in a population (Zeise et al., 2013). Failing to account for these differences leads to UFs serving as an imprecise proxy for within-species variation in risk because the process is agnostic to observed ranges of susceptibility in genetically diverse individuals. Evaluations that are able to quantify the contributions of genetics to toxicant response variation lay the foundation for quantitative genetic dissection, with the specific goal of revealing novel mechanisms of toxicant susceptibility by identifying risk alleles. Wild strains of *C. elegans* harbor rich genetic variation (Andersen et al., 2012; Cook et al., 2017; Lee et al., 2021) and, by combining quantitative and molecular genetic approaches, offer the opportunity to discover genetic modifiers of toxicant susceptibility (Andersen et al., 2015; Bernstein et al., 2019; Evans et al., 2020; Zdraljevic et al., 2019). Quantifying the effects of genetics on toxicant susceptibility in *C. elegans* is an important step towards a full characterization of chemical hazard risk because the additive effects of conserved genes can help us understand novel toxicant response biology in humans. Additionally, the effects of these specific alleles can be dissected in *C. elegans* using genetic crosses and state-of-the-art molecular methods much faster than in mammalian systems.

In this study, we performed dose-response analysis across 25 toxicants representing distinct chemical classes using eight strains of *C. elegans* representative of species-wide genetic diversity. We used a high-throughput imaging platform to assay development after exposing arrested first larval stage animals to each toxicant in a dose-dependent manner and used custom software (Di Tommaso et al., 2017; Nyaanga et al., 2021; Wählby et al., 2012) to measure phenotypic responses to each compound. By estimating dose-response curves for each toxicant and fitting strain-specific model parameters, we demonstrated that natural genetic variation is a key determinant of toxicant susceptibility in *C. elegans*. Moreover, we showed that the specific alleles that segregate between the eight strains in our cohort exert additive effects on toxicant susceptibility, which implies that quantitative genetic dissection of these responses has the potential to yield novel genetic loci underlying toxicant susceptibility. Taking these observations together, we propose that leveraging standing natural genetic variation in *C. elegans* is a necessary and complementary tool for high-throughput hazard risk assessments in translational toxicology.

## METHODS

### Strains

The eight strains used in this study (PD1074, CB4856, MY16, RC301, ECA396, ECA36, ECA248, XZ1516) are available from the *C. elegans* Natural Diversity Resource (CeNDR) (Cook et al., 2017). Isolation details for the eight strains are included on CeNDR. Of the eight strains used, two (PD1074 and ECA248) are referred to by their isotype names (N2 and CB4855, respectively). Prior to measuring toxicant responses, all strains were grown at 20°C on 6 cm plates made with modified nematode growth medium (NGMA) that contains 1% agar and 0.7% agarose to prevent animals from burrowing (Andersen et al., 2014). The NGMA plates were spotted with OP50 *Escherichia coli* as a nematode food source. All strains were propagated for three generations without starvation on NGMA plates prior to toxicant exposure. The specific growth conditions for nematodes used in the high-throughput toxicant response assay are described below (*see Methods, High-throughput toxicant response assay*).

### Nematode food preparation

We prepared a single batch of HB101 *E. coli* as a nematode food source for all assays in this study. In brief, we streaked a frozen stock of HB101 *E. coli* onto a 10 cm Luria-Bertani (LB) agar plate and incubated it overnight at 37°C. The following morning, we transferred a single bacterial colony into a culture tube that contained 5 ml of 1x Horvitz Super Broth (HSB). We then incubated that starter culture and a negative control (1X HSB without bacteria) for 18 hours at 37°C with shaking at 180 rpm. We then measured the OD_600_ value of the starter culture with a spectrophotometer (BioRad, smartspec plus), calculated how much of the 18-hour starter culture was needed to inoculate a one liter culture at an OD_600_ value of 0.001, and used it to inoculate 14 4 L flasks that each contained one liter of pre-warmed 1x HSB. We grew those 14 cultures for 15 hours at 37°C with shaking at 180 rpm until they were in the early stationary growth phase (Supplemental Figure 1A). We reasoned that food prepared from cultures grown to the early stationary phase (15 hours) would be less variable than food prepared from cultures in the log growth phase. At 15 hours, we removed the culture flasks from the incubator and transferred them to a 4°C walk-in cold room to arrest growth. We then removed the 1X HSB from the cultures by three repetitions of pelleting the bacterial cells with centrifugation, disposing of the supernatant, and resuspending the cells in K medium. After the final wash, we resuspended the bacterial cells in K medium and transferred them to a 2 L glass beaker. We measured the OD_600_ value of this bacterial suspension, diluted it to a final concentration of OD_600_ 100 with K medium, aliquoted it to 15 ml conicals, and froze the aliquots at −80°C for use in the dose response assays.

### Toxicant stock preparation

We prepared stock solutions of the 25 toxicants using either dimethyl sulfoxide (DMSO) or water depending on the toxicant’s solubility. The exact sources, catalog numbers, stock concentrations, and preparation notes for each of the toxicants are provided (Supplemental Table 1). Following preparation of the toxicant stock solutions, they were aliquoted to microcentrifuge tubes and stored at −20°C for use in the dose response assays.

### High-throughput toxicant dose response assay

For each replicate assay, populations of each strain were passaged for three generations, amplified, and bleach-synchronized in triplicate (Figure 1A). We replicated the bleach synchronization to control for variation in embryo survival and subsequent effects on developmental rates that could be attributed to bleach effects (Porta-de-la-Riva et al., 2012) (Figure 2A). Following each bleach synchronization, we dispensed approximately 30 embryos into the wells of 96-well microplates in 50 µL of K medium (Boyd et al., 2012). We randomly assigned strains to rows of the 96-well microplates and varied the row assignments across the replicate bleaches. We prepared four replicate 96-well microplates within each of three bleach replicates for each toxicant and control condition tested in the assay. We then labeled the 96-well microplates, sealed them with gas permeable sealing film (Fisher Cat #14-222-043), placed them in humidity chambers, and incubated them overnight at 20°C with shaking at 170 rpm (INFORS HT Multitron shaker). The following morning, we prepared food for the developmentally arrested first larval stage animals (L1s) using frozen aliquots of HB101 *E. coli* suspended in K medium at an optical density at 600 nm (OD_600_) of 100 (see Methods, Nematode food preparation). We thawed the required number of OD_600_ 100 HB101 aliquots at room temperature, combined them into a single conical tube, diluted them to OD_600_ 30 with K medium, and added Kanamycin at 150 µM to inhibit further bacterial growth and prevent contamination. Working with a single toxicant at a time, we then transferred a portion of the OD_600_ 30 food mix to a 12-channel reservoir, thawed an aliquot of toxicant stock solution at room temperature (*see methods, Toxicant stock preparation*), and diluted the toxicant stock to a working concentration. The toxicant working concentration was set to the concentration that would give the highest desired dose when added to the 96-well microplates at 1% of the total well volume. We then performed a serial dilution of the toxicant working solution using the same diluent used to make the stock solution (Figure 1C). The dilution factors ranged from 1.1 to 2 depending on the toxicant used, but all serial dilutions had 12 concentrations, including a 0 µM control. Using a 12-channel micropipette, we added the toxicant dilution series to the 12-channel reservoir containing the food mix at a 3% volume/volume ratio. Next, we transferred 25 µL of the OD_600_ 30 food and toxicant mix from the 12-channel reservoir into the appropriate wells of the 96-well microplates to simultaneously feed the arrested L1s at a final HB101 concentration of OD_600_ 10 and expose them to toxicant at one of 12 levels of the dilution series. We chose to feed at a final HB101 concentration of OD_600_ 10 because nematodes consistently developed to L4s after 48 hours of feeding at 20°C (Supplemental Figure 1B). Immediately after feeding, we sealed the 96-well microplates with a gas permeable sealing film (Fisher Cat #14-222-043), returned them to the humidity chambers, and started a 48-hour incubation at 20°C with shaking at 170 rpm. The remainder of the 96-well microplates were fed and exposed to toxicants in the same manner. After 48 hours of incubation in the presence of food and toxicant, we removed the 96-well microplates from the incubator and treated the wells with sodium azide (325 µL of 50 mM sodium azide in 1X M9) for 10 minutes to paralyze and straighten the nematodes. We then immediately acquired images of nematodes in the microplates using a Molecular Devices ImageXpress Nano microscope (Molecular Devices, San Jose, CA) with a 2X objective (Figure 1D). We used the images to quantify the development of nematodes in the presence of toxicants as described below (*see Methods, Data collection, and Data cleaning*).

**Figure 1:**
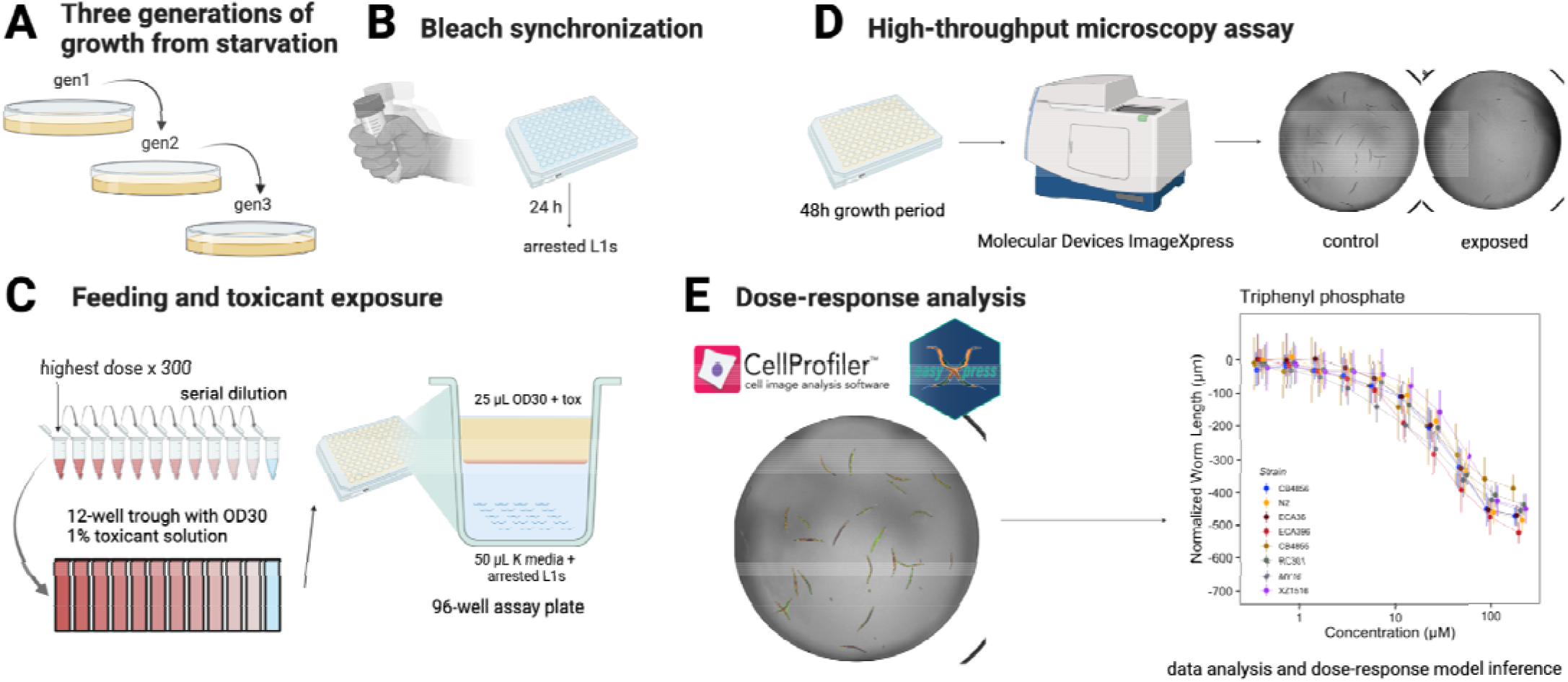
High-throughput microscopy assay enables rapid analysis of *C. elegans* toxicant responses. Detailed descriptions of A) through D) can be found in *Methods; High throughput toxicant dose response assay*. Detailed descriptions of E) can be found in *Methods; Data collection, Data cleaning, LOAEL inference, Dose-response model estimation.* Created with BioRender.com.

**Figure 2:**
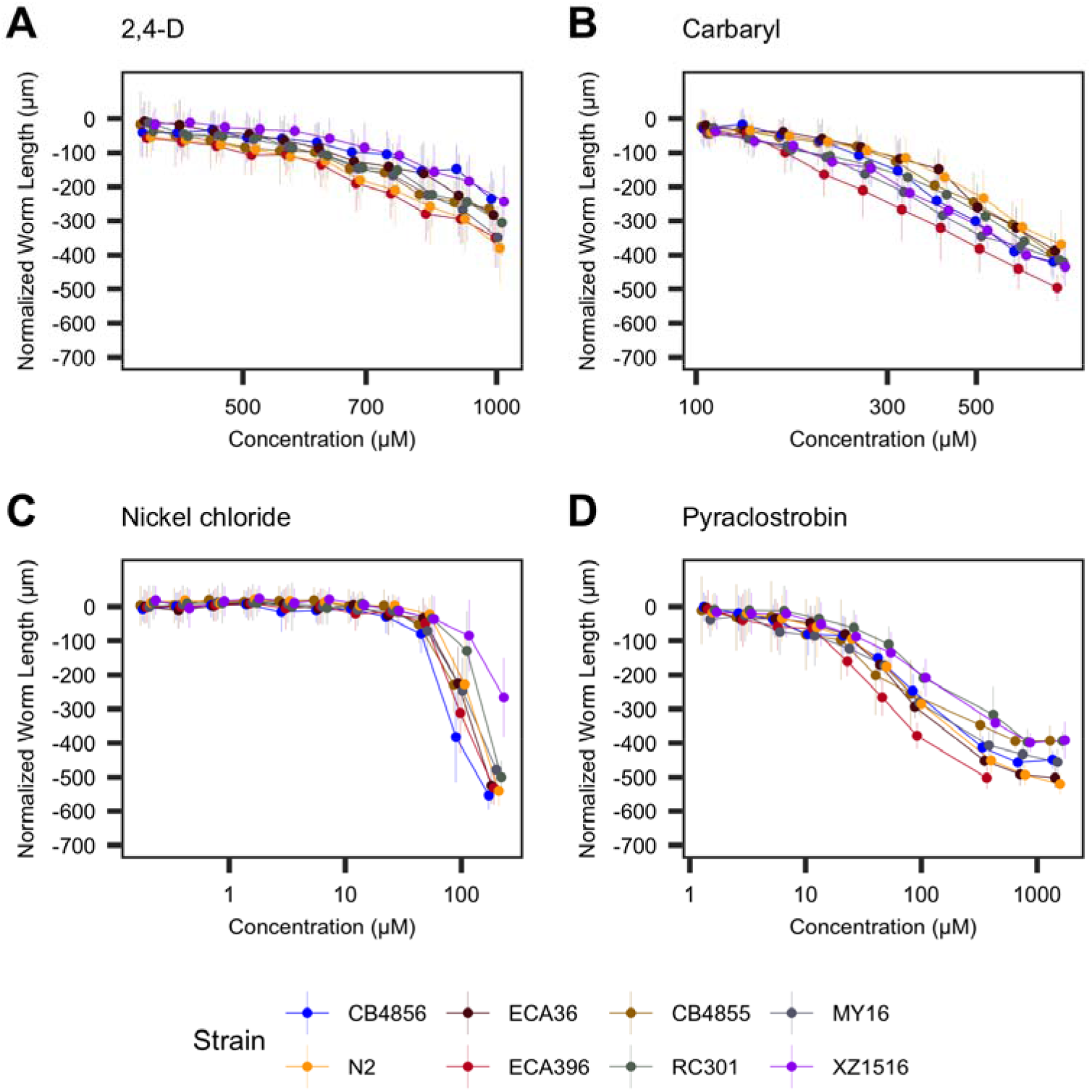
Toxicant responses vary among genetically diverse *C. elegans* strains. Normalized length measurements for each strain at each toxicant dose are shown on the y-axis, and the concentration of each toxicant is shown on the x-axis. Each dose-response curve is colored according to the strain. We observed a wide range of responses that can be combined into four general groups: A) subtle responses with little variation among strains, *e.g.*, 2,4-D; B) subtle responses with moderate variation among strains, *e.g.*, carbaryl; C) strong responses with little variation among strains, *e.g.*, nickel chloride (though for nickel chloride, strain variation is high at high doses, *see* Figure 5); and D) strong responses with moderate variation among strains, *e.g.*, pyraclostrobin.

### Data collection

We wrote custom software packages designed to extract animal measurements from images collected on the Molecular Devices ImageXpress Nano microscope (Figure 1E). CellProfiler is a widely used software program for characterizing and quantifying biological data from image-based assays (Carpenter et al., 2006; Kamentsky et al., 2011; McQuin et al., 2018). A collection of CellProfiler modules known as the WormToolbox were developed to extract morphological features of individual *C. elegans* animals from images from high-throughput *C. elegans* phenotyping assays like the one we use here (Wählby et al., 2012). We estimated worm models and wrote custom CellProfiler pipelines using the WormToolbox in the GUI-based instance of CellProfiler. We then wrote a Nextflow pipeline (Di Tommaso et al., 2017) to run command-line instances of CellProfiler in parallel on the Quest High Performance Computing Cluster (Northwestern University) because each experimental block in this study produced many thousands of well images. This workflow can be found at https://github.com/AndersenLab/cellprofiler-nf. Our custom CellProfiler pipeline generates animal measurements by using four worm models: three worm models tailored to capture animals at the L4 larval stage, in the L2 and L3 larval stages, and the L1 larval stage, respectively, as well as a “multi-drug high dose” (MDHD) model, to capture animals with more abnormal body sizes caused by extreme toxicant responses. We used *R/easyXpress* (Nyaanga et al., 2021) to filter measurements from worm objects within individual wells that were statistical outliers and to parse measurements from multiple worm models down to single measurements for single animals. These measurements comprised our raw dataset.

### Data Cleaning

All data management and statistical analyses were performed using the R statistical environment (version 4.0.4). Our high-throughput imaging platform produced thousands of images across each experimental block. It is unwieldy to manually curate each individual well image to assess the quality of animal measurement data. Therefore, we took several steps to clean the raw data using heuristics indicative of high-quality animal measurements suitable for downstream analysis.

1. We began by censoring experimental blocks for which the coefficient of variation (CV) of the number of animals in *control wells* was greater than 0.6 (Supplemental Figure 2A). Experiments containing wells that meet this criterion in control wells are expected to produce less precise estimates of animal lengths in wells in which animals have been exposed to chemicals that typically increase the variance of the body length trait (Supplemental Figure 2B).
2. We then reduced the data to wells containing between five and thirty animals, under the null hypothesis that the number of animals is an approximation of the expected number of embryos originally titered into wells (approximately 30). This filtering step screened for two problematic features of well images in our experiment. First, given that our analysis relied on well median animal length measurements, we excluded wells with less than five animals to reduce sampling error. Second, insoluble compounds or bacterial clumps were often identified as animals by CellProfiler (Supplemental Figure 3) and would vastly inflate the well census and spuriously deflate the median animal length in wells containing high concentrations of certain toxicants.
3. After the previous two data processing steps, we removed statistical outlier measurements within each concentration for each strain for every toxicant to reduce the likelihood that statistical outliers influence dose-response curve fits.
4. Next, we removed measurements from all doses of each toxicant that were no longer represented in at least 80% of the independent assays because of previous data filtering steps, or had fewer than 10 measurements per strain.
5. Finally, we normalized the data by (1) regressing variation attributable to assay and technical replicate effects and (2) normalizing these extracted residual values with respect to the average control phenotype. For each compound, we estimated a linear model using the raw phenotype measurement as the response variable and both assay and technical replicate identity as explanatory variables following the formula *median_wormlength_um ∼ Metadata_Experiment + bleach* using the *lm()* function in base R. We then extracted the residuals from this linear model for each dose and subtracted normalized phenotype measurements in each dose from the mean normalized phenotype in control conditions. These normalized phenotype measurements were used in all downstream statistical analyses.

### LOAEL inference

We determined the lowest observed adverse effect level (LOAEL) for each compound by performing a one-way analysis of variance using the normalized phenotype measurements as a response variable and toxicant dosage as an explanatory variable. We then performed a Tukey *post hoc* test, filtered to only comparisons to control doses, and determined the lowest dose that exhibited a significantly different phenotypic response as distinguished by an adjusted *p*-value less than 0.05. This analysis was performed on all phenotype measurements, as well as for each strain individually to determine if genetic background differences explain differences in LOAEL for each toxicant.

### Dose-response model estimation and statistics

We estimated overall and strain-specific dose-response models for each compound by fitting a log-logistic regression model using *R/drc* (Ritz et al., 2015). The log-logistic model that we used specified four parameters: *b*, the slope of the dose-response curve; *c*, the upper asymptote of the dose-response curve; *d*, the lower asymptote of the dose-response curve; and *e*, the specified effective dose. This model was fit to each compound using the *drc::drm()* function with strain specified as a covariate for parameters *b* and *e*, allowing us to estimate strain-specific dose-response slopes and effective doses, as well as a specified lower asymptote *d* at −600, which is the theoretical normalized length of animals at the L1 larval stage. We used the *drc::ED()* function to extract strain-specific EC10 values, and extracted the strain-specific slope values using base R. We quantified the relative susceptibilities of each strain pair for each compound based on their estimated EC10 values using the *drc::EDcomp()* function, which uses an approximate *F*-test to determine whether the variances (represented by delta-specified confidence intervals) calculated for each strain-specific dose response model’s *e* parameter estimates are significantly different. We quantified the relative slope steepness of dose-response models estimated for each strain within each compound using the *drc::compParm()* function, which uses a *z*-test to compare means of each *b* parameter estimate. Results shown are filtered to just comparisons against PD1074 dose-response parameters (Figures 2 and 3), and significantly different estimates in both cases were determined by correcting to a family-wise type I error rate of 0.05 using Bonferroni correction. To determine whether strains were significantly more resistant or susceptible to more toxicants or toxicant classes by chance, we conducted 1000 Fisher exact tests using the *fisher.test()* function with 2000 Monte Carlo simulations.

**Figure 3:**
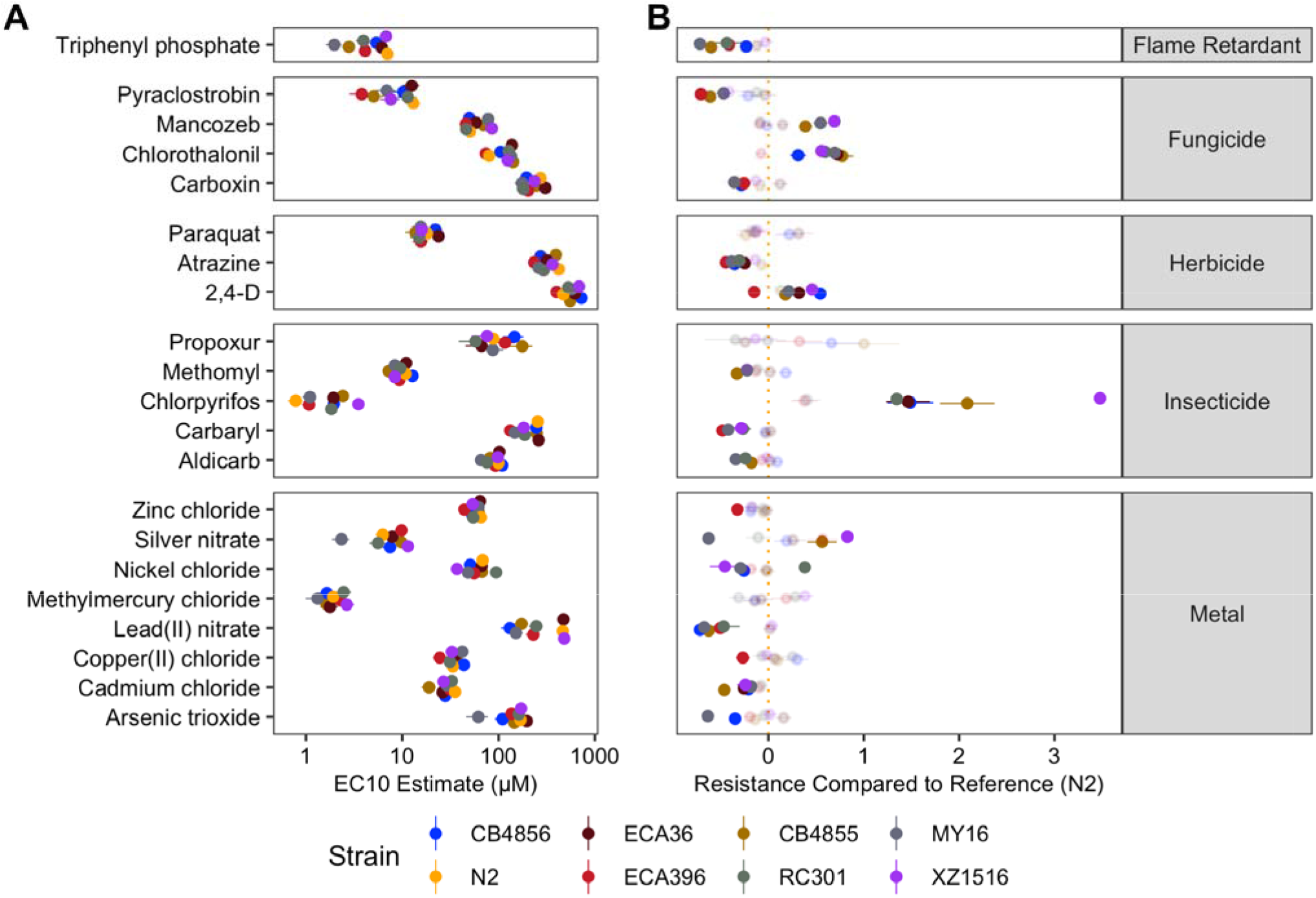
Variation in EC10 estimates can be explained by genetic differences among strains. A) Strain-specific EC10 estimates for each toxicant are displayed for each strain. Standard errors for each strain- and toxicant-specific EC10 estimate are indicated by the line extending from each point. B) For each toxicant, the relative potency of that toxicant against each strain compared to the N2 strain is shown. Solid points denote strains with significantly different relative resistance to that toxicant (Student’s t-test and subsequent Bonferroni correction with a *p_adj_* < 0.05), and faded points denote strains not significantly different than the N2 strain. The broad category to which each toxicant belongs is denoted by the strip label for each facet.

### Broad-sense and narrow-sense heritability calculations

We estimated the broad-sense heritability (*H^2^*) using the *lme4* (v1.1.27.1) R package to fit a linear mixed-effects model to the normalized phenotype data with strain as a random effect. We then extracted the among strain variance (*V_G_*) and the residual variance (*V_E_*) from the model and calculated *H^2^* with the equation *H^2^ = V_G_ / (V_G_+V_E_)*. Genetic variance (*V_G_*) can be partitioned into additive (*V_A_*) and non-additive (*V_NA_*) variance components. Narrow-sense heritability (*h^2^*) is defined as the ratio of additive genetic variance over the total phenotypic variance (*V_P_*), *i.e.*, *h^2^ = V_A_ / V_P_*. We generated a genotype matrix using the *genomatrix* profile of NemaScan, a GWAS analysis pipeline(Widmayer et al., 2022), using the variant call format (VCF) file generated in the latest CeNDR release (https://www.elegansvariation.org/data/release/latest). We then calculated *h^2^* using the *sommer* (v4.1.5) R package by calculating the variance-covariance matrix (*M_A_*) from this genotype matrix using the *sommer::A.mat* function. We estimated *V_A_* using the linear mixed-effects model function *sommer::mmer* with strain as a random effect and *M_A_* as the covariance matrix. We then estimated *h^2^* and its standard error using the *sommer::vpredict* function.

### Data availability

All code and data used to replicate the data analysis and figures presented are available for download at https://github.com/AndersenLab/toxin_dose_responses.

## RESULTS

We performed dose-response assessments using a microscopy-based high-throughput phenotyping assay (Figure 1) for developmental delay in response to 25 toxicants belonging to five major chemical classes: heavy metals (9), insecticides (8), herbicides (3), fungicides (4), flame retardants (1). Dose-response assessments for each compound were conducted using eight *C. elegans* strains representative of the genetic variation present across the species. We first quantified the population-wide lowest observed adverse effect level (LOAEL) for each compound (Supplemental Table 2). We then cleaned and normalized phenotype data in order to censor measurements obtained at problematic concentrations of various compounds and harmonized phenotypic responses across technical replicates (*see Methods*). Out of the 25 toxicants, twelve toxicants elicited variable LOAELs among the panel of strains: the insecticides aldicarb, chlorfenapyr, carbaryl, chlorpyrifos, and malathion; the fungicides pyraclostrobin and chlorothalonil; the metals manganese(II) chloride, methylmercury chloride, nickel chloride, and silver nitrate; and the flame retardant triphenyl phosphate (one-way ANOVA, Tukey HSD; *p_adj_* < 0.05).

We next estimated dose-response curves for each compound in order to more precisely describe the contributions of genetic variation to different dynamics of susceptibility among strains (Figure 1). To accomplish this step, we modeled four-parameter log-logistic dose-response curves for each compound using normalized median animal length as the phenotypic response. The slope (*b)* and effective concentration (*e*) parameters of each dose-response model were estimated using strain as a covariate, allowing us to extract strain-specific dose-response parameters. Undefined EC10 estimates (estimates greater than the maximum dose to which animals were exposed) were observed for at least one strain from two compounds (chlorfenapyr and manganese(II) chloride). Additionally, we observed virtually uniform responses and high within-strain phenotypic variance across the dose curves of deltamethrin and malathion across all strains. We speculate that this high variance is in part driven by insoluble particles in culture wells that interfered with reliable inference of animal lengths and have consequently excluded these four compounds from further dose-response analyses (Supplemental Figure 4).

Dose-response models using strain as a covariate explained significantly more variation than those models without the strain covariate for the other 21 compounds (F*-*test; *p <* 0.001). We observed substantial variation in effective concentration between toxicants within classes of compounds known to have similar modes of toxicity (Two-way ANOVA; *p* < 0.001) but not across strains (Two-way ANOVA; *p* ≥ 0.163) (Figure 3A). All fungicides and herbicides exhibited significantly different EC10 estimates (two-way ANOVA, Tukey HSD; *p_adj_* ≤ 0.003). EC10 estimates for propoxur were not significantly different from aldicarb, nor were the estimates for methomyl compared to chlorpyrifos (two-way ANOVA, Tukey HSD; *p_adj_* ≥ 0.934) but EC10 estimates for all other compounds within the insecticide class were significantly different (two-way ANOVA, Tukey HSD; *p_adj_* ≤ 0.001). EC10 estimates for lead(II) nitrate were significantly different from all other tested metals (two-way ANOVA, Tukey HSD; *p_adj_* < 0.001). EC10 estimates for arsenic trioxide were significantly different from all tested metals (two-way ANOVA, Tukey HSD; *p_adj_* ≤ 0.050), except nickel chloride (two-way ANOVA, Tukey HSD; *p_adj_* = 0.068). EC10 estimates for all other metals were not significantly different from each other (two-way ANOVA, Tukey HSD; *p_adj_* ≥ 0.392). These results suggest that susceptibility to different toxicants in *C. elegans* is not explained by differences in the mode of action of each toxicant.

Most differences in EC10 were explained by differences among compounds of different classes. However, variation in EC10 estimates caused by genetic differences among strains were pervasive (Figure 3B). In order to quantify these differences, we calculated the relative potency of each compound in pairwise comparisons among all strains (Supplemental Table 3). To contextualize these differences, we filtered down to comparisons between the reference strain N2 and all others and subsequently calculated the difference in potency with respect to the laboratory reference strain. In total, we observed 66 instances across 18 compounds where at least one strain was significantly more resistant or sensitive than the reference strain N2 using EC10 as a proxy (Student’s t-test, Bonferroni correction; *p_adj_* < 0.05) with paraquat and propoxur being the exceptions (Figure 3B). Twenty-two strain comparisons showed greater resistance than responses in the N2 strain, and 44 strain comparisons showed greater susceptibility across all compounds. Relative resistance was more generalized across strains, with four different strains exhibiting significant sensitivity to at least three toxicants with respect to the N2 strain. Of the instances in which a strain was significantly more sensitive than the N2 strain, 47.8% of the cases were either the ECA396 or MY16 strains, which were the two strains with the greatest number of compounds that elicited sensitivity. Furthermore, the observed frequency of strains with significantly greater toxicant sensitivity with respect to the N2 strain was significantly different than expected under the null (*see Methods*; Fisher’s exact test; *p* < 0.05), suggesting that diverse *C. elegans* strains are not equally likely to be susceptible or resistant with respect to the commonly used reference strain N2.

Strain-specific slope (*b*) estimates for each dose-response model varied substantially as well but followed different patterns than those estimates observed for EC10 (Figure 4A). We again observed substantial variation in slope estimates between toxicants within classes of compounds known to have similar modes of toxicity (two-way ANOVA; *p* < 0.001) but not across strains (two-way ANOVA; *p* ≥ 0.074). Slope estimates for pyraclostrobin were significantly lower than all other fungicides (two-way ANOVA, Tukey HSD; *p_adj_* ≤ 0.0002). Slope estimates for 2,4-D were significantly lower than those estimates for the other two herbicides (two-way ANOVA, Tukey HSD; *p_adj_*< 0.0001). Among insecticides, the only slope estimates that were not significantly different from each other were methomyl and aldicarb (two-way ANOVA, Tukey HSD; *p_adj_* = 0.999). Slope estimates for nickel chloride were significantly different from all other metals (two-way ANOVA, Tukey HSD; *p_adj_* ≤ 0.031).

**Figure 4:**
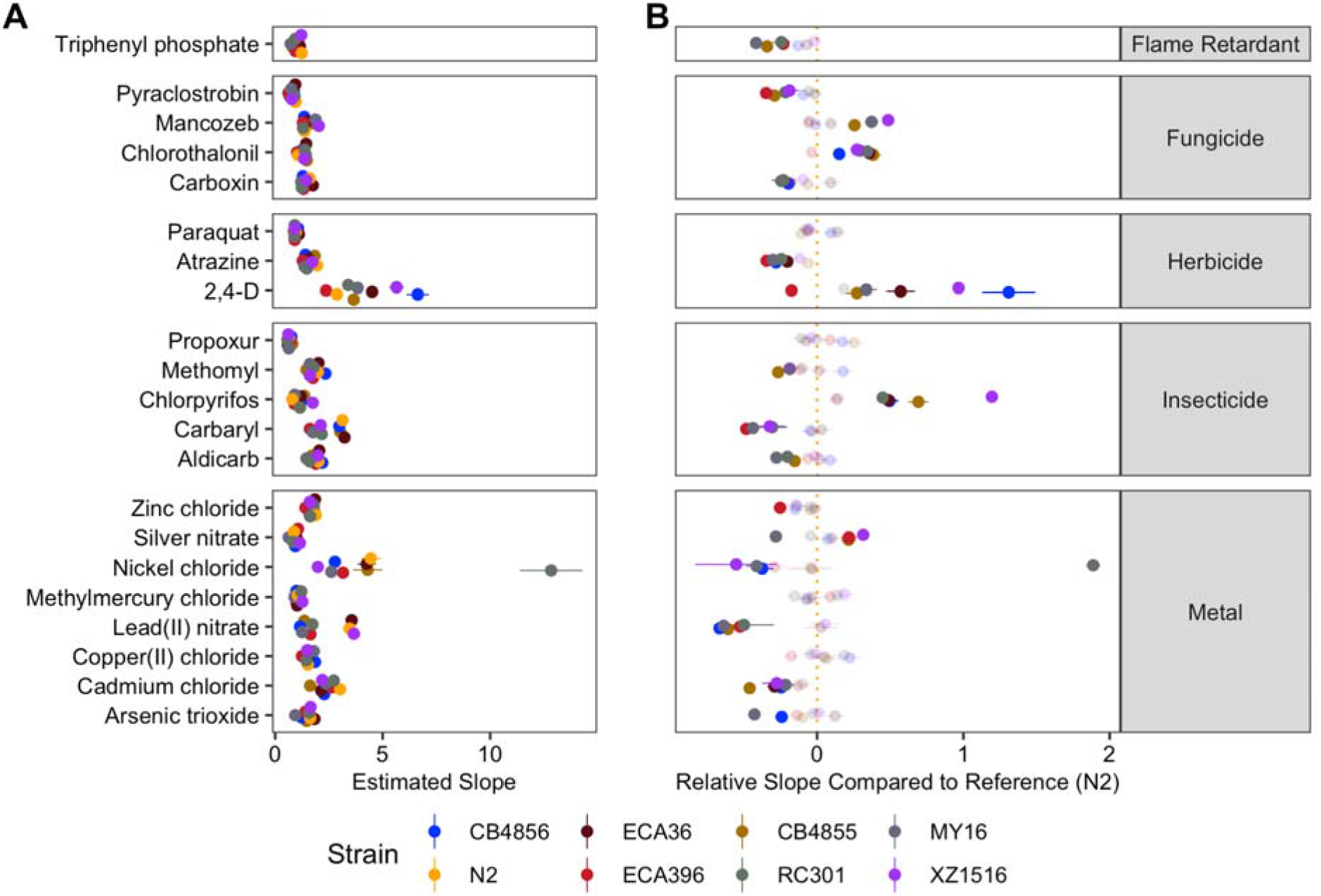
Variation in dose-response slope estimates can be explained by genetic differences among strains. A) Strain-specific slope estimates for each toxicant are displayed for each strain. Standard errors for each strain- and toxicant-specific slope estimate are indicated by the line extending from each point. B) For each toxicant, the relative steepness of the dose-response slope inferred for that strain compared to the N2 strain is shown. Solid points denote strains with significantly different dose-response slopes (Student’s t-test and subsequent Bonferroni correction with a *p_adj_* < 0.05), and faded points denote strains without significantly different slopes than the N2 strain. The broad category to which each toxicant belongs is denoted by the strip label for each facet.

We next compared the relative steepness of dose-response slope estimates compared to the N2 reference strain, analogously to our EC10 relative potency analysis (all strain-by-strain comparisons can be found in Supplemental Table 4) and observed 76 significantly different slope steepness comparisons with the reference strain (Figure 4B). The greatest number of significantly different slope estimates among strains were observed in insecticides, which comprised 24 (31%) of the comparisons. Four strains exhibited at least ten significantly different slope estimates (CB4855, CB4856, MY16, XZ1516), and five strains (CB4855, CB4856, ECA396, MY16, RC301) exhibited more instances of significantly shallower dose-response slopes than N2. Furthermore, the number of significantly shallower dose-response slopes for each strain compared to the N2 strain was significantly different from that expected under the null (*see Methods*; Fisher’s exact test; *p* = 0.041).

Taken together, these results suggest that genetic differences between *C. elegans* strains mediate differential susceptibility and toxicodynamics across a diverse range of toxicants. In order to quantify the degree of phenotypic variation attributable to segregating genetic differences among strains, we first estimated the broad-sense heritability of the phenotypic response for each dose of every compound. We observed a wide spectrum of broad-sense and narrow-sense heritability estimates across compounds and dose ranges (Figure 5). Excluding control doses, the average broad-sense heritability across all doses of each compound ranged from 0.05 (atrazine) to 0.36 (chlorpyrifos), and narrow-sense heritability ranged from 0.05 (copper(II) chloride) to 0.37 (chlorpyrifos). Motivated by the wide range of additive genetic variance estimates that we observed across doses of each compound, we asked how closely the doses that exhibited the greatest narrow-sense heritability aligned with EC10s estimated for each compound. We compared the narrow-sense heritabilities between the dose closest to the estimated EC10 and the doses that exhibited the maximum narrow-sense heritability for each of the 21 compounds with definitive EC10 estimates. We observed a strong relationship between the doses that approximate the EC10 for each compound and the doses that yielded the greatest narrow-sense heritability (Figure 6). Interestingly, although the correlation between these two endpoints was strong, the dosage of each compound that exhibited the greatest additive genetic variance was always greater than the dose that approximated the EC10 for that compound, demonstrating that the additive genetic variation responsible for the greatest differences in toxicant responses among *C. elegans* strains is typically revealed at greater exposure levels than the average estimated EC10.

**Figure 5:**
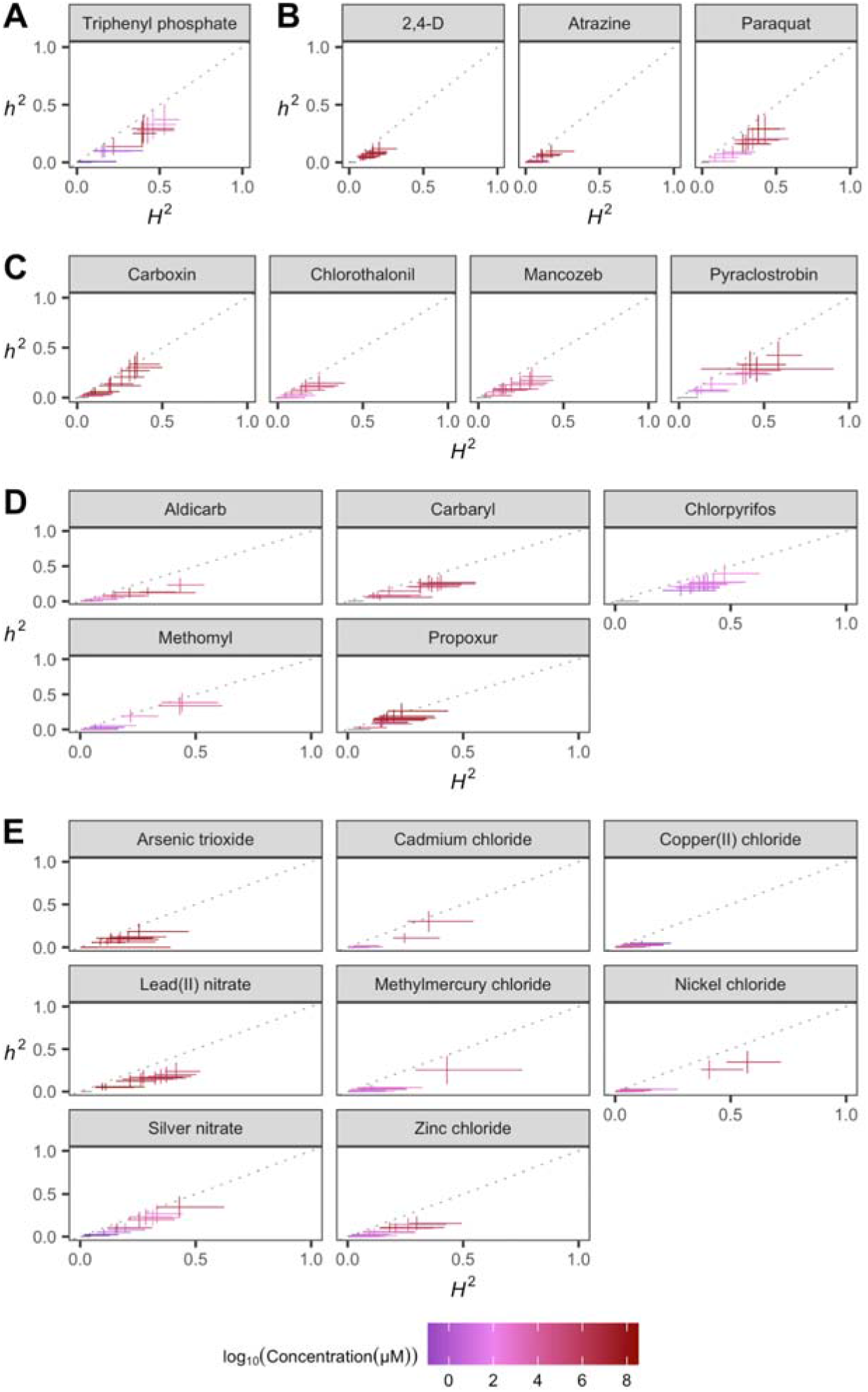
Variation in toxicant responses is heritable among genetically diverse *C. elegans* strains. The broad-sense (x-axis) and narrow-sense heritability (y-axis) of normalized animal length measurements was calculated for each concentration of each toxicant (*Methods; Broad-sense and narrow-sense heritability calculations*). The color of each cross corresponds to the log-transformed dose for which those calculations were performed. The horizontal line of the cross corresponds to the confidence interval of the broad-sense heritability estimate obtained by bootstrapping, and the vertical line of the cross corresponds to the standard error of the narrow-sense heritability estimate.

**Figure 6:**
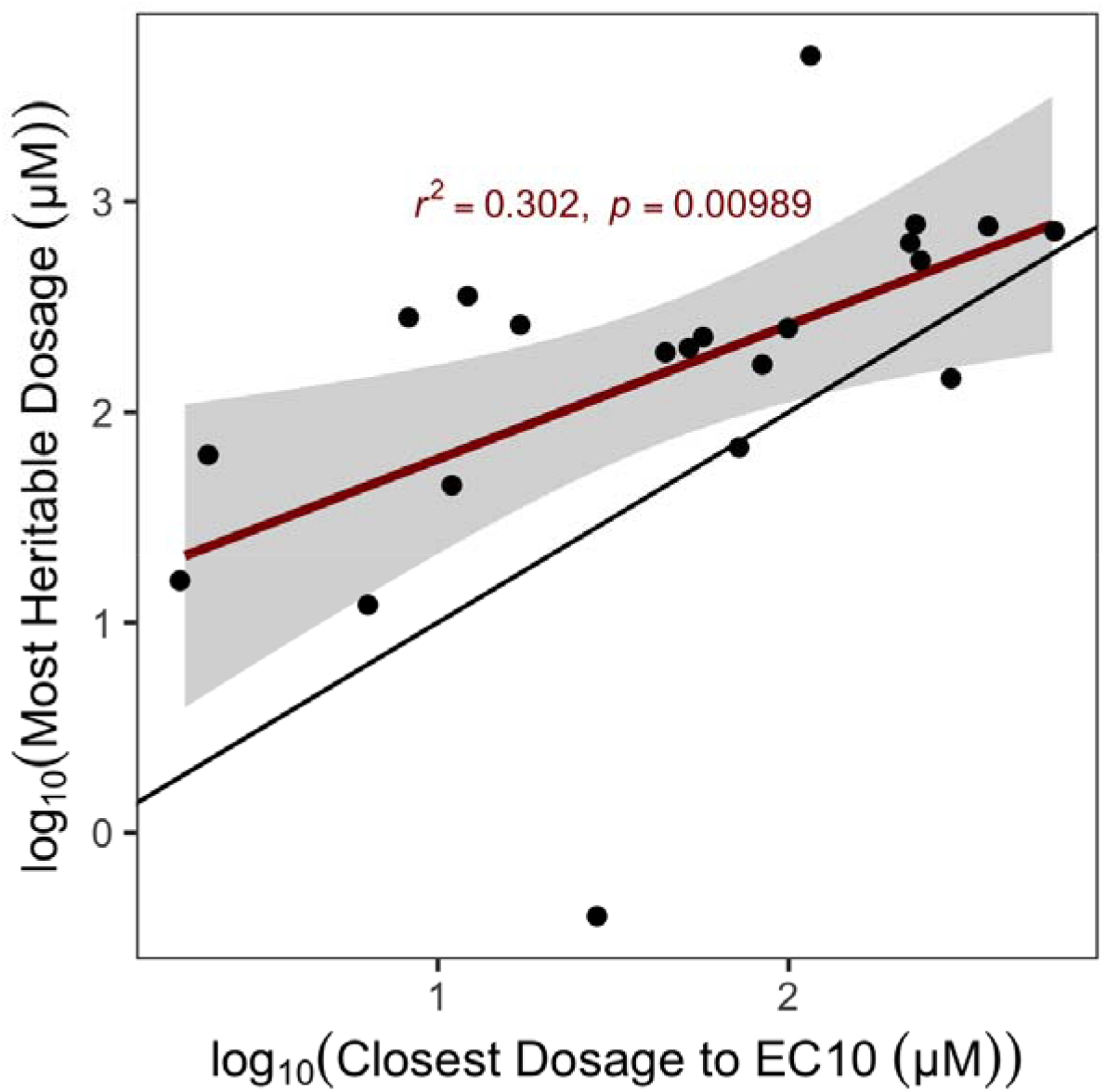
EC10 estimates from genetically diverse individuals predict doses eliciting heritable responses. The log-transformed dose that elicited the most heritable response to each toxicant (y-axis) is plotted against the log-transformed dose of that same toxicant nearest to the inferred EC10 from the dose-response assessment. The dose closest to the EC10 across all toxicants exhibited significant explanatory power to determine the dose that elicited heritable phenotypic variation.

## DISCUSSION

One of the central goals of toxicology is to achieve precise chemical risk assessments in populations characterized by diversity over broad socioeconomic, environmental, and genetic scales. At the level of initial screening in model organisms, these assessments have typically been limited to a single strain or cell line’s genetic background. However, given the sheer amount of uncharacterized toxicants being produced, it is economically unfeasible to rely entirely on mammalian systems to rigorously evaluate these hazards on a reasonable time scale. Research using *C. elegans* as a model is a staple of toxicology, particularly when it comes to identifying key regulators of cellular responses to heavy metal and pesticide exposures (Hartman et al., 2021; Hunt, 2017). However, these discoveries have typically relied on perturbing a single genome (and therefore a singular collection of “wild-type” alleles) using RNA interference or knockout alleles for individual genes. In this study, we expanded the scope of *C. elegans*-based chemical hazard evaluations to consider the effects of naturally occurring genetic variants in the *C. elegans* species by performing dose-response analysis using the PD1074 laboratory-adapted reference strain as well as seven wild strains representing the major axes of species-wide genetic variation. We conducted these analyses using a high-throughput microscopy assay that facilitates rigorous control over experimental noise, genetic effects, and toxic exposure across millions of *C. elegans* individuals from each of our eight genetic backgrounds. This paradigm allowed us to precisely estimate the effects of genetics on impaired development in the presence of a toxicant and tease them apart from experimental noise. Estimating toxic endpoints of chemical hazards has been previously executed using high-throughput screening of *C. elegans* responses (Boyd et al., 2012; Evans et al., 2018). In our study, we have leveraged and expanded on these types of platforms by explicitly estimating genetic effects on dose-response parameters.

One goal of dose-response analysis is to identify a point of departure (POD) for exposure to a certain compound (*e.g.*, a dosage at which a population begins to respond adversely to a hazard) based on empirical data. We demonstrated that EC10 estimates and slope parameters vary significantly between genetically distinct *C. elegans* strains and that, in fact, the N2 reference strain exhibits a significantly different dose-response profile than at least one other strain with respect to every toxicant we assessed. Additionally, strain-agnostic EC10 estimates are correlated with, but generally lower than, the dose at which we observed the largest additive genetic variance. These observations suggest that previous analyses of toxicity in *C. elegans* might suffer from “genetic blindspots” in that significant intrinsic drivers of population-level toxicity are being systematically ignored, which then masks a source of complexity in toxicant susceptibility. For example, we observed that the strains ECA396 and MY16 are significantly more sensitive than other strains across more toxicants than expected by chance. The susceptibility profiles of these strains underscore the need to assess hazard risk across individuals that are intrinsically susceptible or resistant in order to understand the implications of dose-response endpoints. Because our high-throughput assay only reports the magnitude of developmental retardation over one generation as a trait, it remains unknown whether the resistance we observed in these strains, or for a given toxicant more broadly, extend to other toxicity endpoints (*e.g.*, germline mutagenesis, effects on reproduction, metabolic signatures, or neurotoxicity). The toxicants in our study belong to classes of chemicals with documented effects on all of these organ systems, so the identification of putatively resistant genetic backgrounds could represent fertile ground for the discovery of novel pathways that potentiate well characterized stress responses.

An open question in toxicogenomics is the degree to which variation in human disease and development can be explained by our chemical environment, and whether these contributions exceed those from genetic differences among individuals. Our study suggests that for any given compound, we can find a dosage for which at least 20% of the variation in developmental delay can be explained by genetic differences between *C. elegans* strains. Furthermore, we show empirical support for the notion that toxic endpoints derived in experimental studies from one genetic background cannot be neatly ported across genetically diverse individuals. These findings build upon similar analyses conducted using human cell lines derived from the 1000 Genomes Project (Abdo et al., 2015), which revealed substantial heritability of dose-response endpoints. Given that high-throughput platforms exist that facilitate these analyses, stakeholders in toxicology must prioritize the derivation of PODs derived in genetically diverse model organism populations in order to precisely account for this source of uncertainty in hazardous chemical evaluations. Also, given the high heritabilities of the compounds we tested, quantitative genetic analyses such as genome-wide association studies in genetically diverse model organisms provide an opportunity to identify conserved genetic loci that mediate population-level differences in toxicant susceptibility.

## ACKNOWLEDGEMENTS

We would like to thank members of the Andersen laboratory for helpful comments on the manuscript. This work was supported by an NIH NIEHS grant (ES029930) to E.C.A.

## SUPPLEMENTAL FIGURES

**Supplemental Figure 1.**
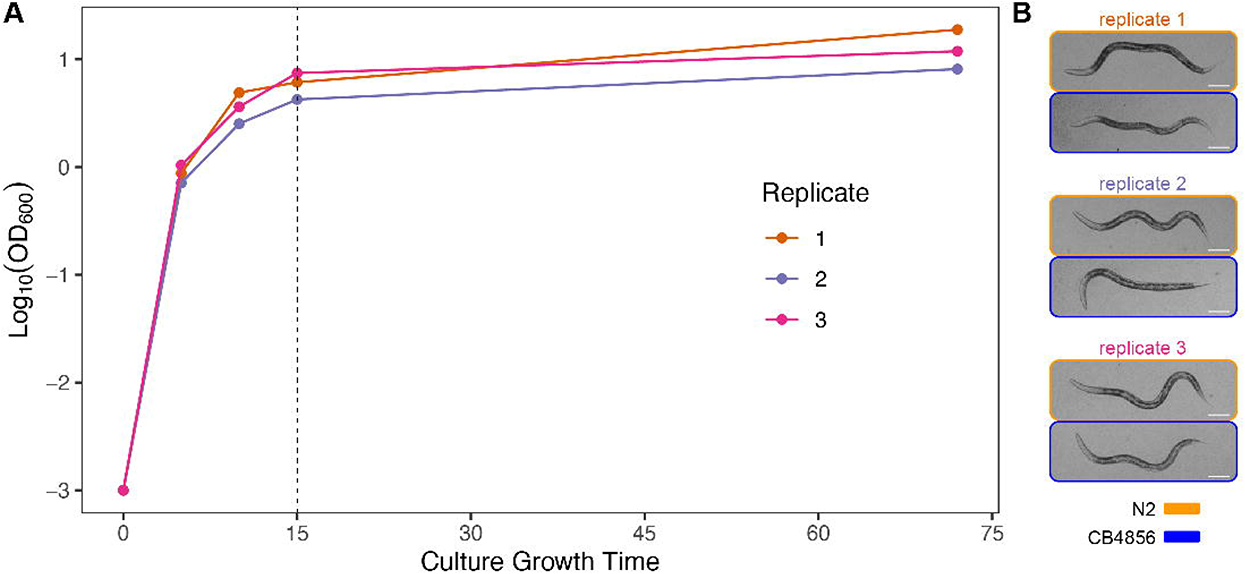
Repeatability of HB101 *E. coli* culture growth dynamics and nematode development when fed with 15 hour cultures at OD_600_ 10. Three independent growth curves of HB101 *E. coli* are shown by color (A). The replicate cultures were inoculated at an OD_600_ value of 0.001 from 18-hour starter cultures and grown at 37°C with shaking at 180 rpm. The points represent the mean of triplicate OD_600_ values for each timepoint. The 15-hour time point (vertical dashed line) was chosen for nematode food preparations because it represented the early stationary phase and reproducibly supported nematode development to the L4 stage after 48 hours of feeding from arrested L1s. Representative images of N2 (orange) and CB4856 (blue) strains that were fed with independent 15-hour *E. coli* HB101 preparations for 48 hours post L1 arrest at a final concentration of OD_600_ 10 in K medium (B). The images were cropped from a 4.2 megapixel image captured using a 2X objective, which is identical to the resolution of images that we used for our toxicant assays. The white scale bars at the lower right represent 10 µm.

**Supplemental Figure 2.**
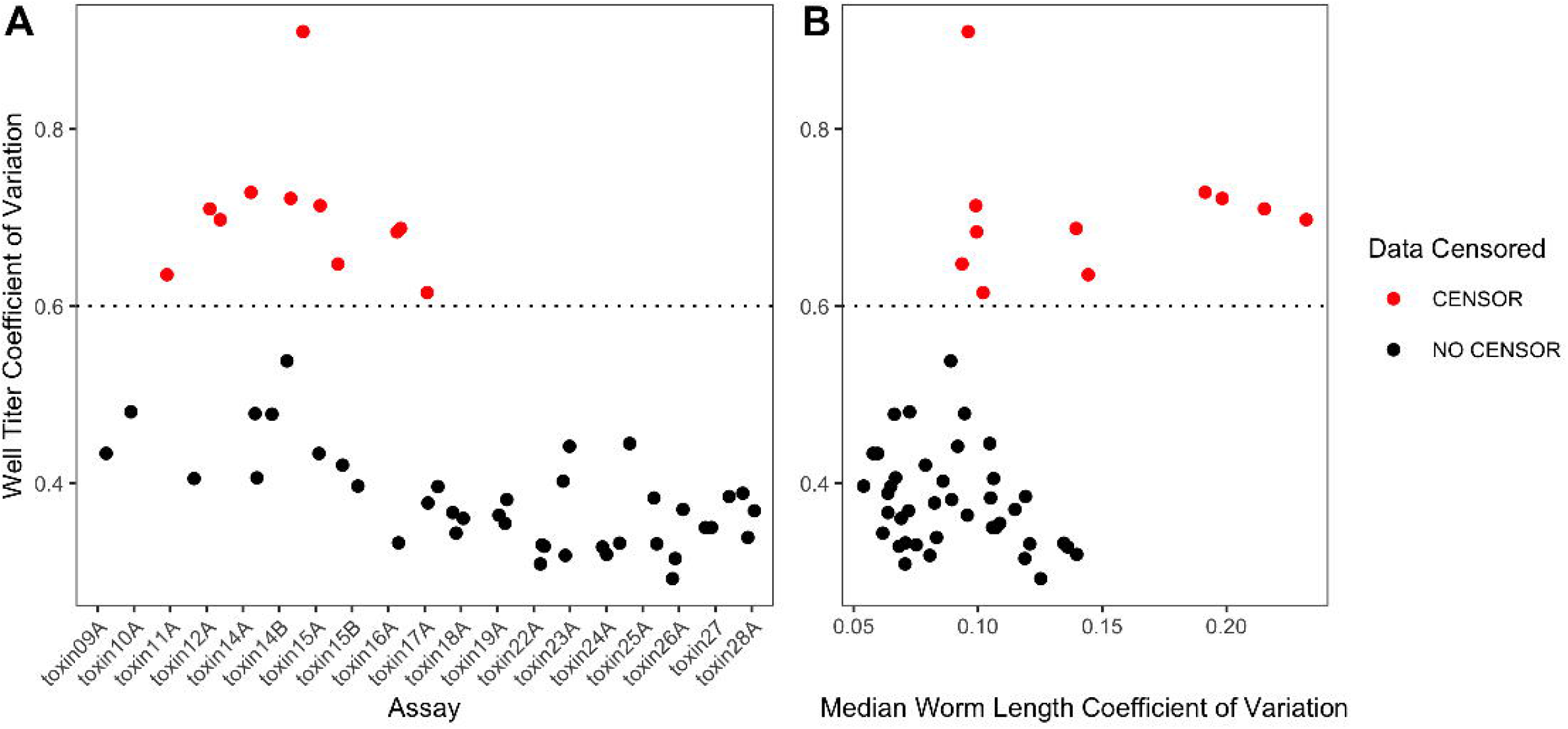
Toxicant dose-response data cleaning heuristics. Coefficients of variation in the number of animals detected in control wells (control well CV) calculated across all technical replicates, toxicants, and strains for each experimental block are shown. These metrics are shown as a function of experimental block (A) and median animal length coefficient of variation (B). Technical replicates exhibiting a control well CV greater than 0.6 (above the dashed line in red) were censored from dose-response analyses.

**Supplemental Figure 3.**
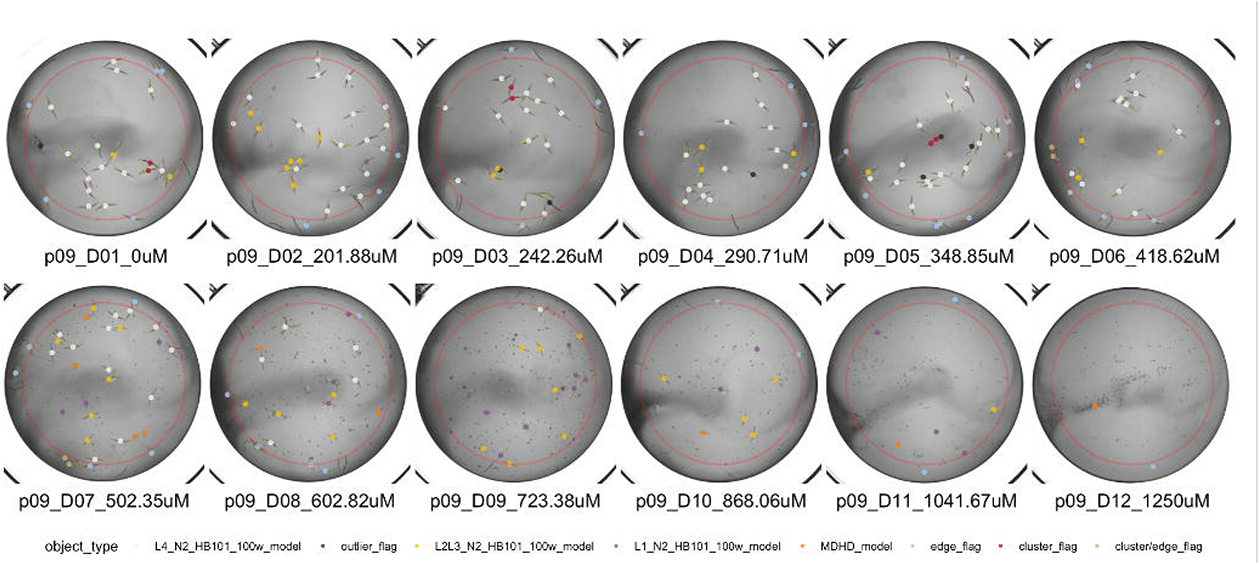
Problematic well features for downstream data analysis. Snapshots of well images obtained in arsenic trioxide dose-response experiments demonstrating problematic well features. In the top panel, most animals in each well are identified with our trained worm models indicated by colored dots. In the bottom panel, animals and particulate matter are inaccurately positively and negatively identified.

**Supplemental Figure 4.**
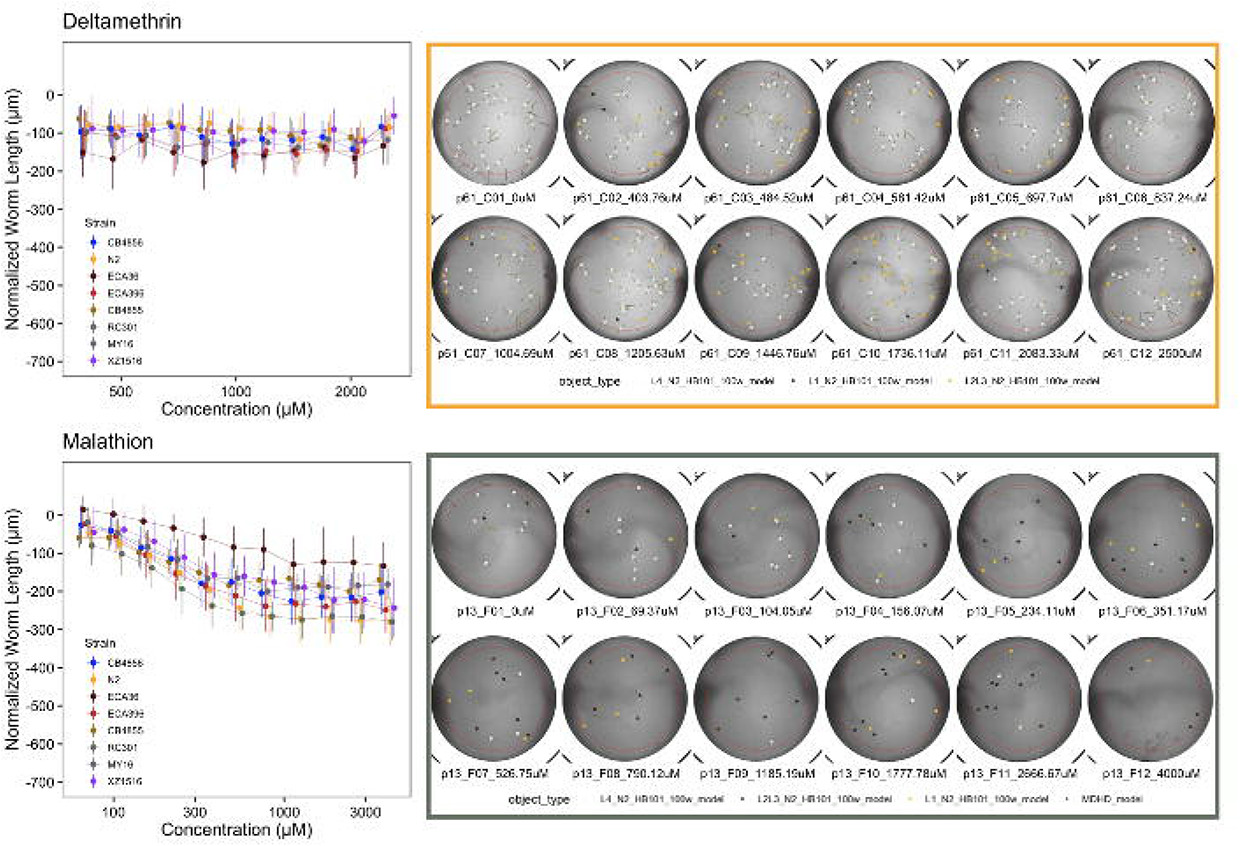
Dose-response curves and well images for deltamethrin and malathion dose-response experiments. Dose-response curves of deltamethrin and malathion with representative well snapshots. Most *C. elegans* strains exhibited negligible responses across the dose range for these exposures.

**Supplemental Figures 5.**
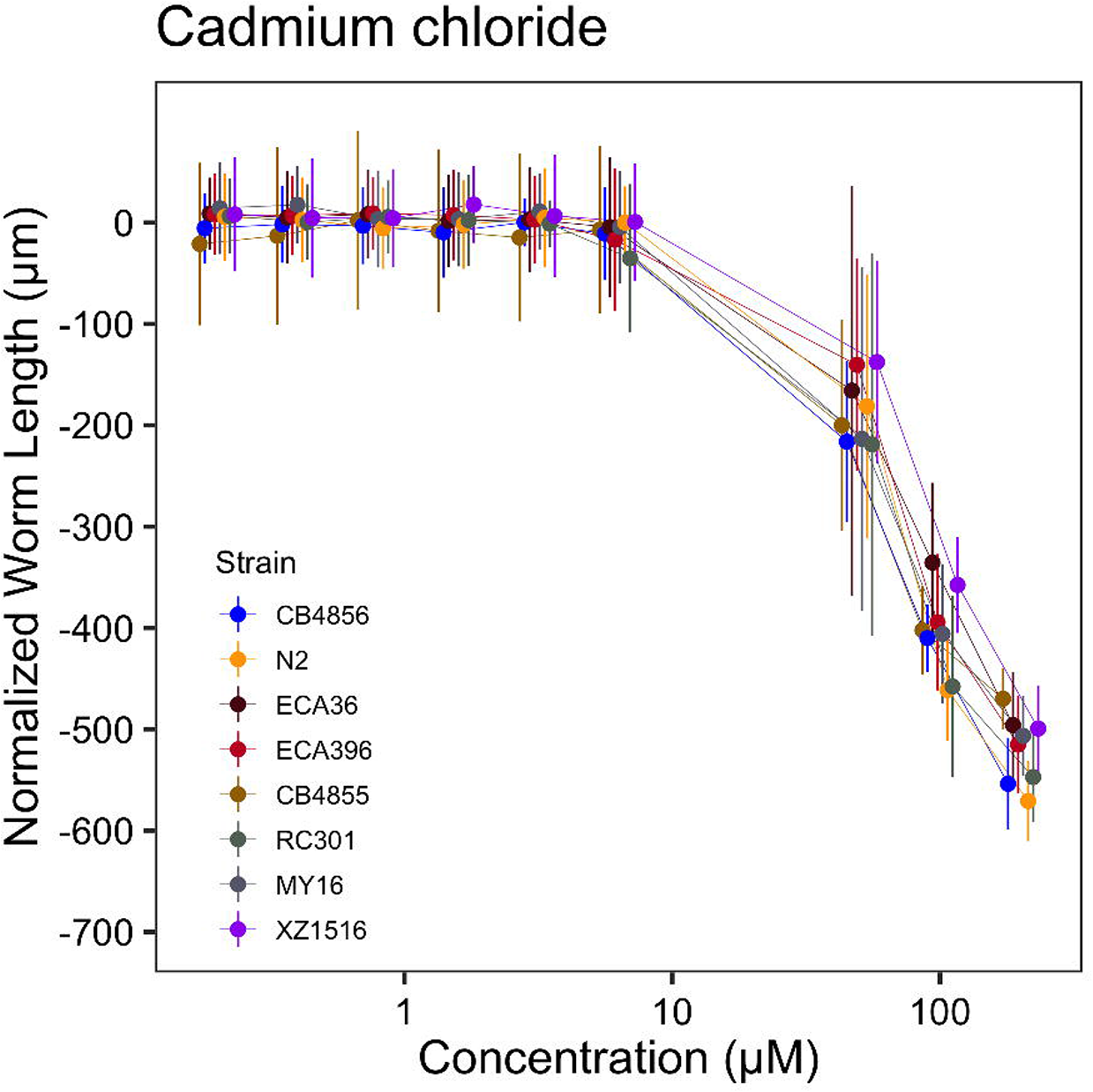
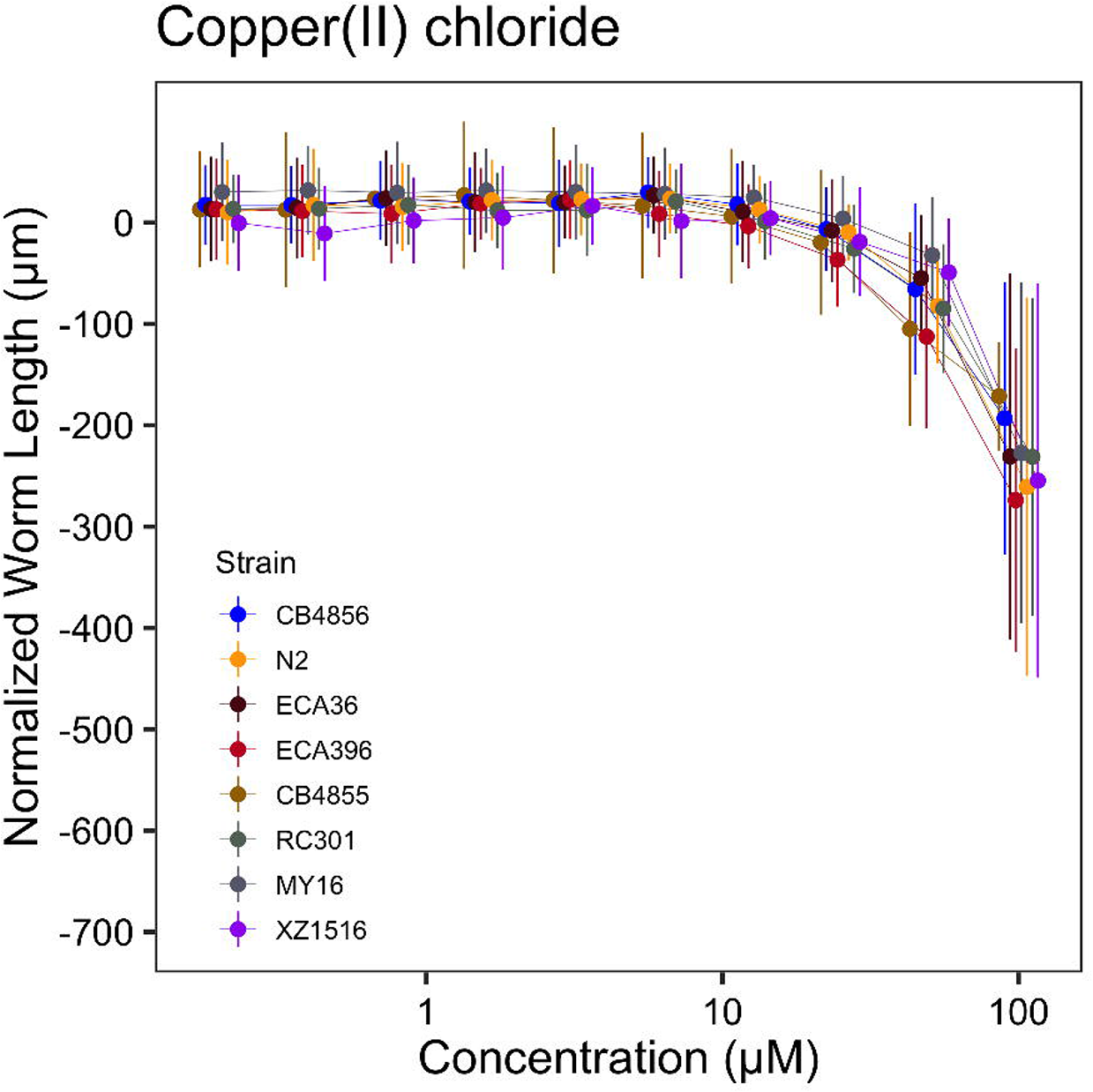
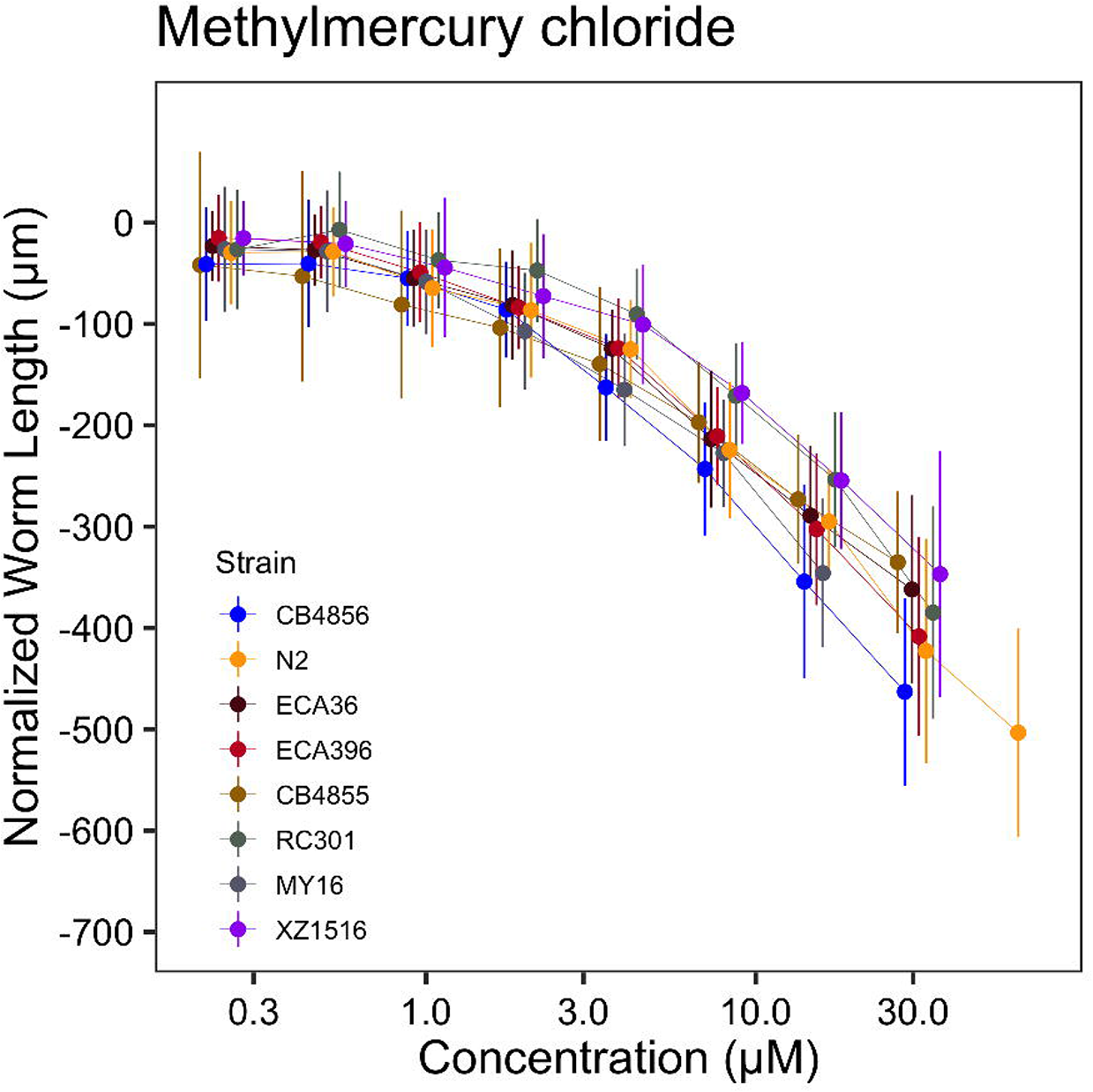
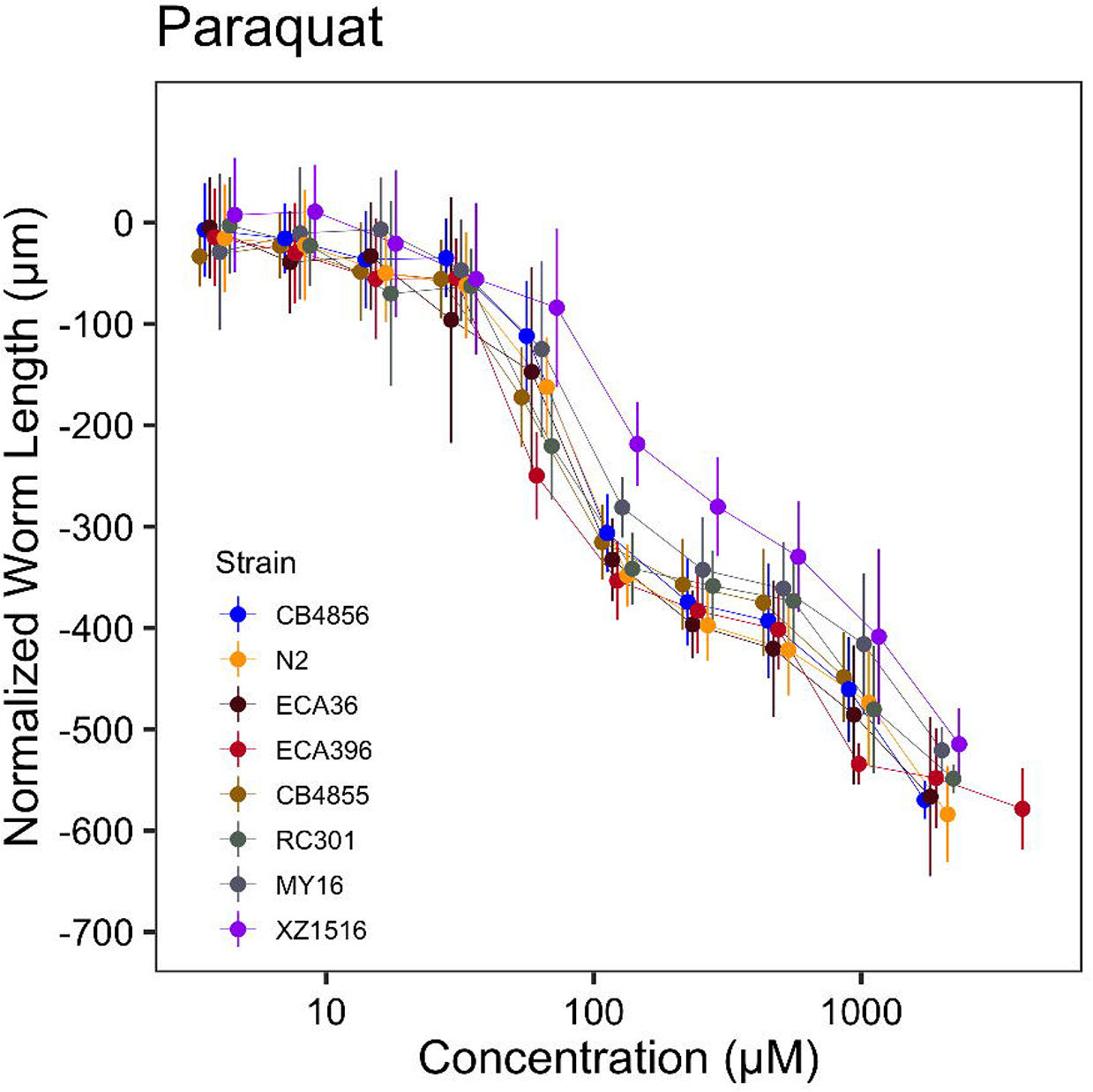
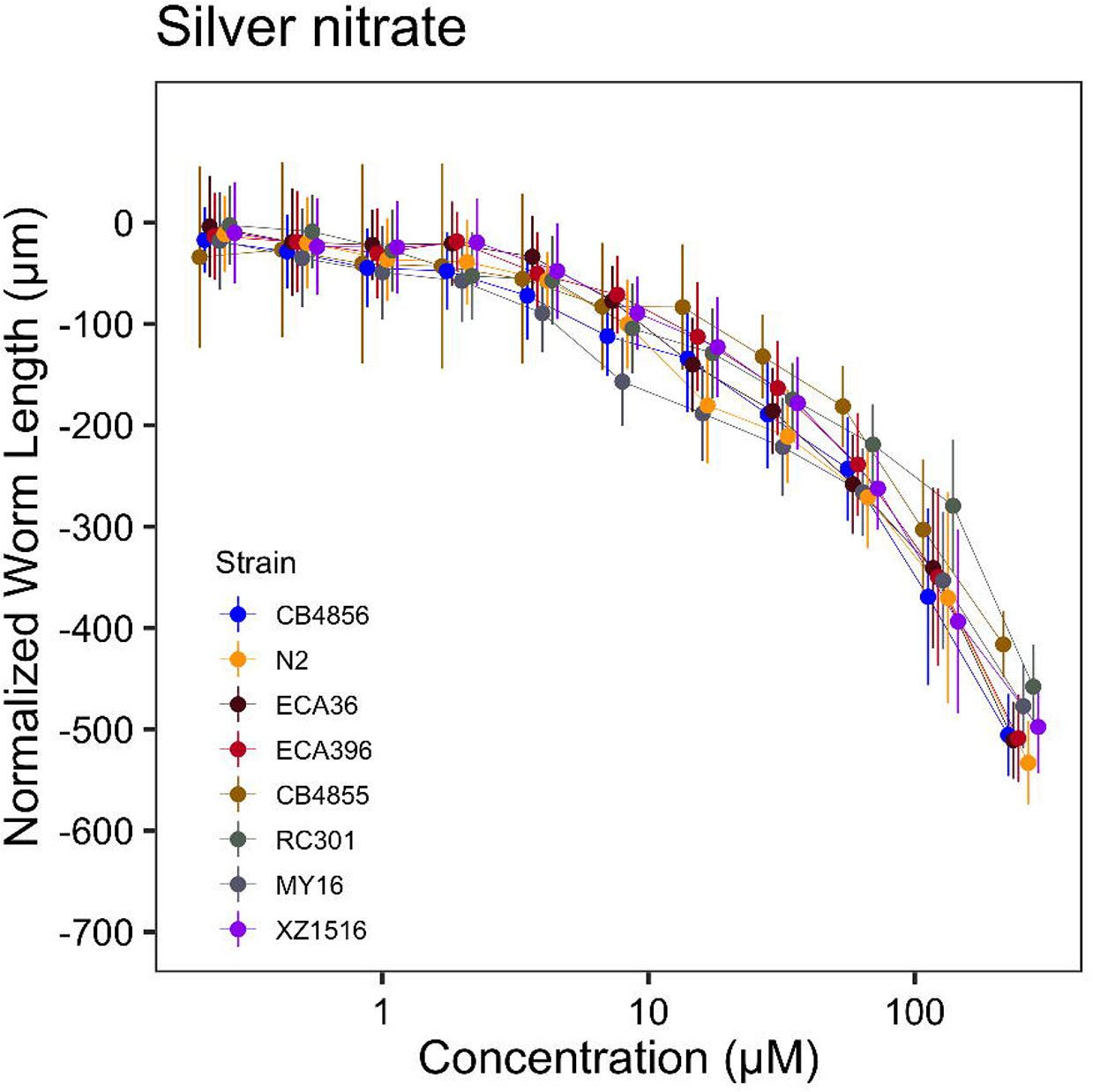
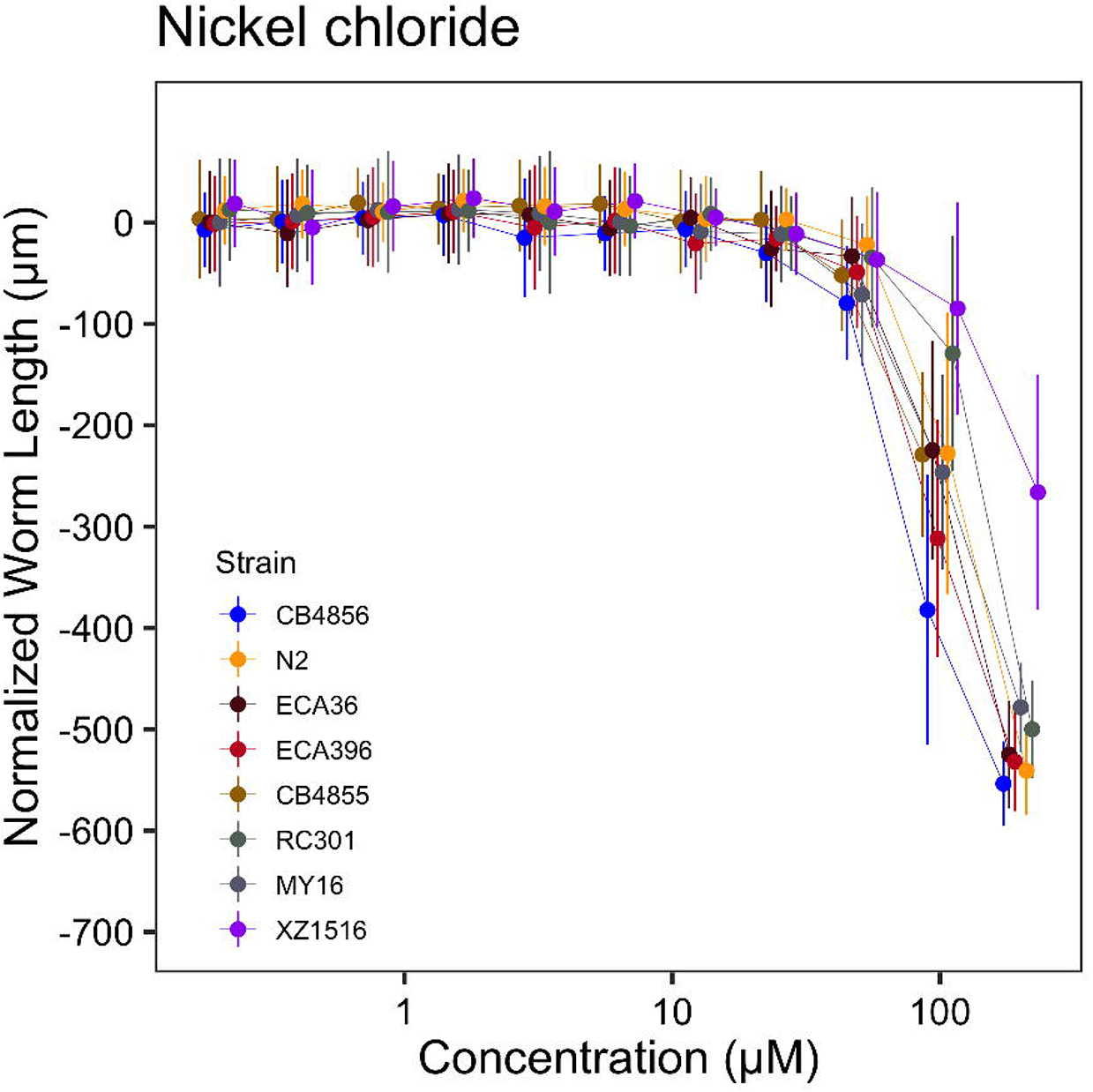
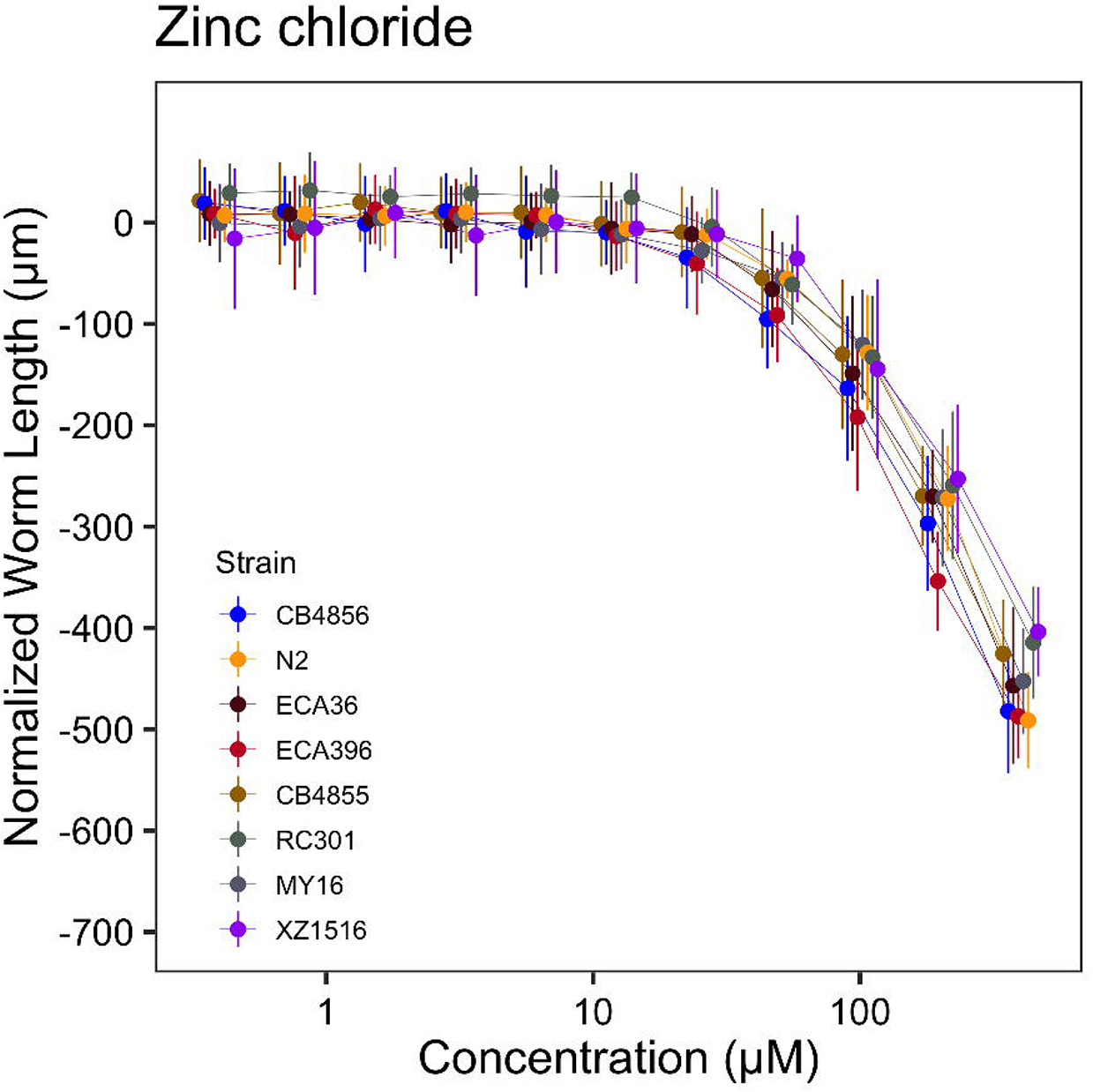
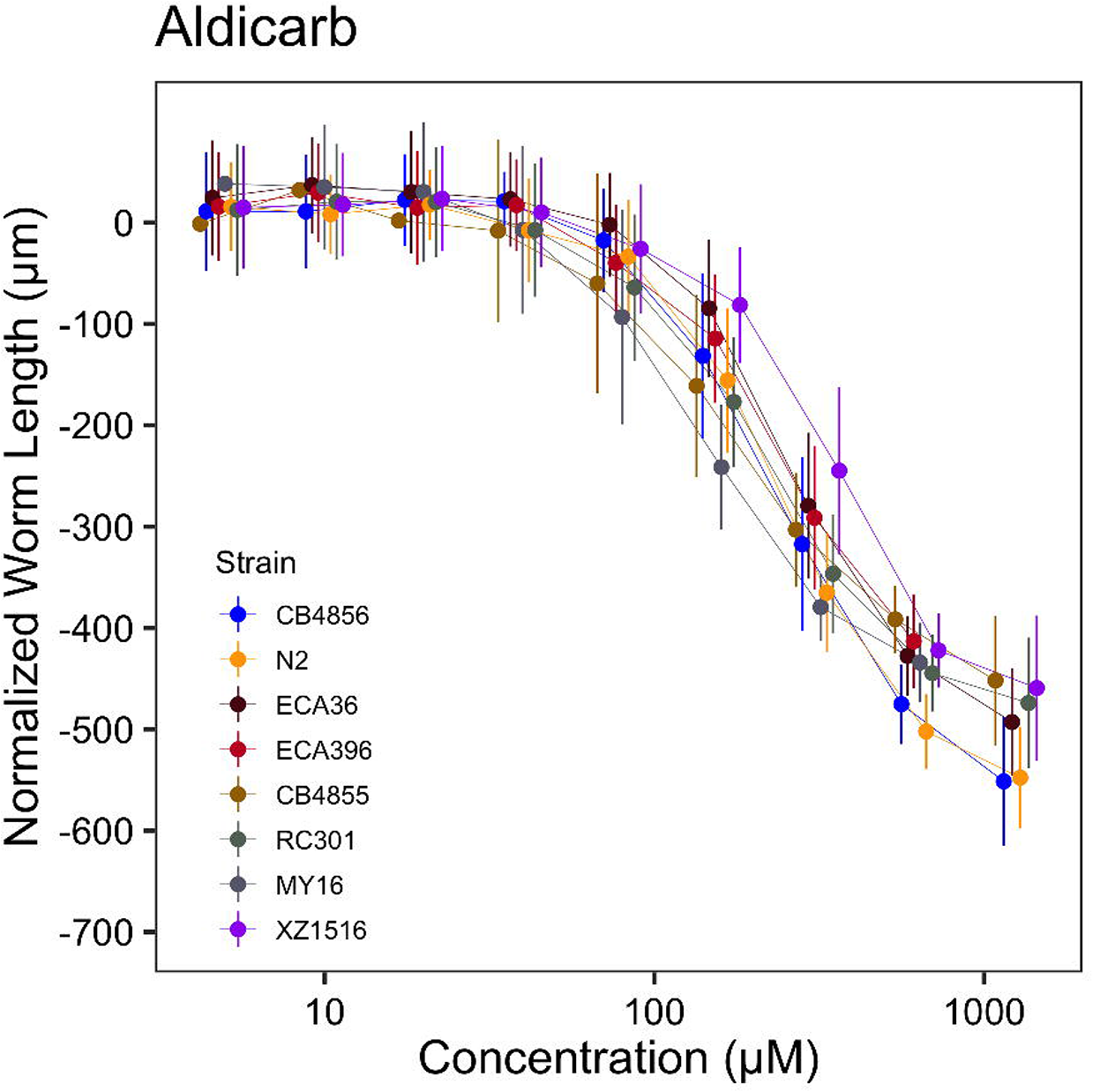
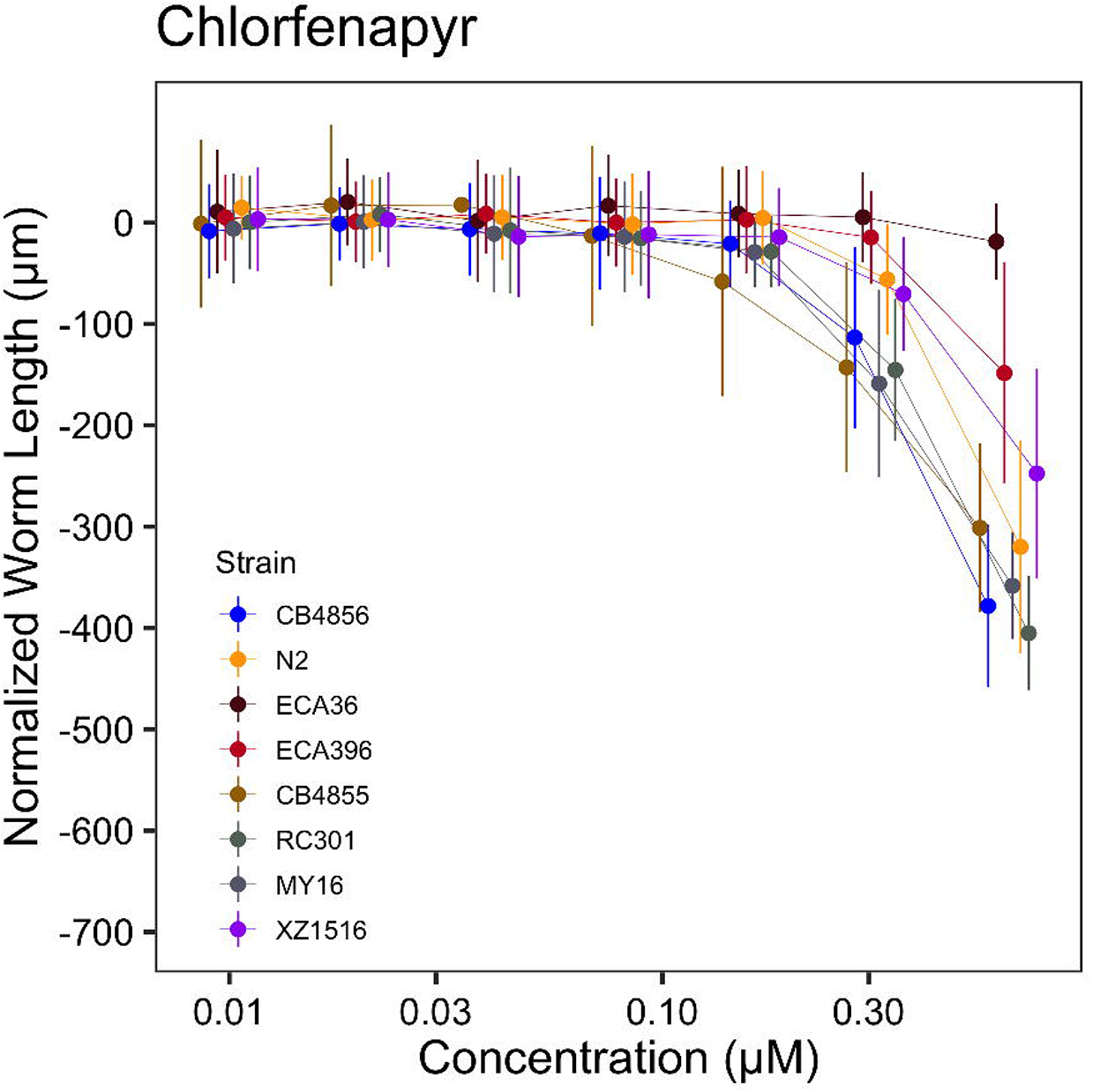
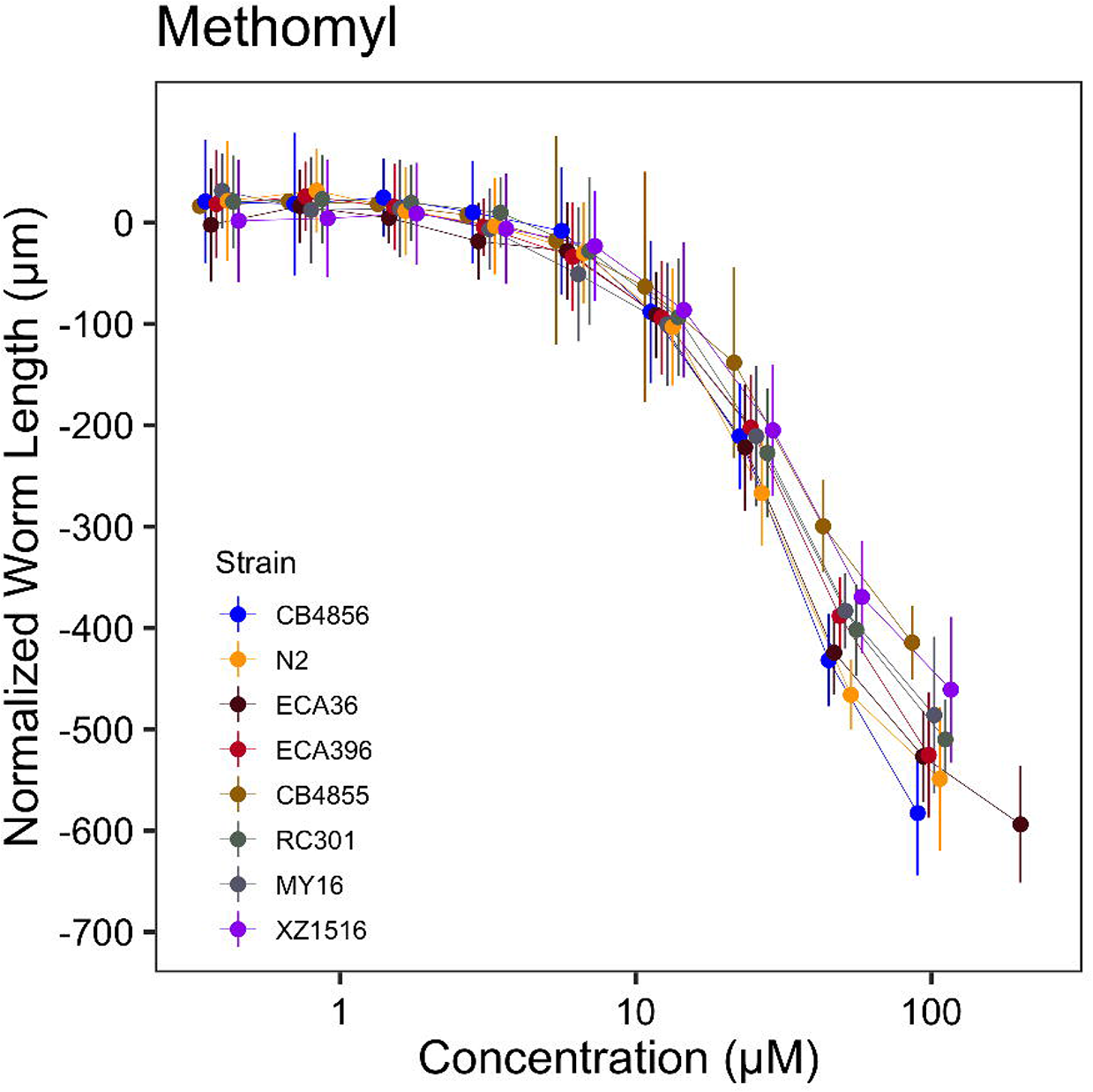
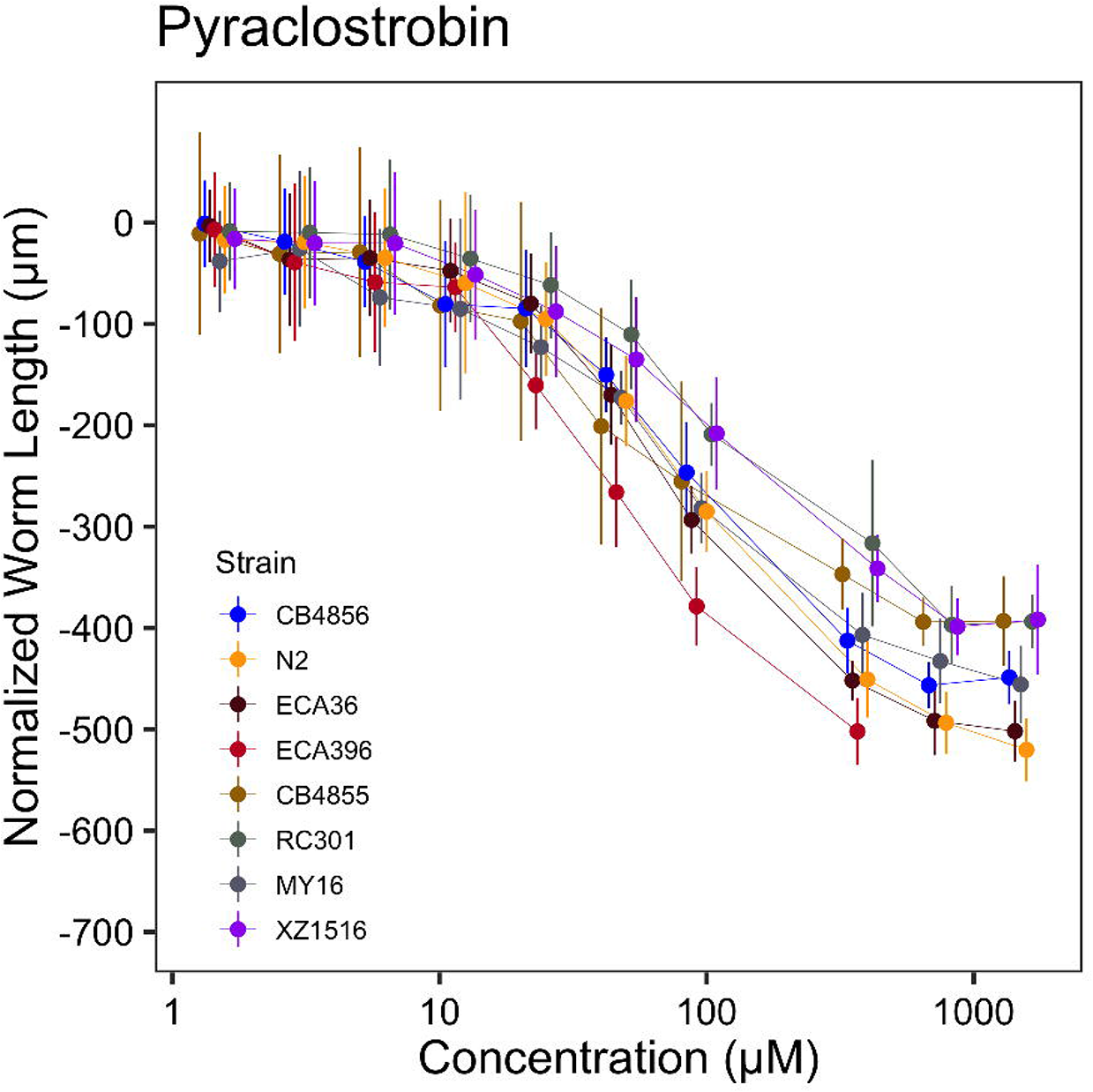
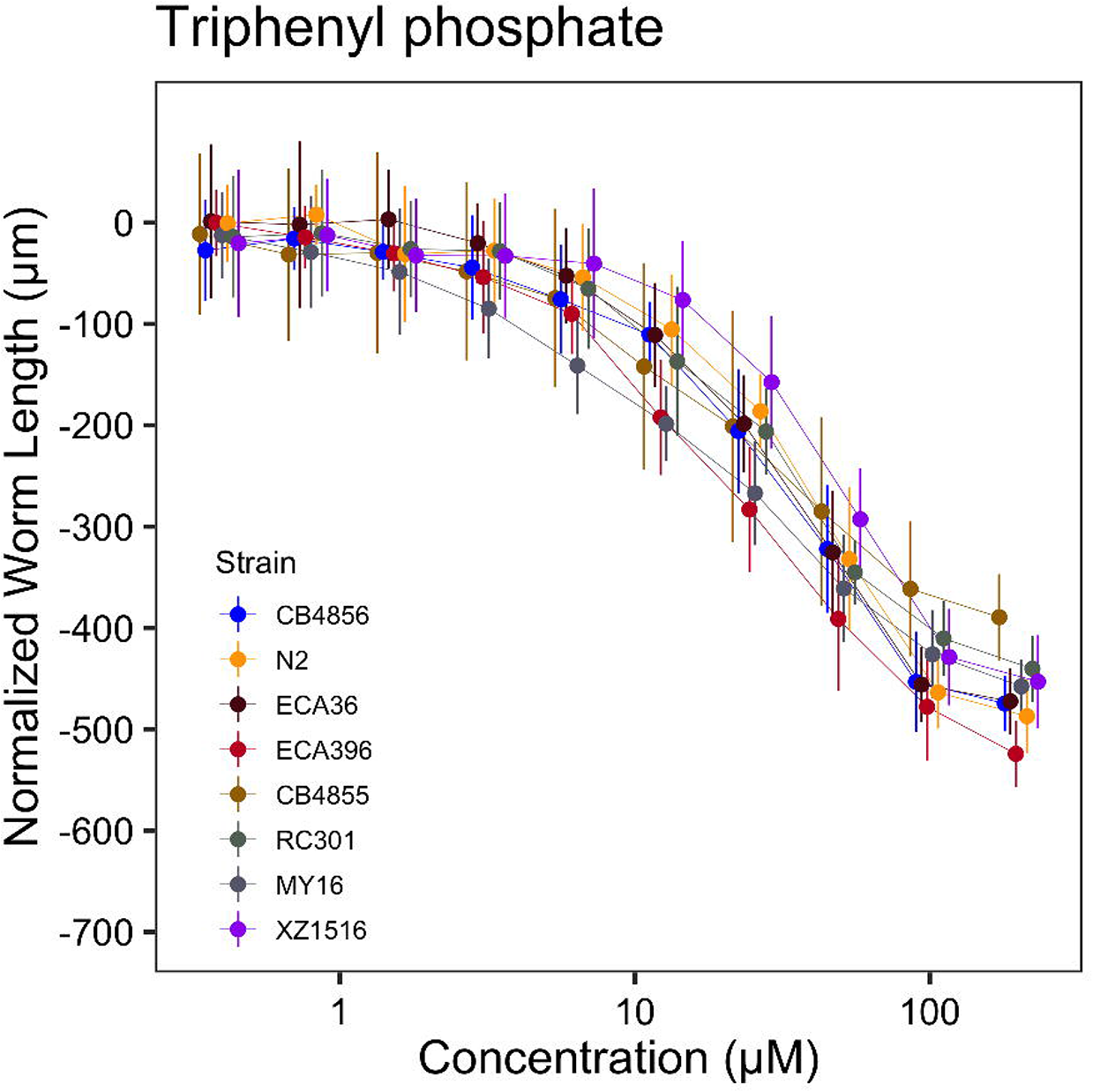
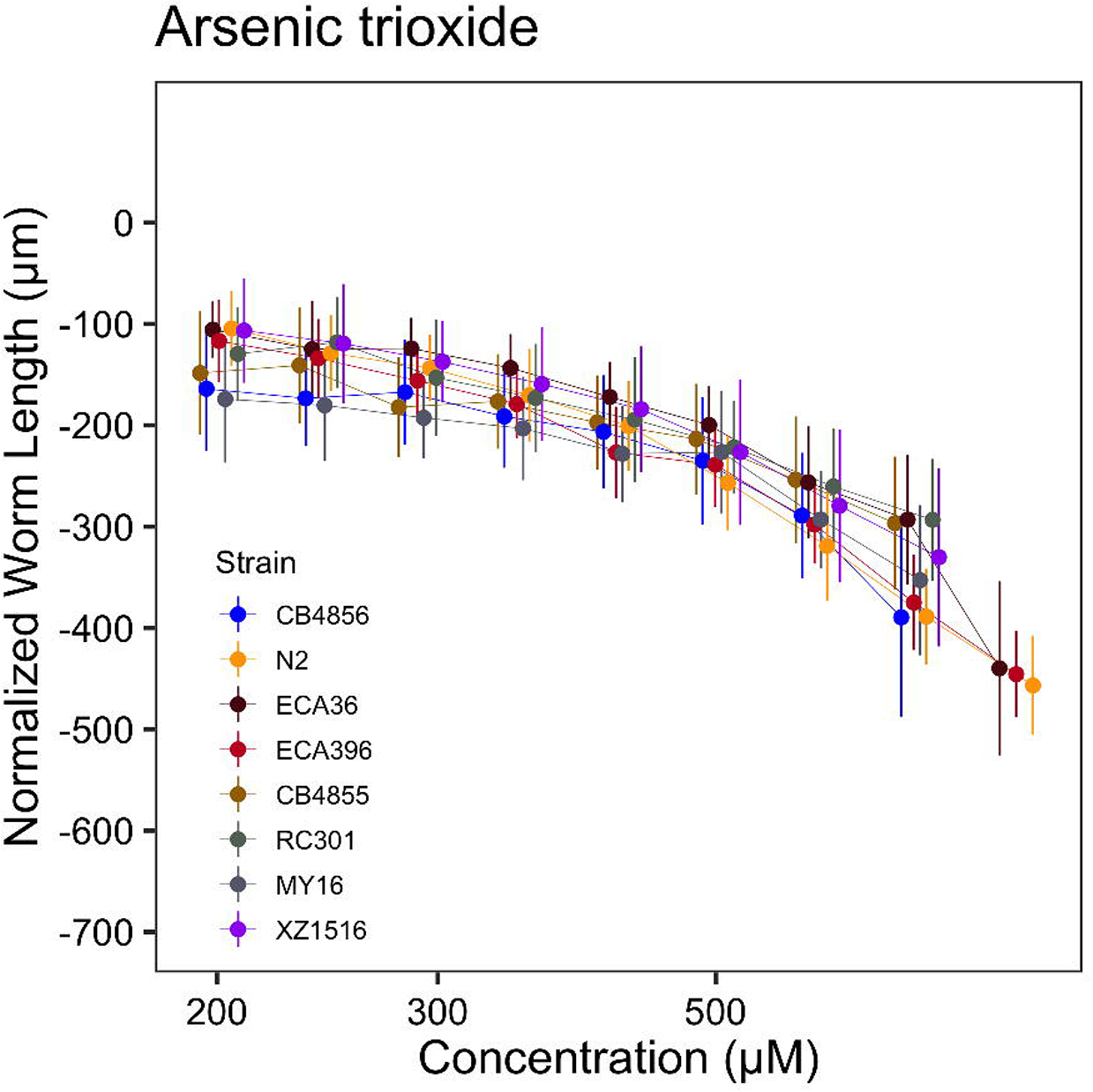
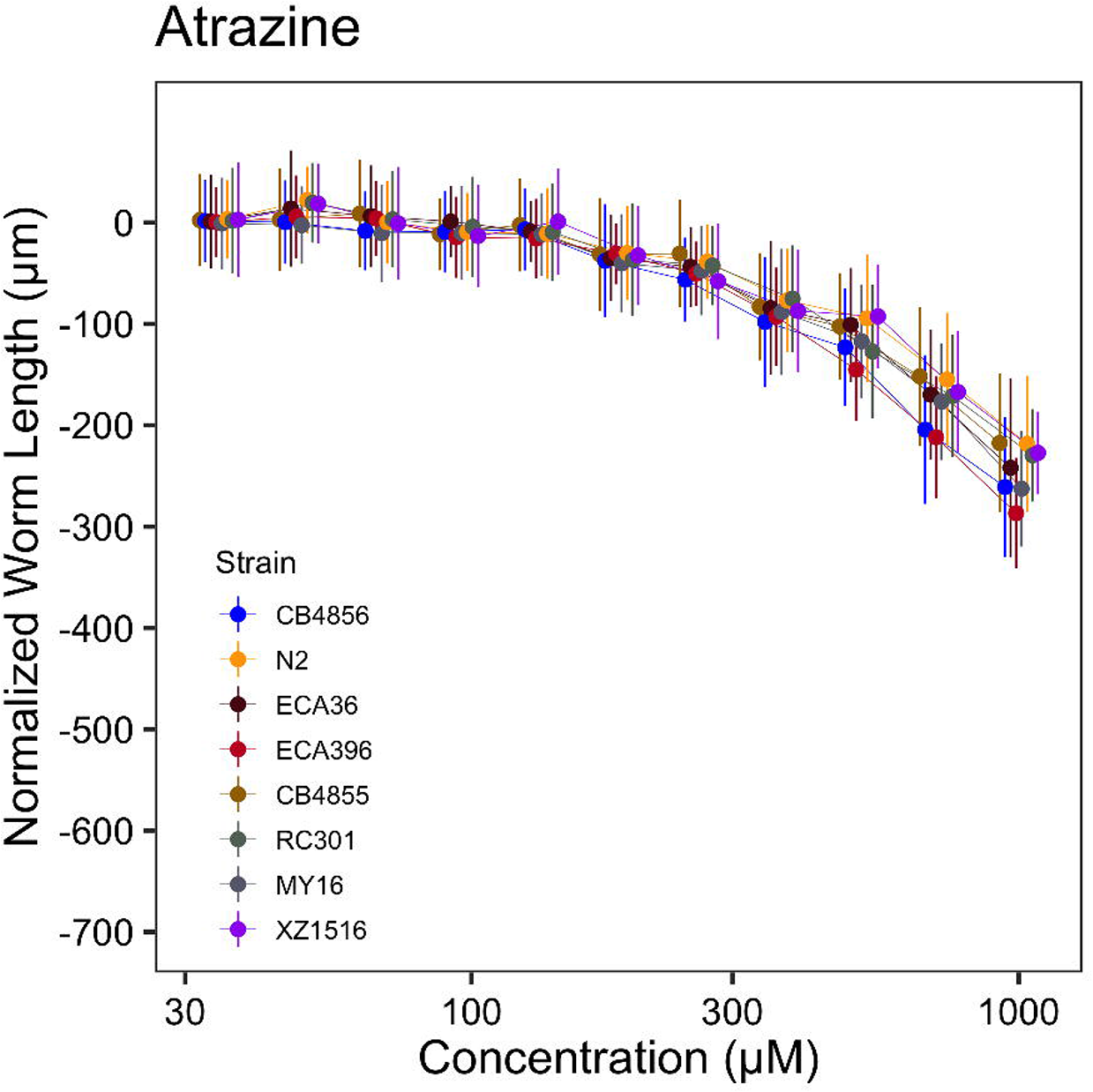
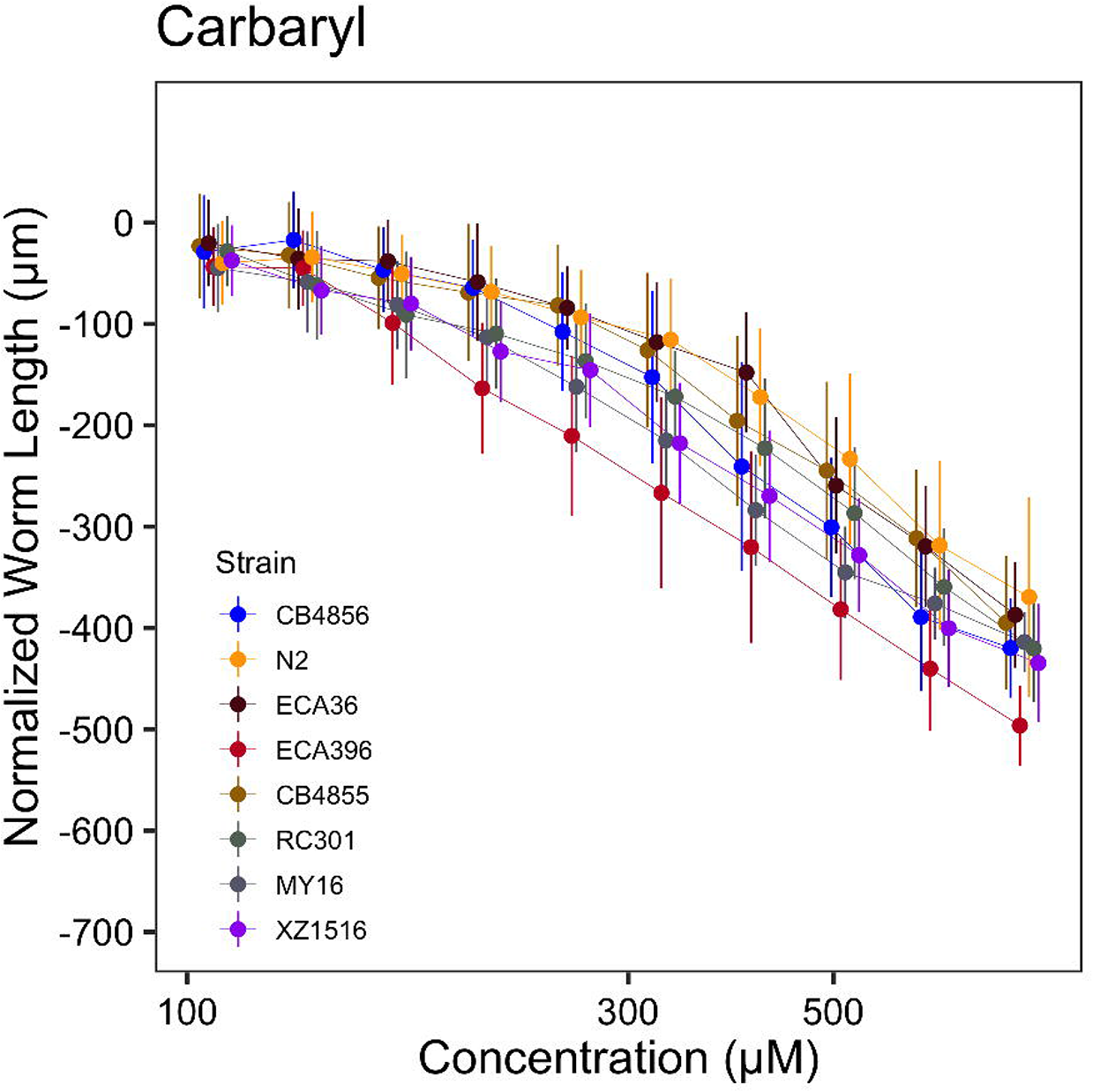
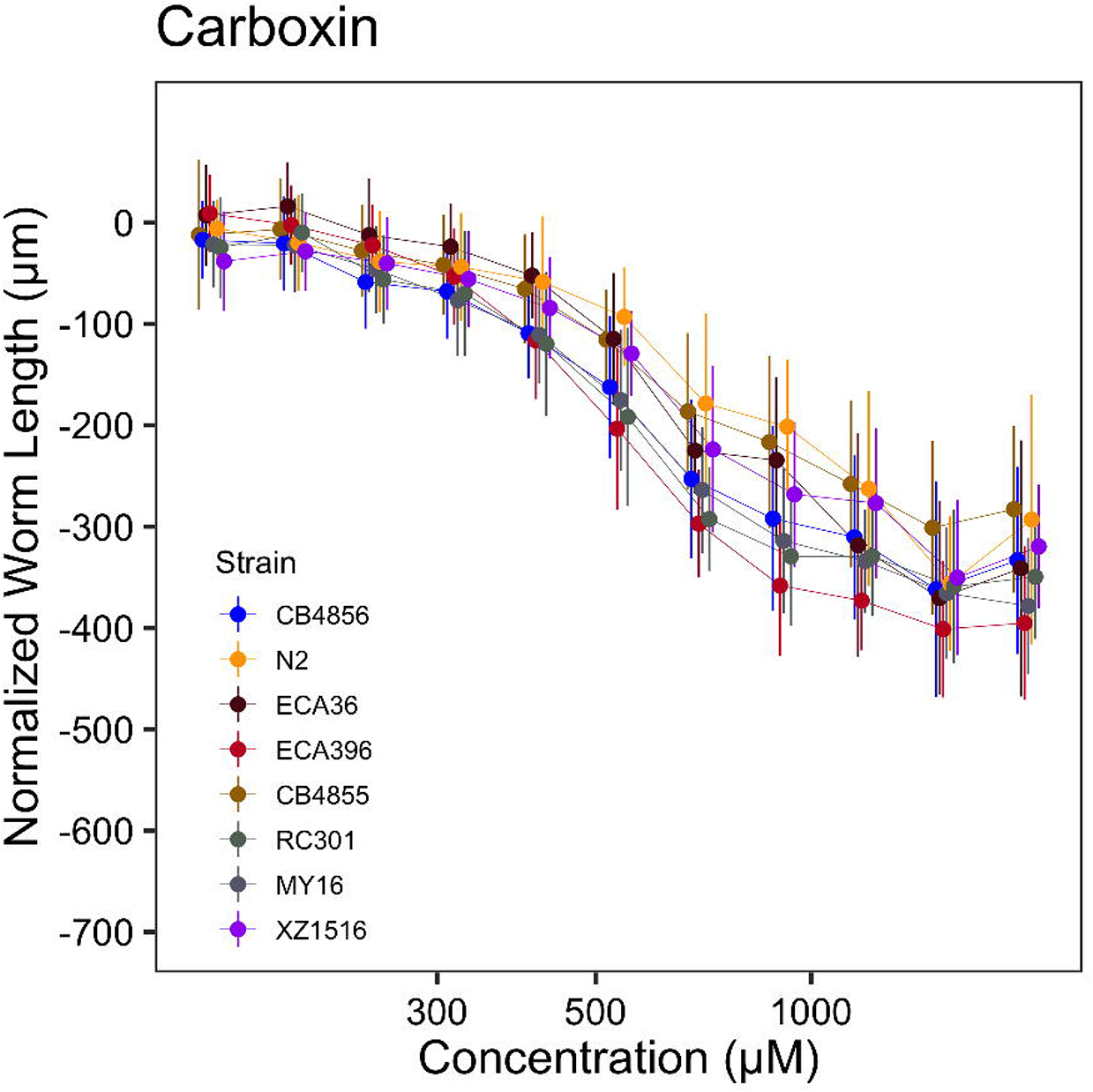
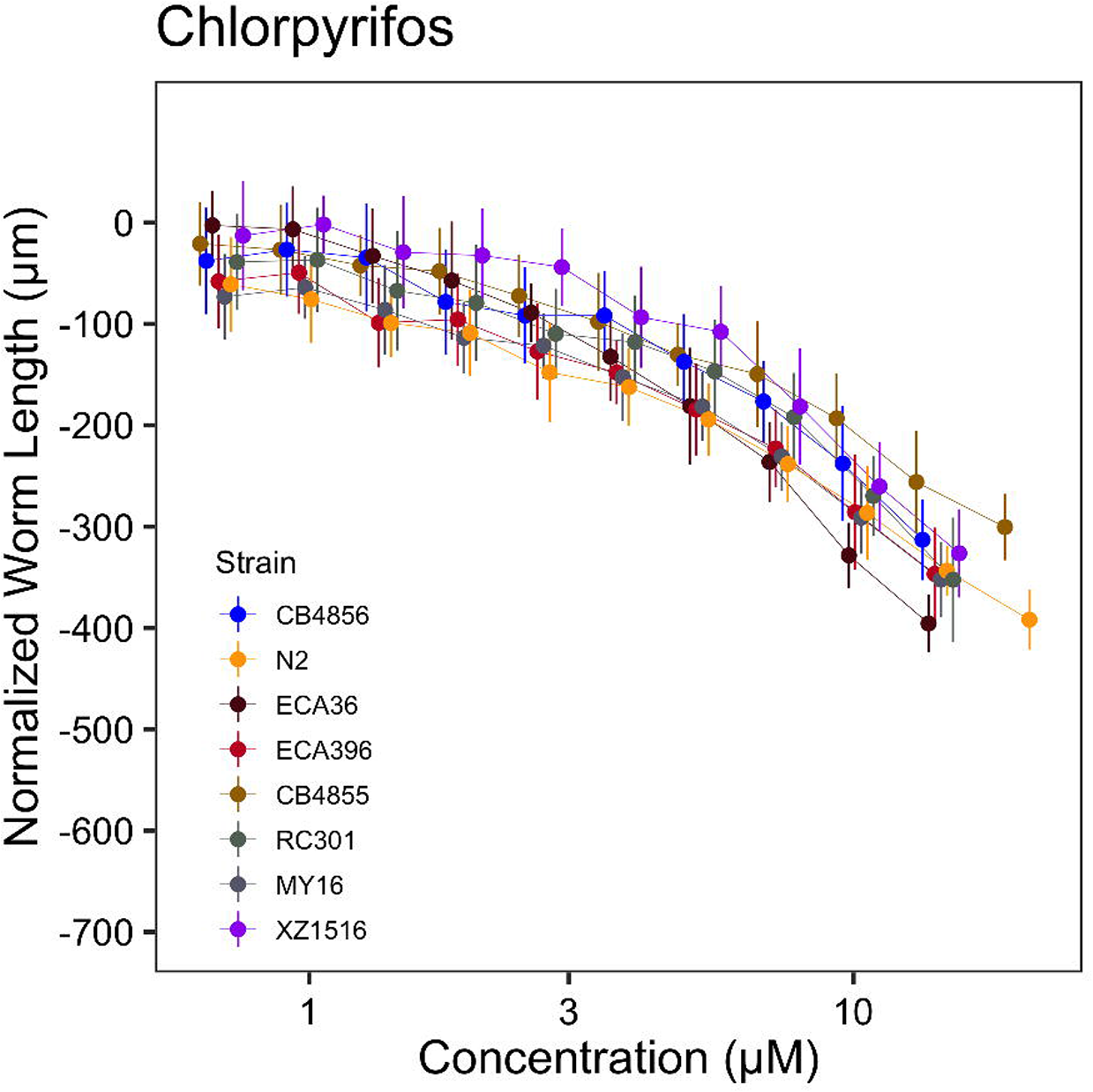
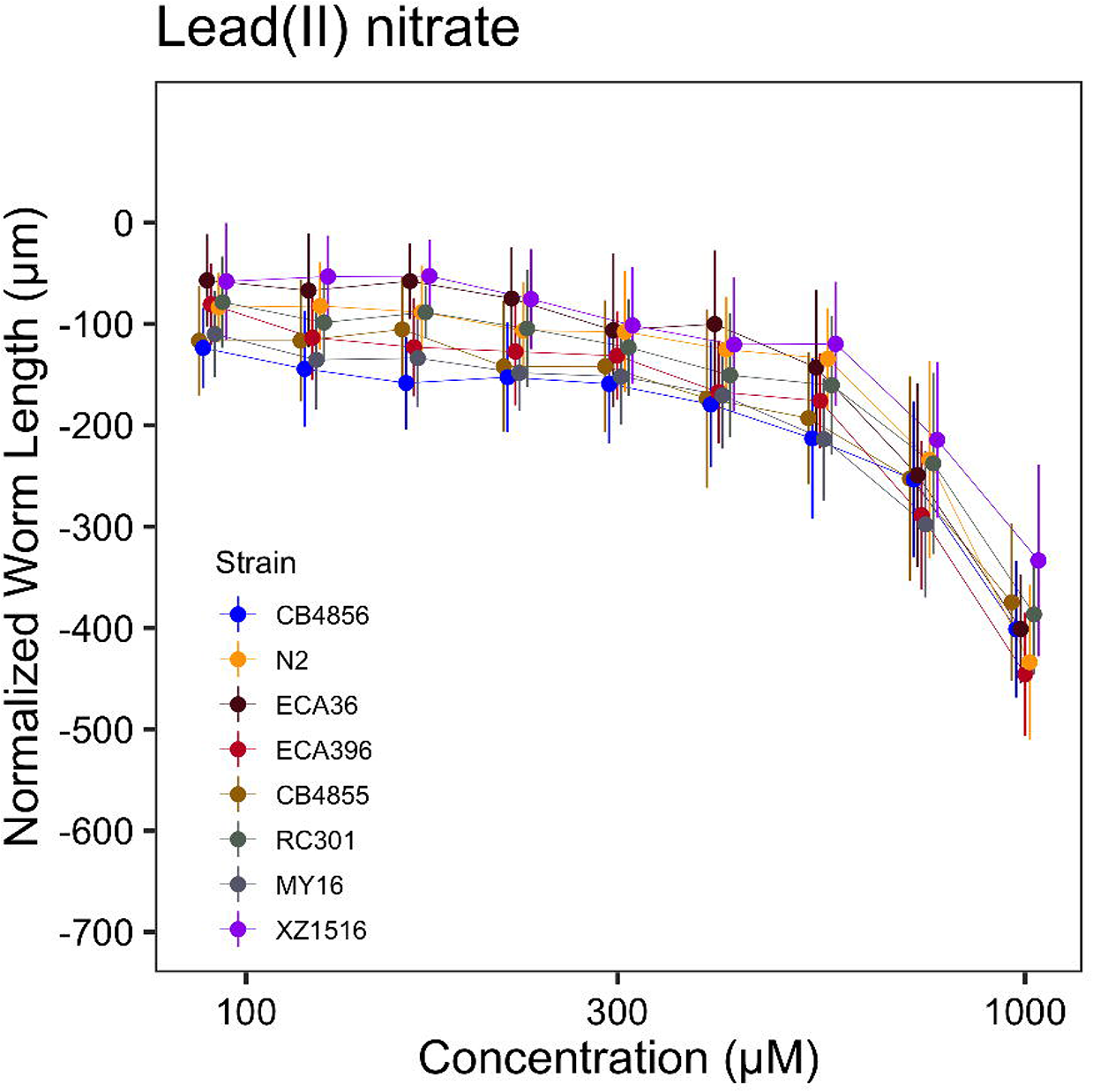
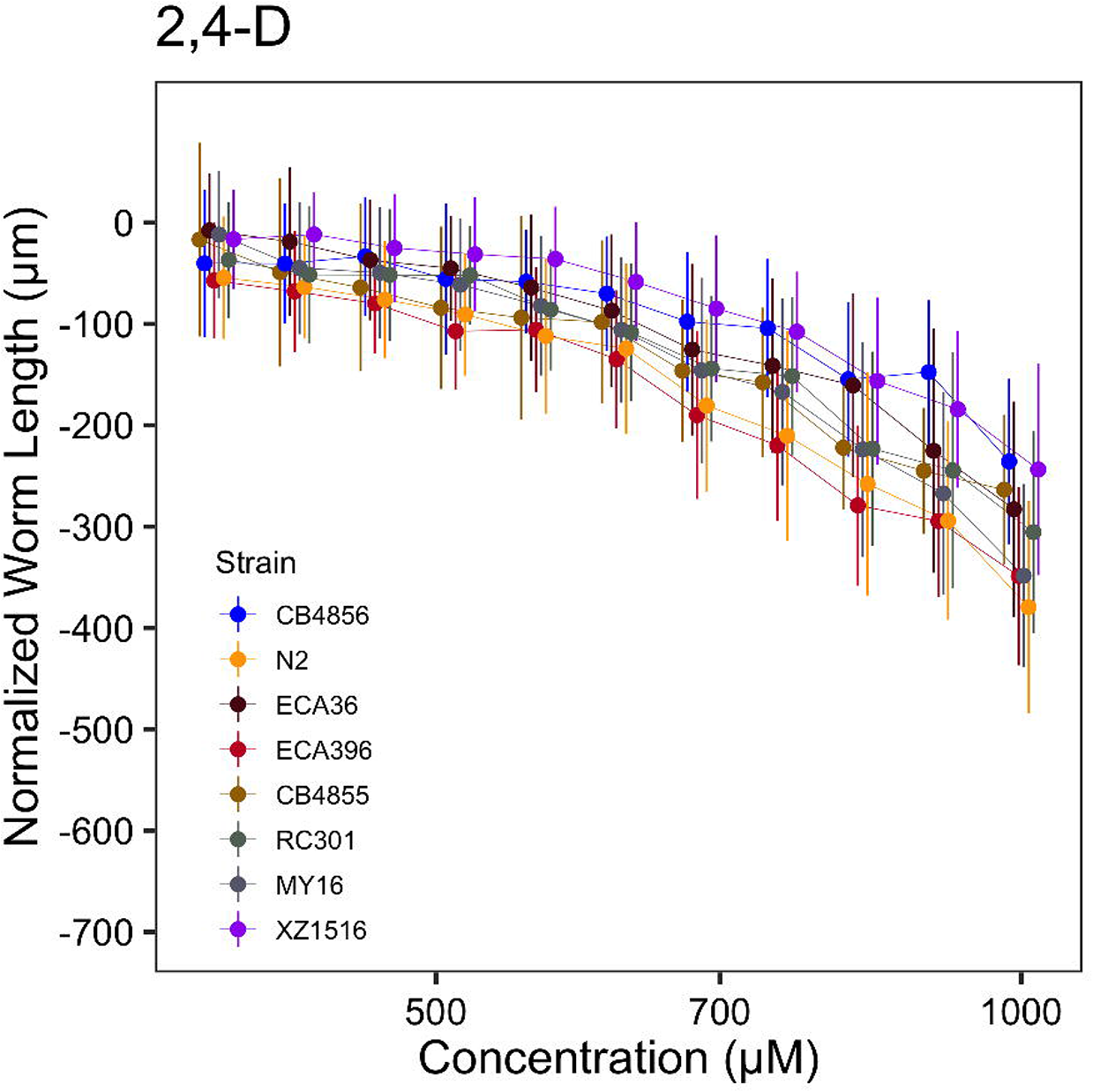
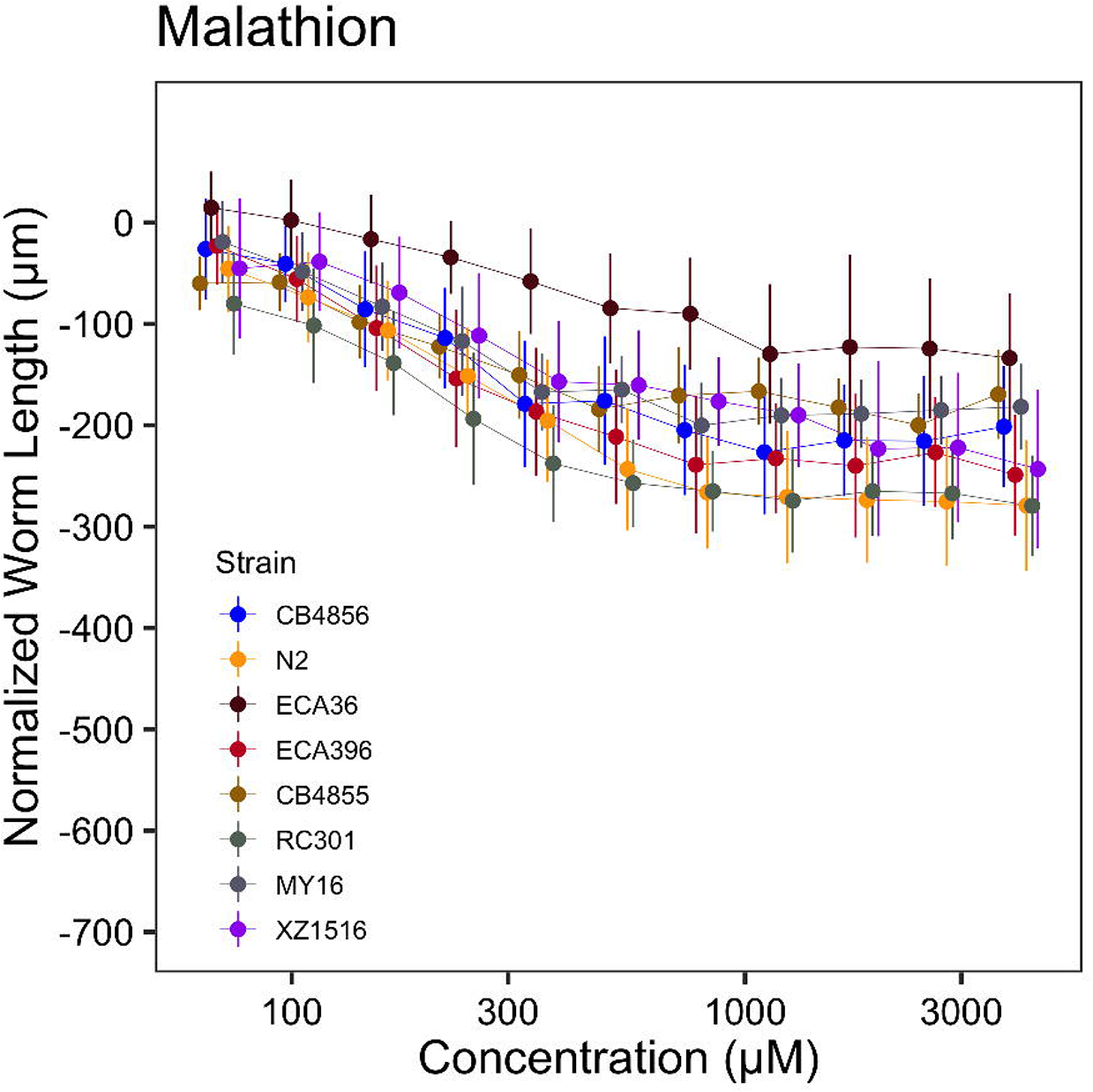
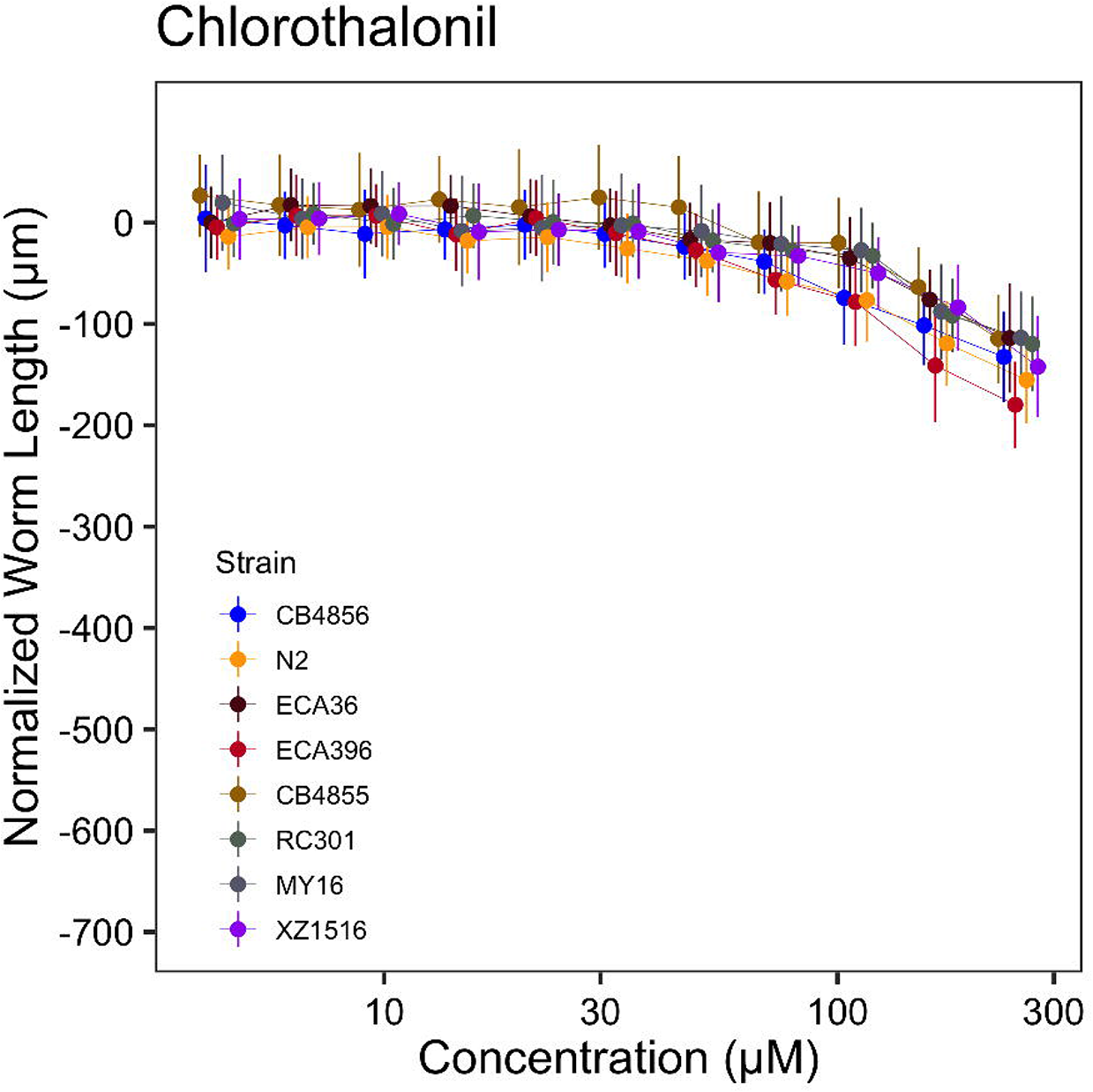
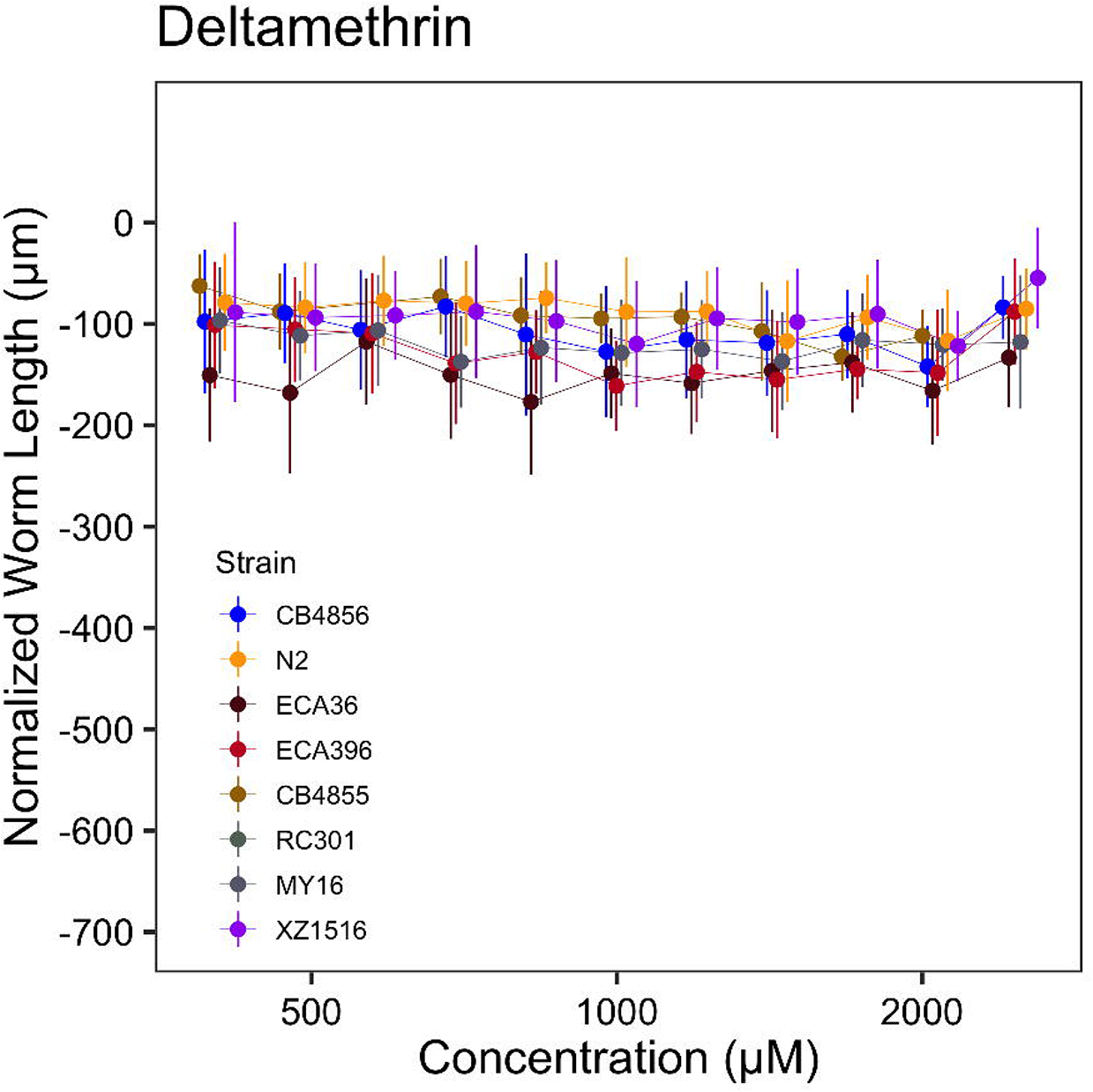
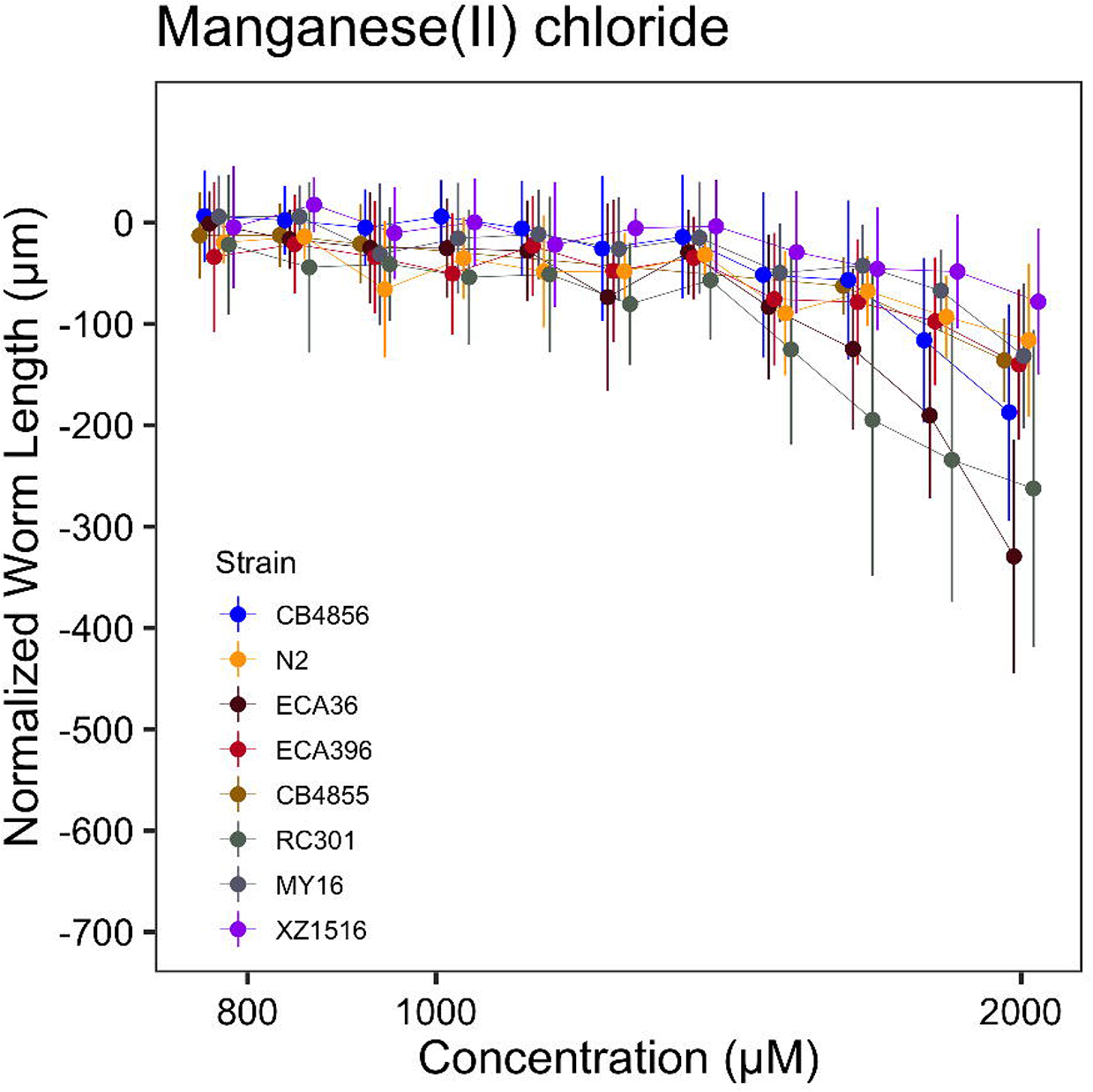
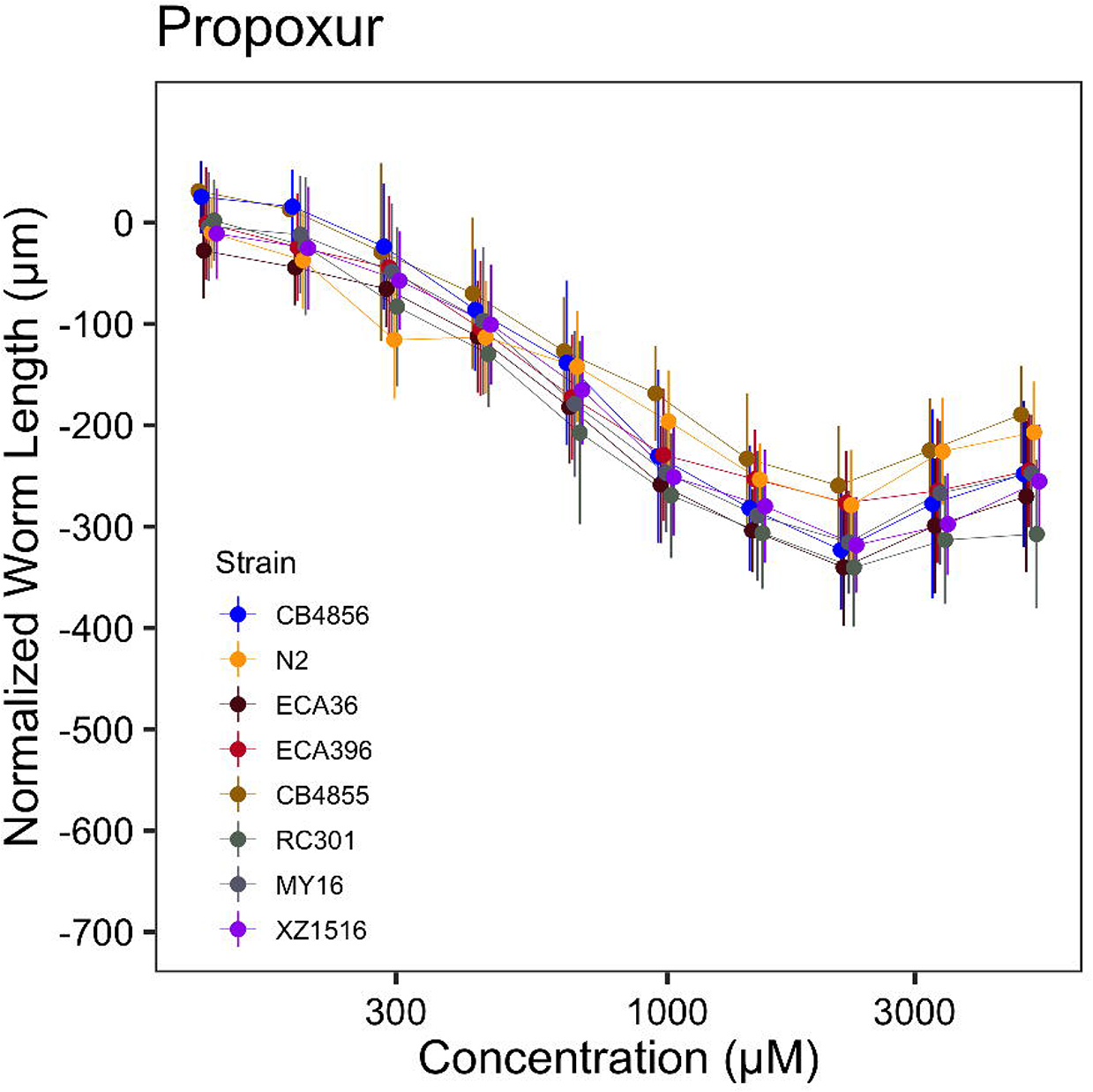
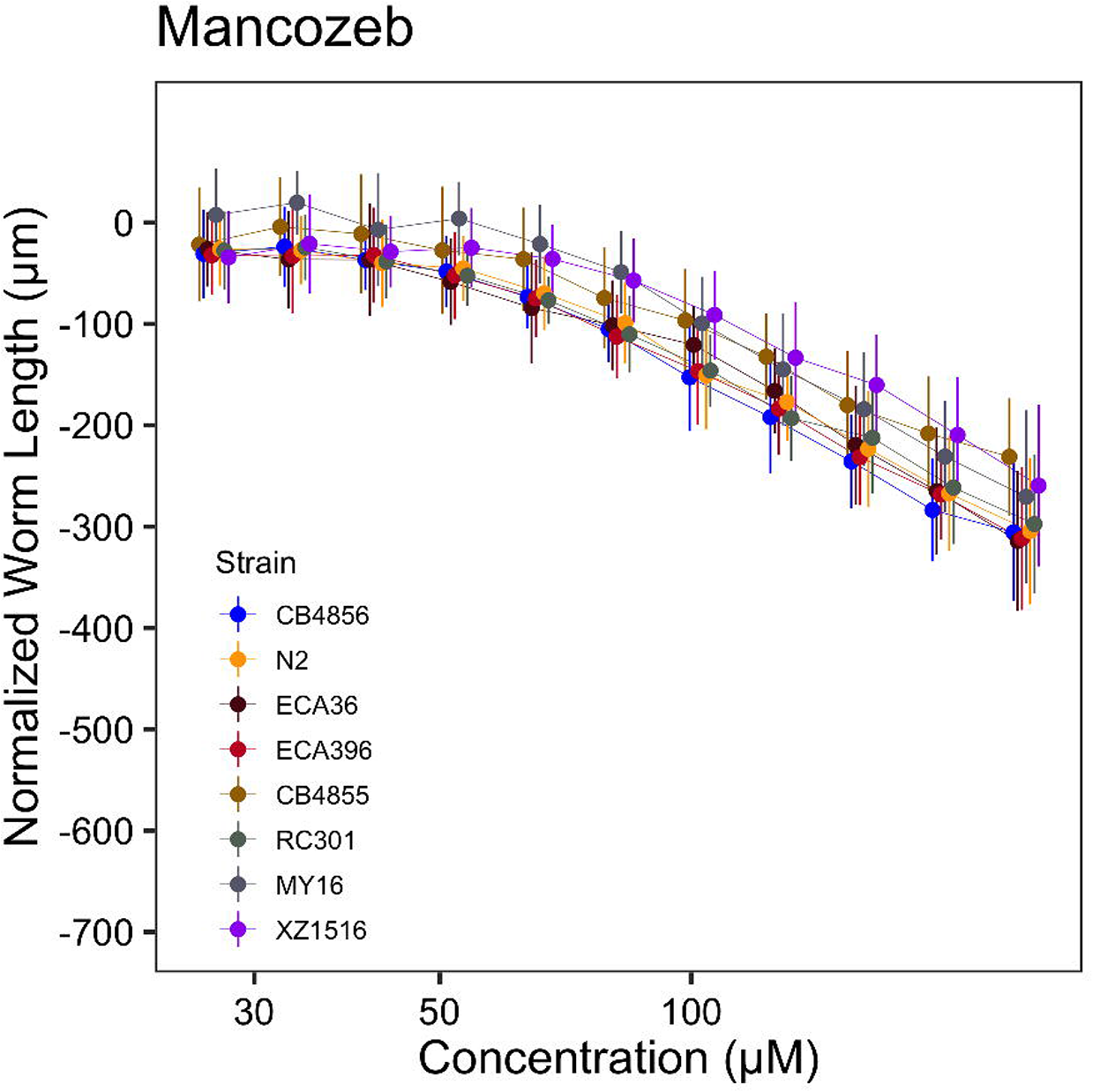
29 – Dose-response curves for all toxicants studied. Normalized animal lengths (y-axis) are plotted for each strain as a function of the dose of toxicant supplied in our high-throughput microscopy assay (x-axis). Each strain is denoted with each color. Lines extending from points represent the standard deviation from the mean response. Statistical normalization of animal lengths is described in *Methods*.

## SUPPLEMENTAL TABLES

**Supplemental Table 1.**
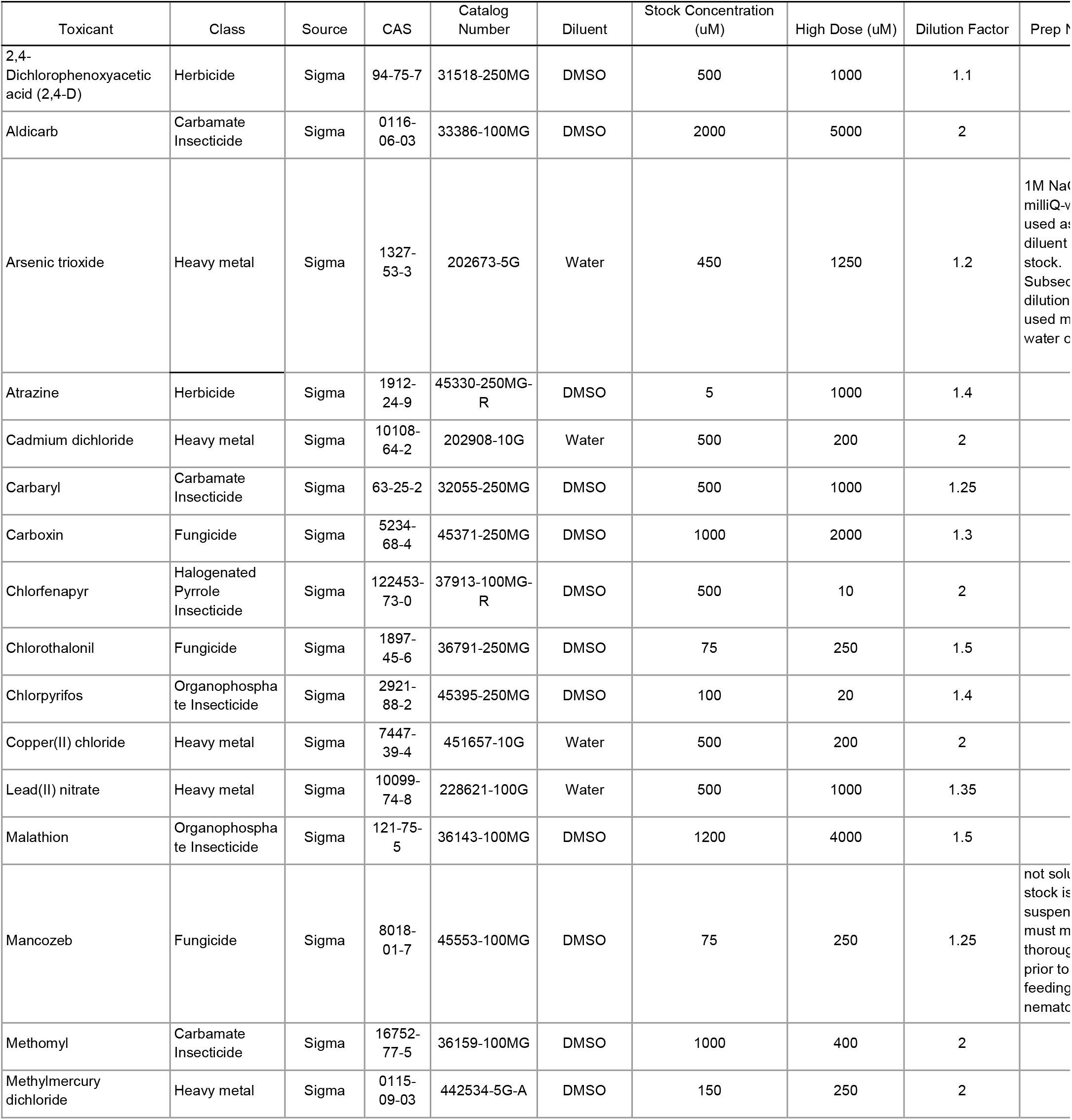
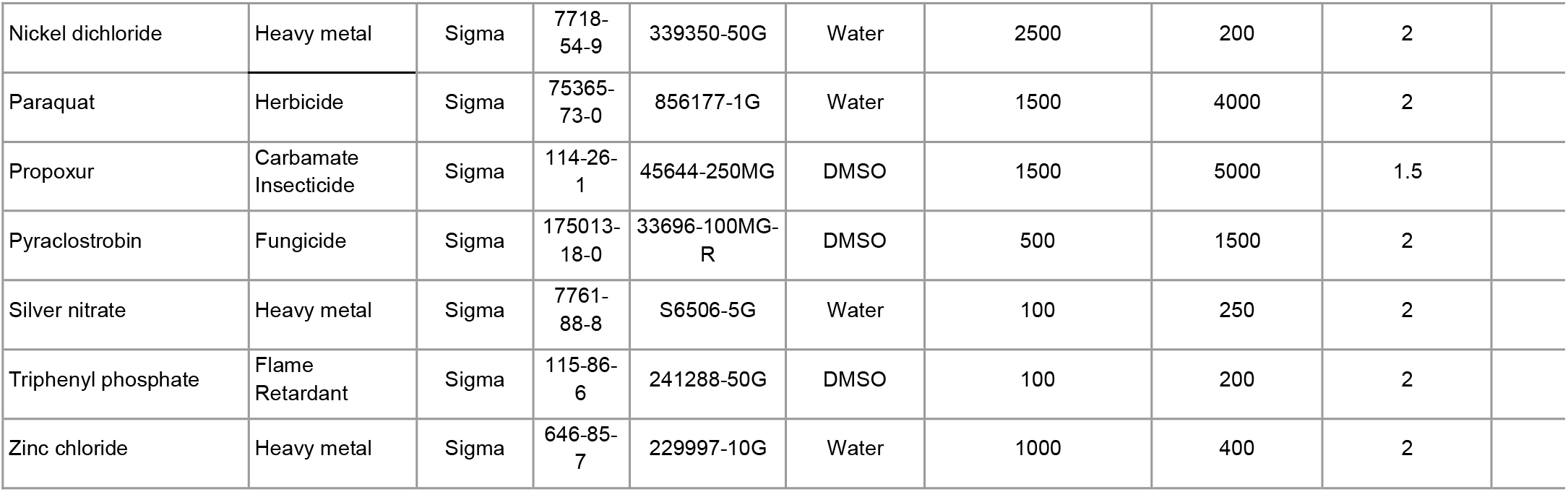
Toxicant stock solution preparation details. The details of each toxicant stock solution preparation are shown.

| Toxicant | Class | Source | CAS | Catalog Number | Diluent | Stock Concentration (uM) | High Dose (uM) | Dilution Factor | Prep t |
| --- | --- | --- | --- | --- | --- | --- | --- | --- | --- |
| 2,4-Dichlorophenoxyacetic acid (2,4-D) | Herbicide | Sigma | 94-75-7 | 31518-250MG | DMSO | 500 | 1000 | 1.1 |  |
| Aldicarb | Carbamate Insecticide | Sigma | 0116-06-03 | 33386-100MG | DMSO | 2000 | 5000 | 2 |  |
| Arsenic trioxide | Heavy metal | Sigma | 1327-53-3 | 202673-5G | Water | 450 | 1250 | 1.2 | 1M NaCl milliQ-water used as diluent stock. Subsequent dilution used in water c |
| Atrazine | Herbicide | Sigma | 1912-24-9 | 45330-250MG-R | DMSO | 5 | 1000 | 1.4 |  |
| Cadmium dichloride | Heavy metal | Sigma | 10108-64-2 | 202908-10G | Water | 500 | 200 | 2 |  |
| Carbaryl | Carbamate Insecticide | Sigma | 63-25-2 | 32055-250MG | DMSO | 500 | 1000 | 1.25 |  |
| Carboxin | Fungicide | Sigma | 5234-68-4 | 45371-250MG | DMSO | 1000 | 2000 | 1.3 |  |
| Chlorfenapyr | Halogenated Pyrrole Insecticide | Sigma | 122453-73-0 | 37913-100MG-R | DMSO | 500 | 10 | 2 |  |
| Chlorothalonil | Fungicide | Sigma | 1897-45-6 | 36791-250MG | DMSO | 75 | 250 | 1.5 |  |
| Chlorpyrifos | Organophosphate Insecticide | Sigma | 2921-88-2 | 45395-250MG | DMSO | 100 | 20 | 1.4 |  |
| Copper(II) chloride | Heavy metal | Sigma | 7447-39-4 | 451657-10G | Water | 500 | 200 | 2 |  |
| Lead(II) nitrate | Heavy metal | Sigma | 10099-74-8 | 228621-100G | Water | 500 | 1000 | 1.35 |  |
| Malathion | Organophosphate Insecticide | Sigma | 121-75-5 | 36143-100MG | DMSO | 1200 | 4000 | 1.5 |  |
| Mancozeb | Fungicide | Sigma | 8018-01-7 | 45553-100MG | DMSO | 75 | 250 | 1.25 | not solt stock is suspen must m thorough prior to feeding nematoc |
| Methomyl | Carbamate Insecticide | Sigma | 16752-77-5 | 36159-100MG | DMSO | 1000 | 400 | 2 |  |
| Methylmercury dichloride | Heavy metal | Sigma | 0115-09-03 | 442534-5G-A | DMSO | 150 | 250 | 2 |  |
| Nickel dichloride | Heavy metal | Sigma | 7718-54-9 | 339350-50G | Water | 2500 | 200 | 2 |  |
| Paraquat | Herbicide | Sigma | 75365-73-0 | 856177-1G | Water | 1500 | 4000 | 2 |  |
| Propoxur | Carbamate Insecticide | Sigma | 114-26-1 | 45644-250MG | DMSO | 1500 | 5000 | 1.5 |  |
| Pyraclostrobin | Fungicide | Sigma | 175013-18-0 | 33696-100MG-R | DMSO | 500 | 1500 | 2 |  |
| Silver nitrate | Heavy metal | Sigma | 7761-88-8 | S6506-5G | Water | 100 | 250 | 2 |  |
| Triphenyl phosphate | Flame Retardant | Sigma | 115-86-6 | 241288-50G | DMSO | 100 | 200 | 2 |  |
| Zinc chloride | Heavy metal | Sigma | 646-85-7 | 229997-10G | Water | 1000 | 400 | 2 |  |

**Supplemental Table 2.** Overall and strain-specific LOAEL for 23 toxicants among *C. elegans* strains

| Toxicant Class | Toxicant | Population-wide LOAEL | CB4856 | ECA248 | ECA36 | ECA396 | MY16 | PD1074 | RC301 | XZ1516 |
| --- | --- | --- | --- | --- | --- | --- | --- | --- | --- | --- |
| Flame Retardant | Triphenyl phosphate | 1.56 | 100 | 100 | 100 | 100 | 1.56 | 100 | 1.56 | 100 |
| Fungicide | Pyraclostrobin | 11.72 | 11.72 | 1500 | 11.72 | 11.72 | 11.72 | 11.72 | 1500 | 11.72 |
| Fungicide | Carboxin | 1183.43 | 1183.43 | 1183.43 | 1183.43 | 1183.43 | 1183.43 | 1183.43 | 1183.43 | 1183.43 |
| Fungicide | Chlorothalonil | 111.11 | 111.11 | 166.67 | 111.11 | 111.11 | 166.67 | 111.11 | 111.11 | 111.11 |
| Fungicide | Mancozeb | 102.4 | 102.4 | 102.4 | 102.4 | 102.4 | 102.4 | 102.4 | 102.4 | 102.4 |
| Herbicide | Paraquat | 1000 | 1000 | 1000 | 1000 | 1000 | 1000 | 1000 | 1000 | 1000 |
| Herbicide | Atrazine | 1000 | 1000 | 1000 | 1000 | 1000 | 1000 | 1000 | 1000 | 1000 |
| Herbicide | 2,4-D | 1000 | 1000 | 1000 | 1000 | 1000 | 1000 | 1000 | 1000 | 1000 |
| Insecticide | Aldicarb | 1250 | 1250 | 1250 | 1250 | 1250 | 1250 | 1250 | 1250 | 1250 |
| Insecticide | Chlorfenapyr | 0.31 | 0.31 | 0.16 | 0.63 | 0.63 | 0.31 | 0.31 | 0.31 | 0.31 |
| Insecticide | Methomyl | 100 | 100 | 100 | 100 | 100 | 100 | 100 | 100 | 100 |
| Insecticide | Carbaryl | 107.37 | 209.72 | 209.72 | 167.77 | 167.77 | 107.37 | 209.72 | 167.77 | 167.77 |
| Insecticide | Chlorpyrifos | 0.69 | 0.69 | 1.36 | 1.36 | 0.69 | 0.69 | 0.69 | 0.69 | 10.2 |
| Insecticide | Malathion | 104.05 | 1185.19 | 104.05 | 1185.19 | 104.05 | 1185.19 | 104.05 | 104.05 | 1185.19 |
| Insecticide | Propoxur | 1481.48 | 1481.48 | 1481.48 | 1481.48 | 1481.48 | 1481.48 | 1481.48 | 1481.48 | 1481.48 |
| Metal | Cadmium chloride | 100 | 100 | 100 | 100 | 100 | 100 | 100 | 100 | 100 |
| Metal | Copper(II) chloride | 100 | 100 | 100 | 100 | 100 | 100 | 100 | 100 | 100 |
| Metal | Methylmercury chloride | 0.98 | 1.95 | 1.95 | 1.95 | 1.95 | 1.95 | 1.95 | 15.63 | 15.63 |
| Metal | Silver nitrate | 0.49 | 0.98 | 125 | 125 | 0.98 | 0.98 | 1.95 | 1.95 | 125 |
| Metal | Nickel chloride | 100 | 100 | 100 | 100 | 100 | 100 | 100 | 100 | 200 |
| Metal | Zinc chloride | 100 | 100 | 100 | 100 | 100 | 100 | 100 | 100 | 100 |
| Metal | Arsenic trioxide | 201.88 | 201.88 | 201.88 | 201.88 | 201.88 | 201.88 | 201.88 | 201.88 | 201.88 |
| Metal | Lead(II) nitrate | 1000 | 1000 | 1000 | 1000 | 1000 | 1000 | 1000 | 1000 | 1000 |

**Supplemental Table 3.**
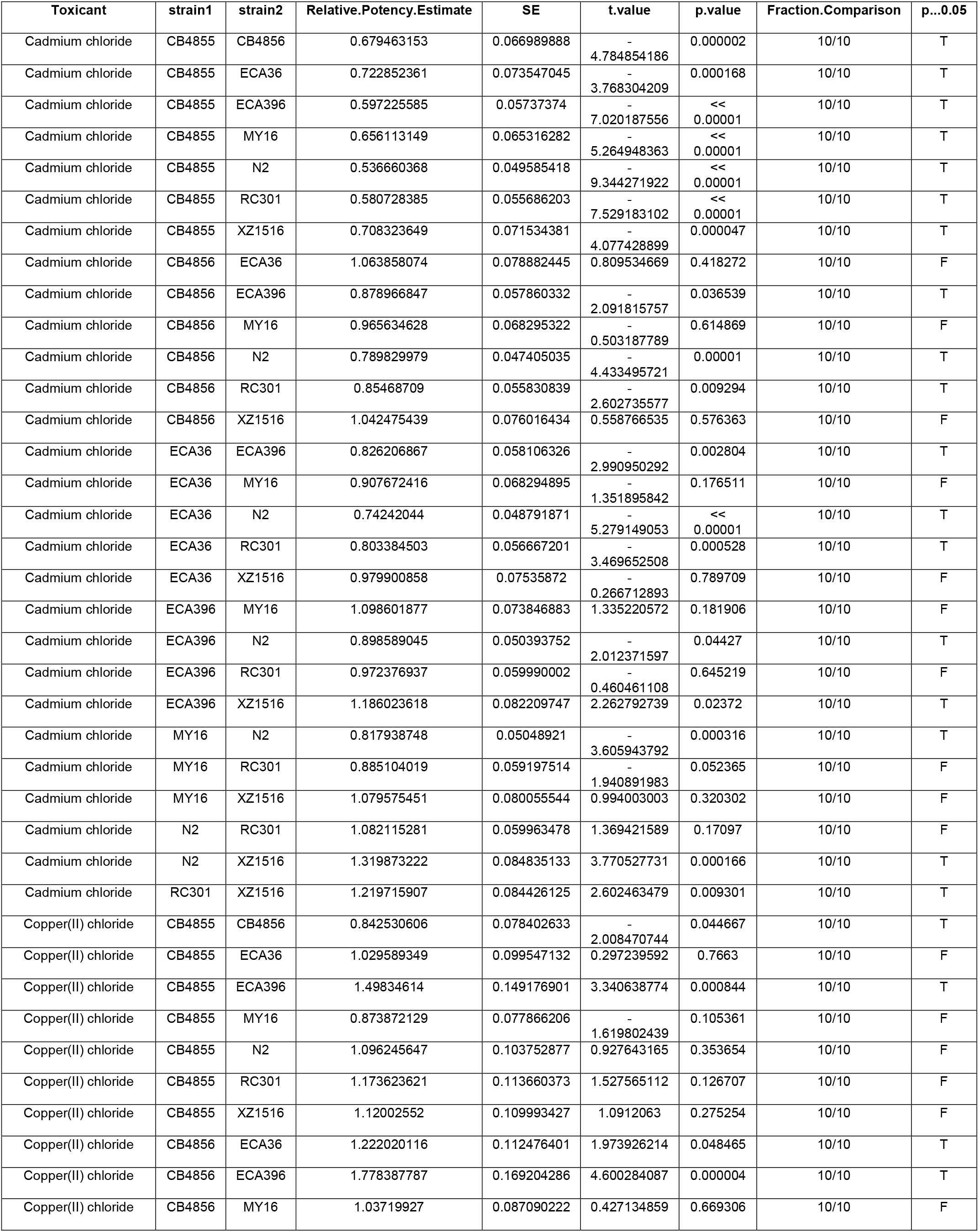
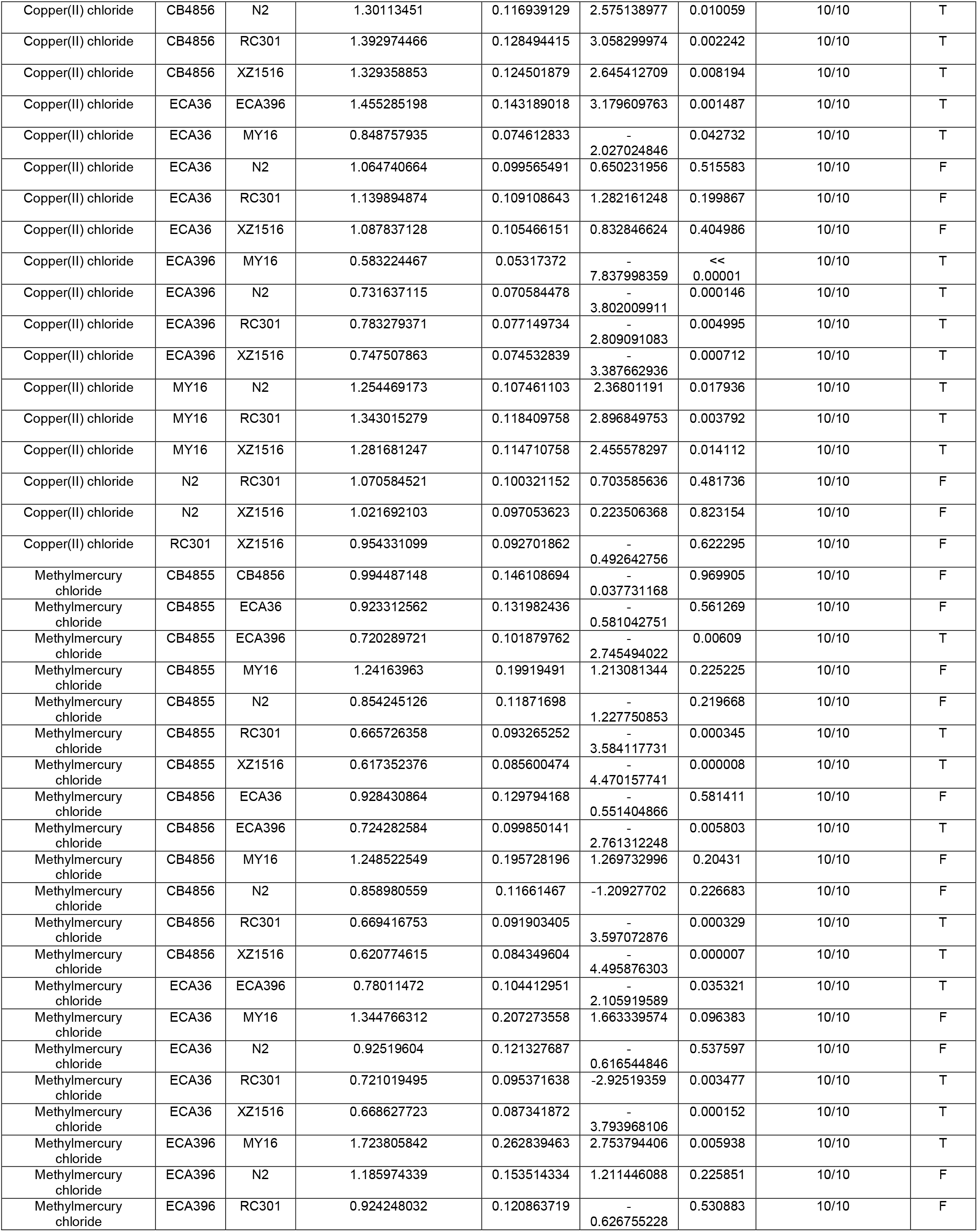
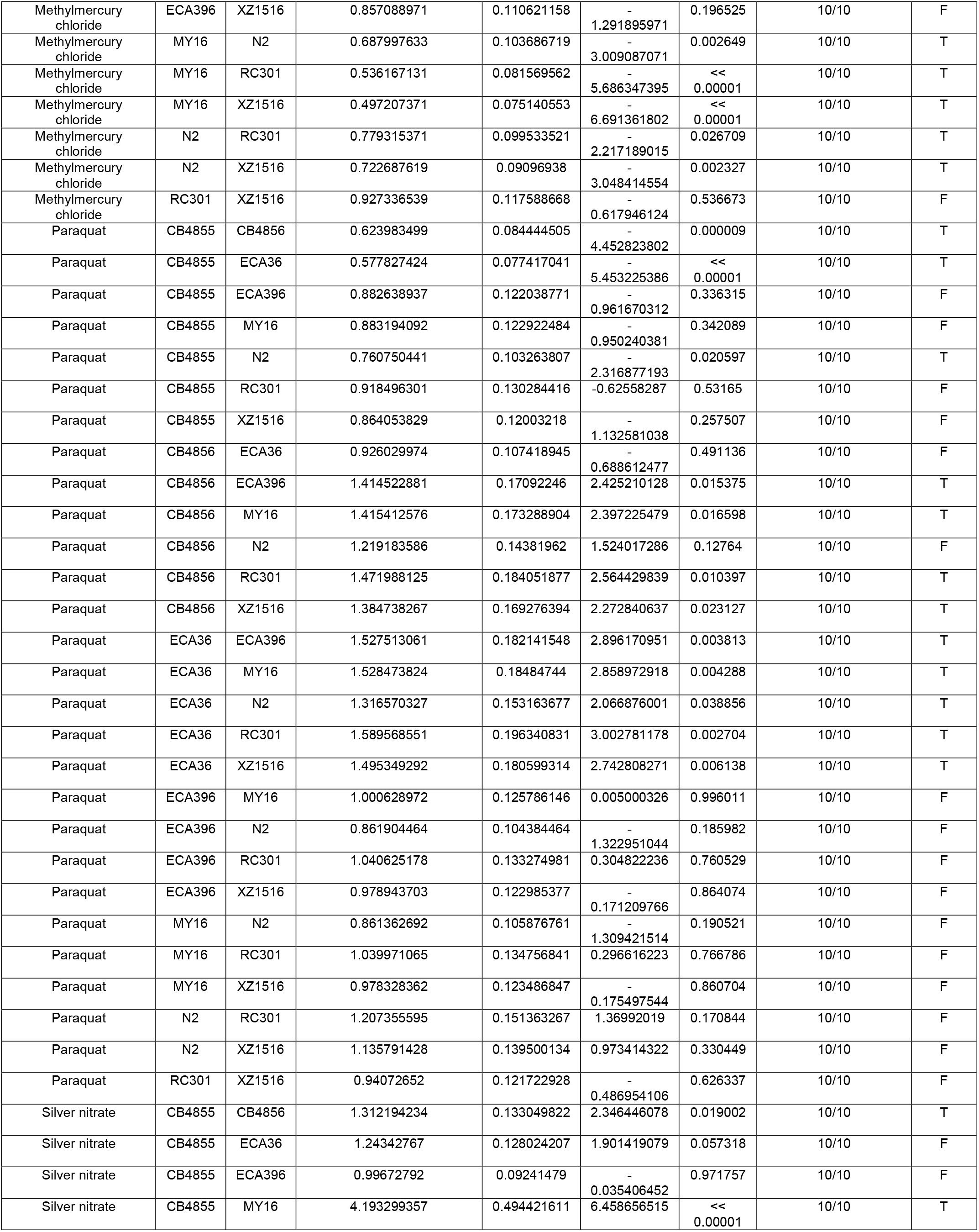
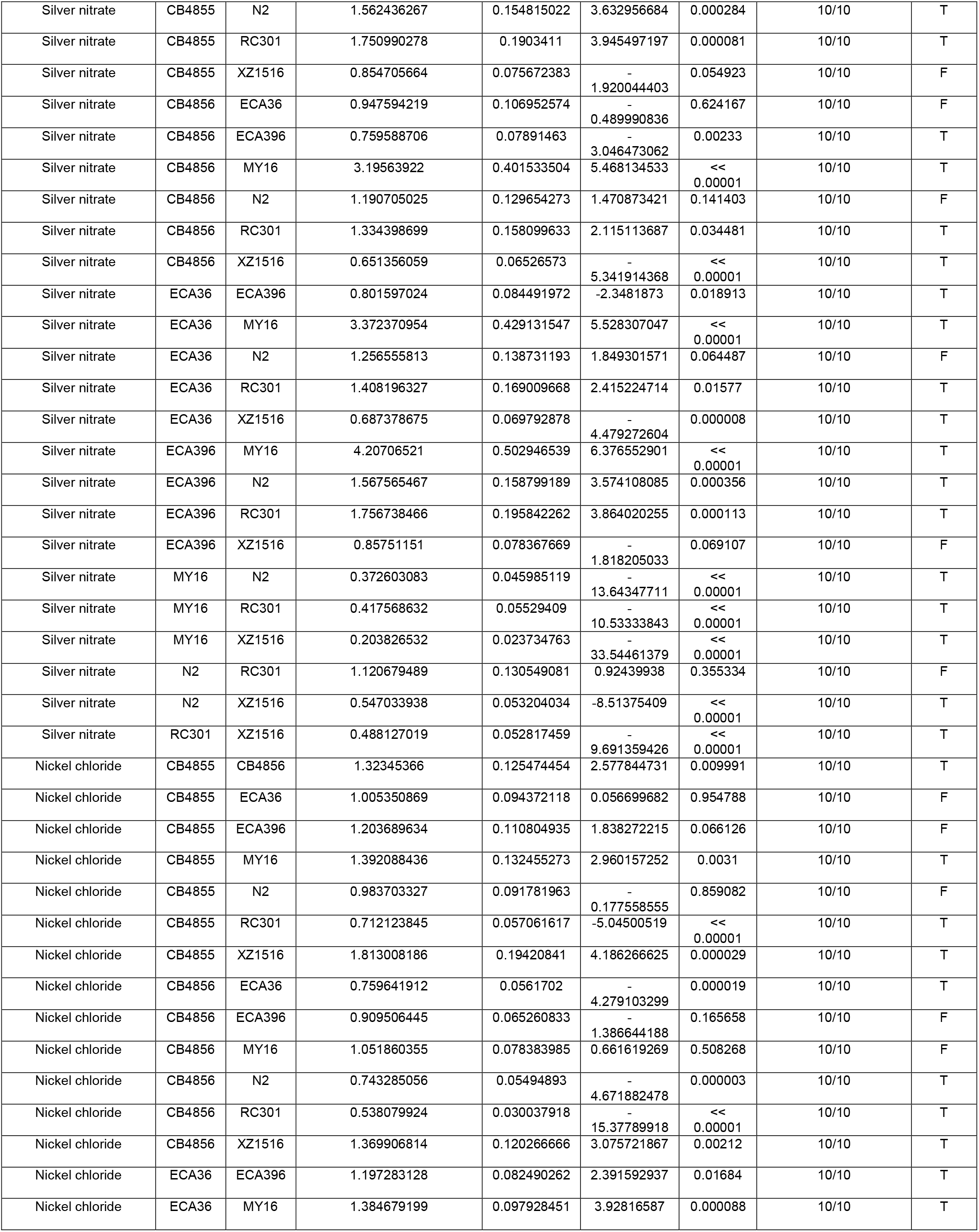
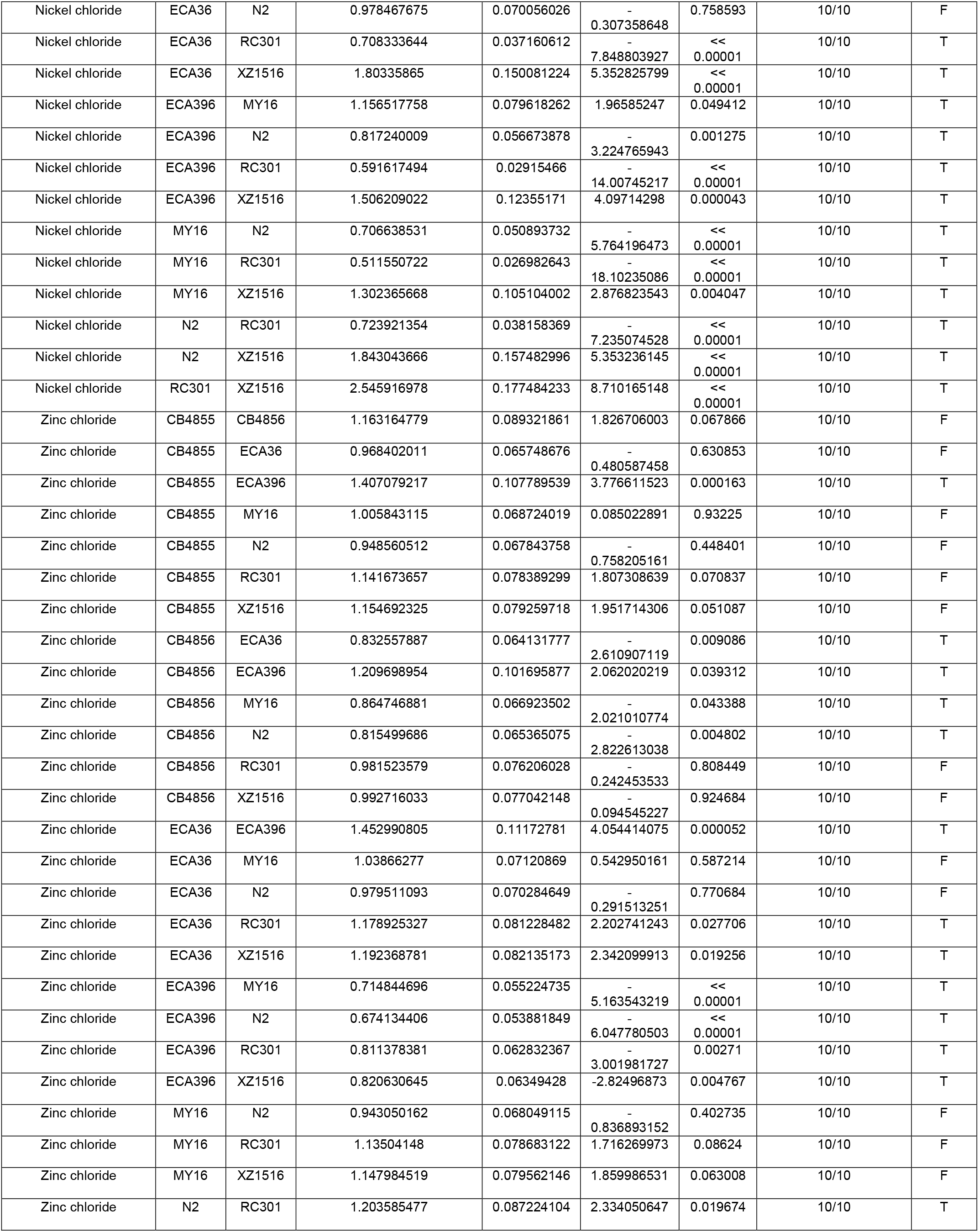
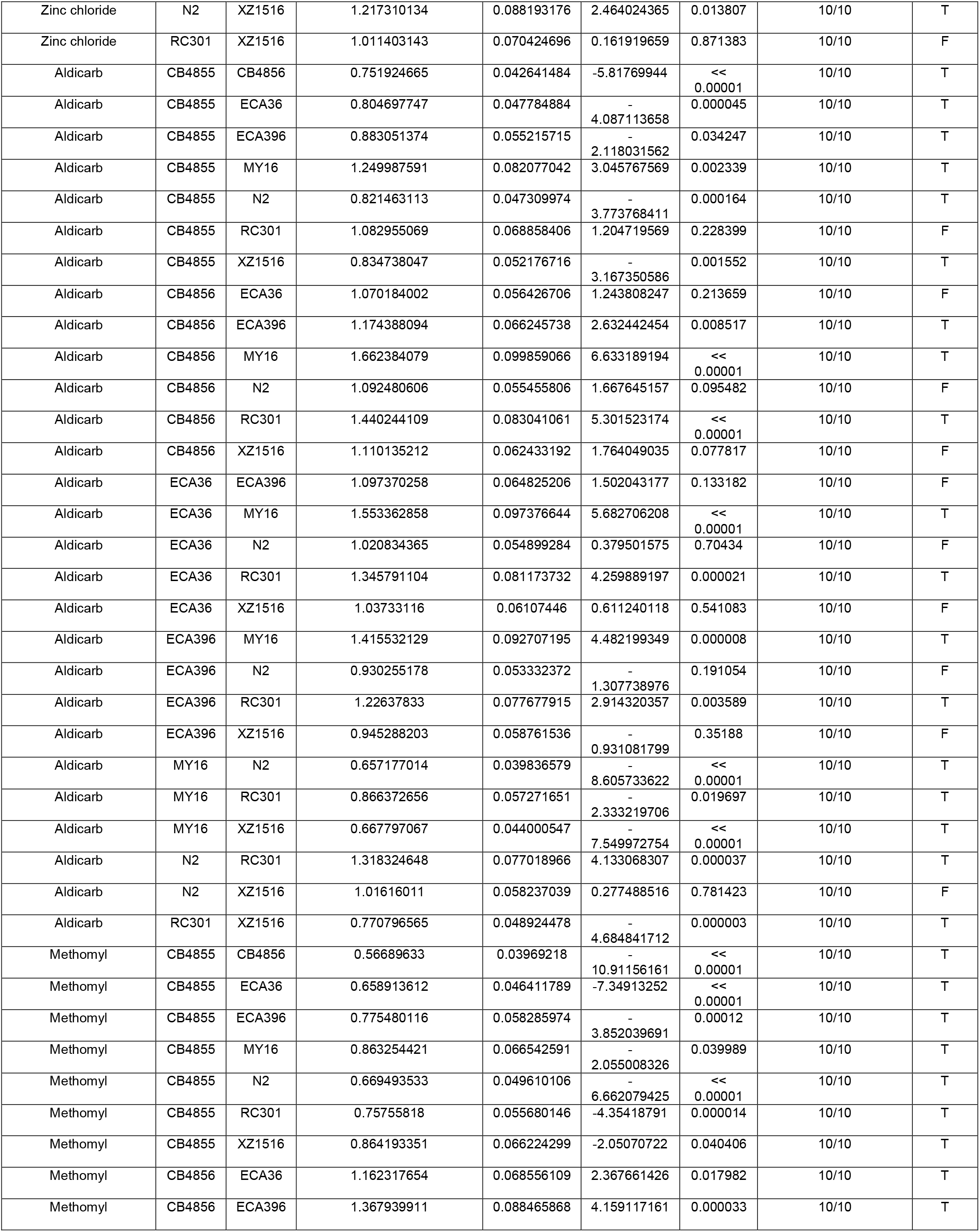
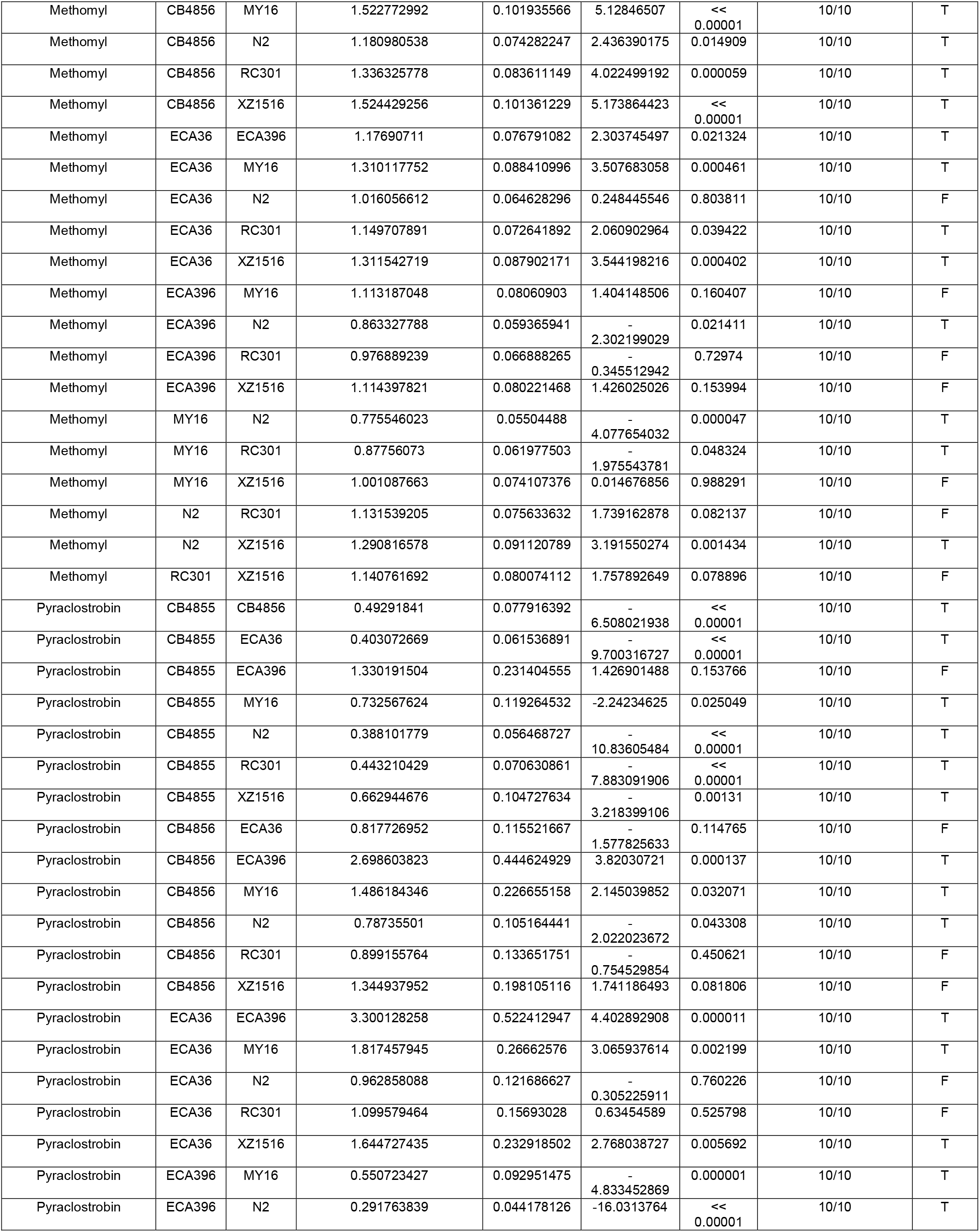
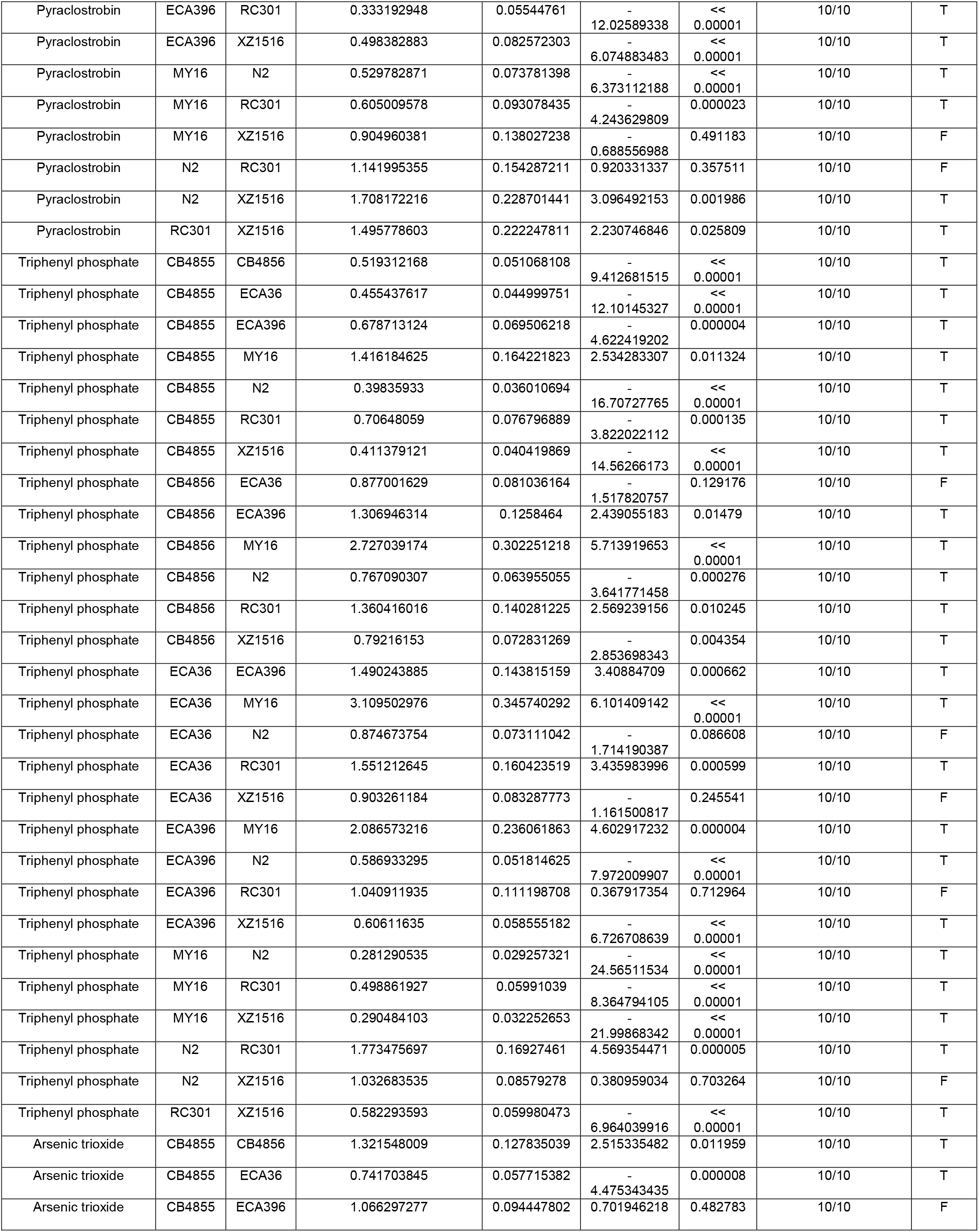
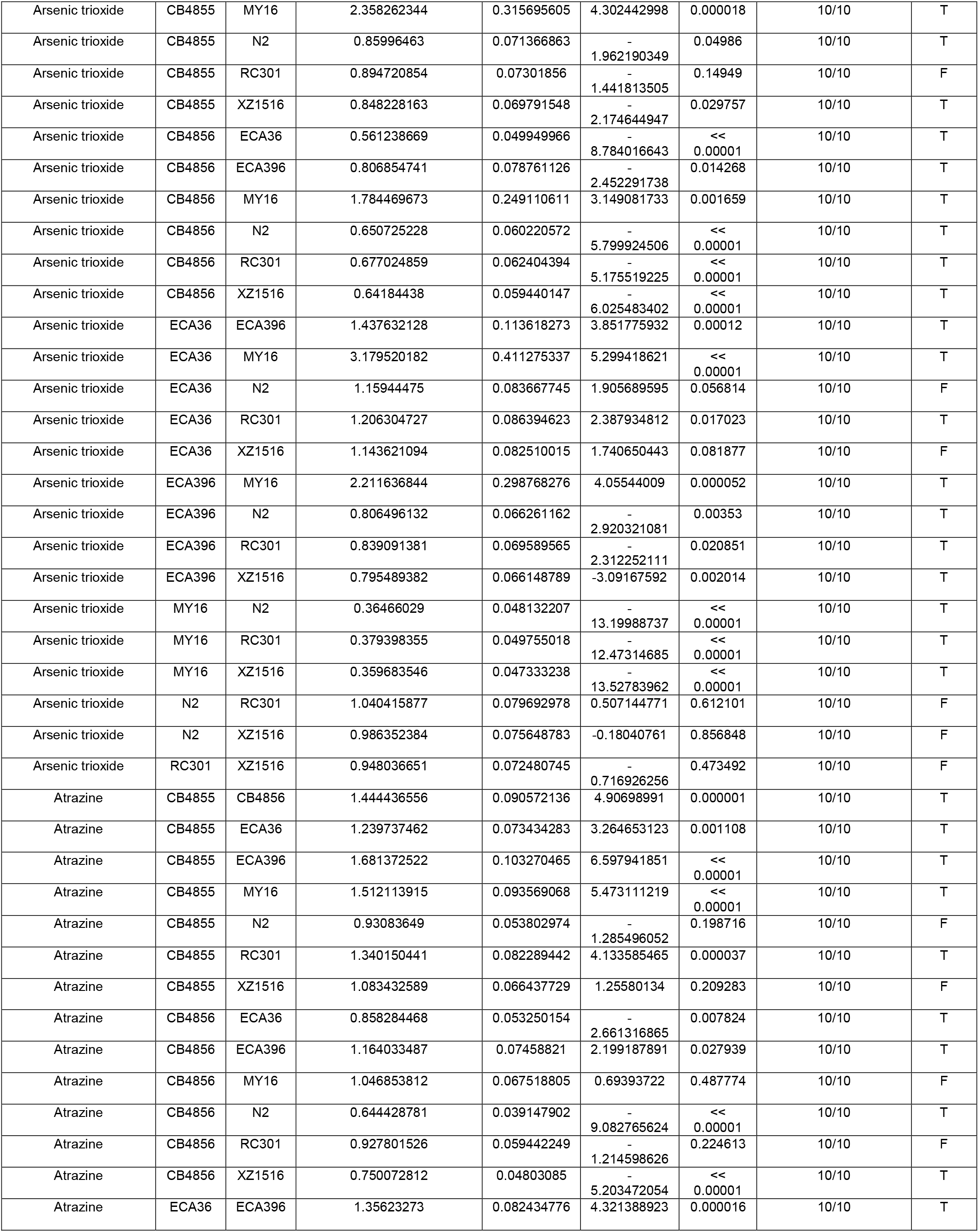
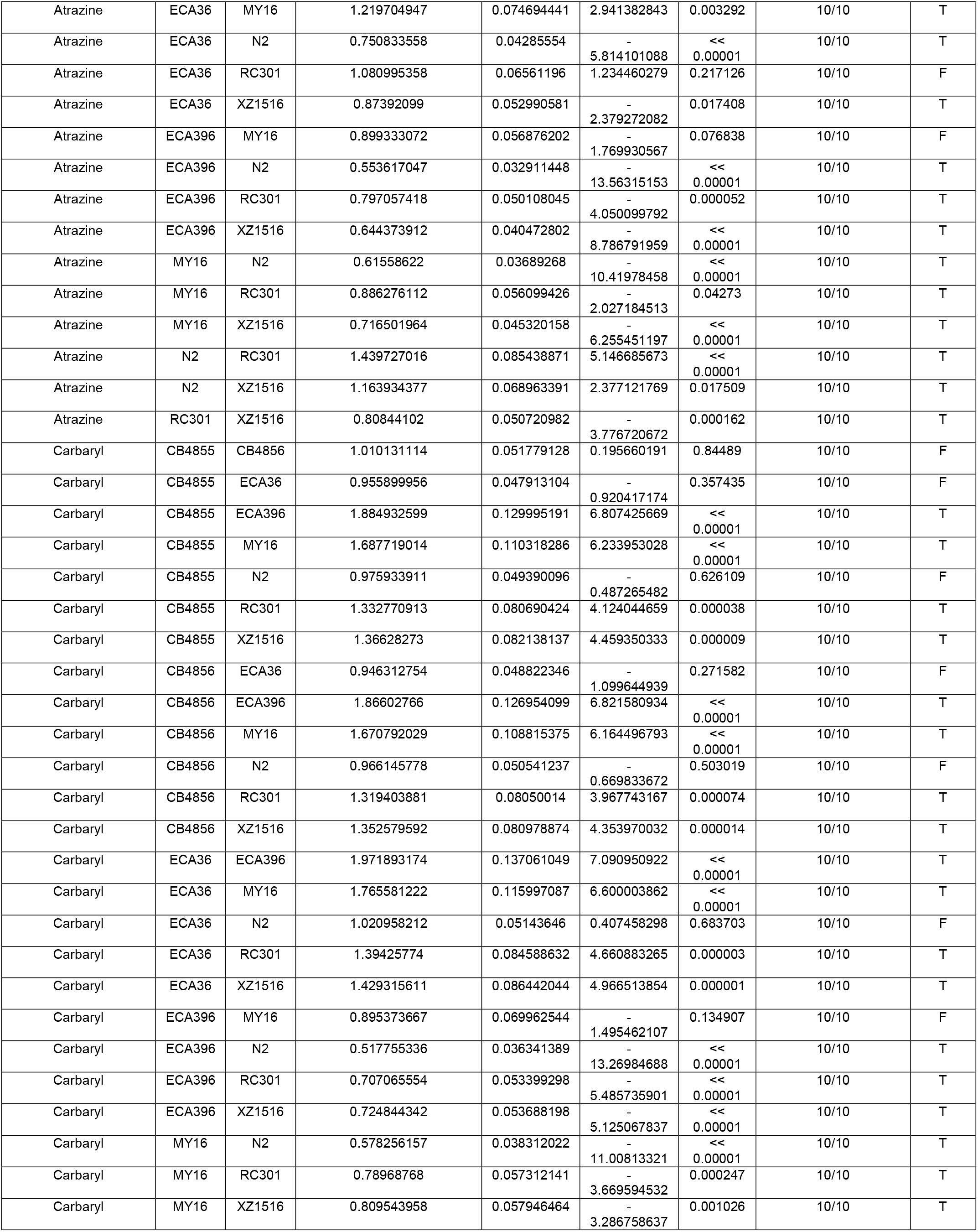
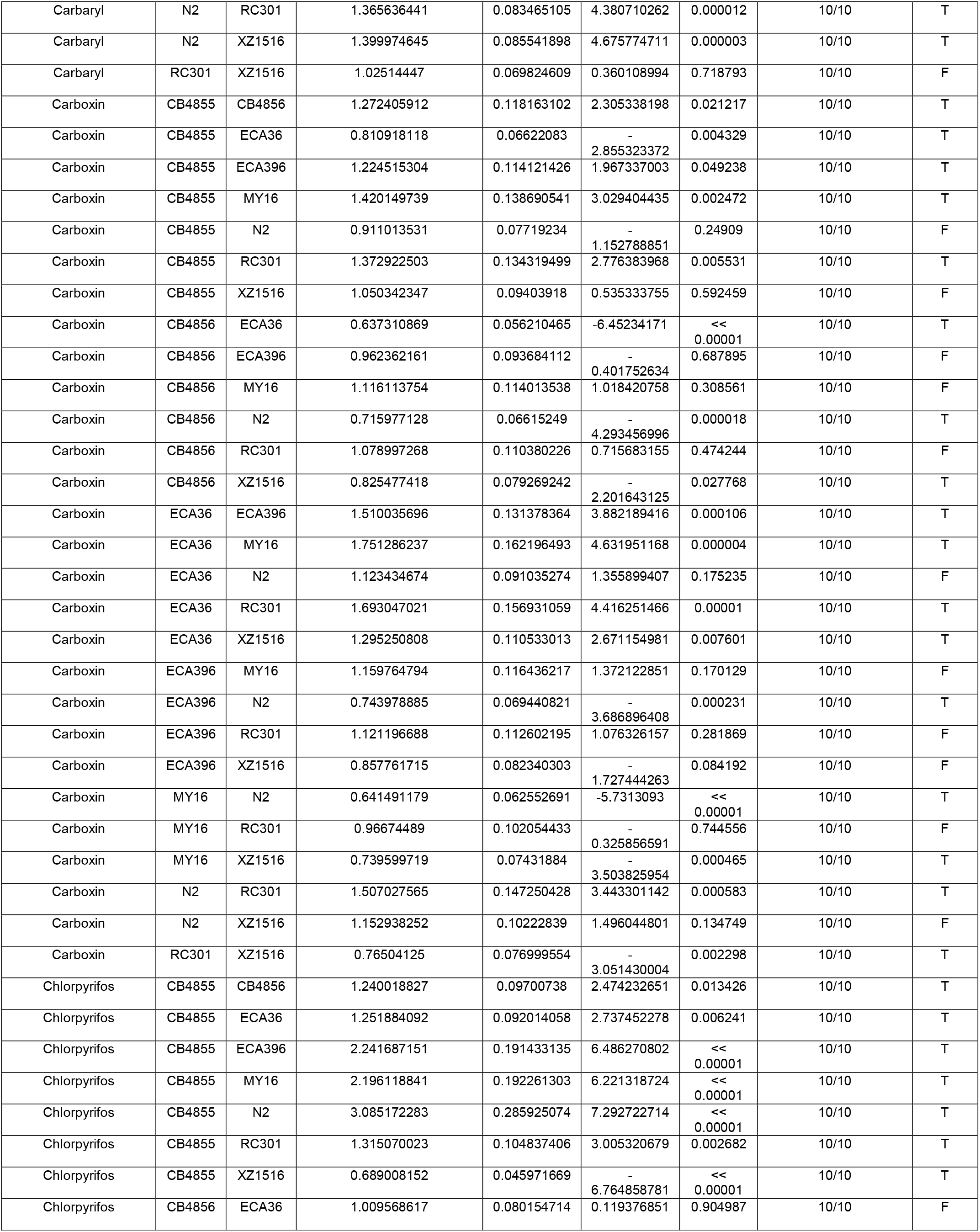
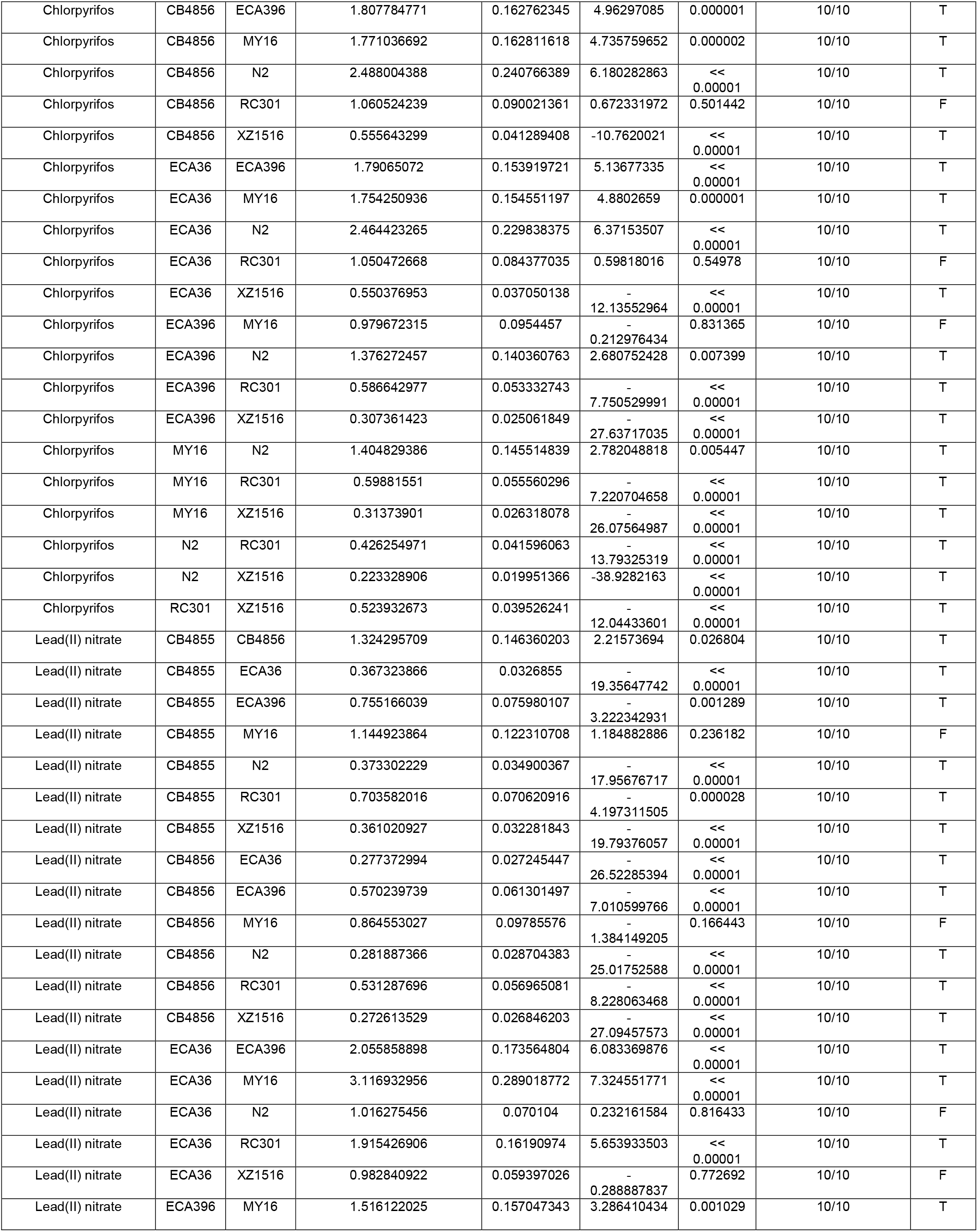
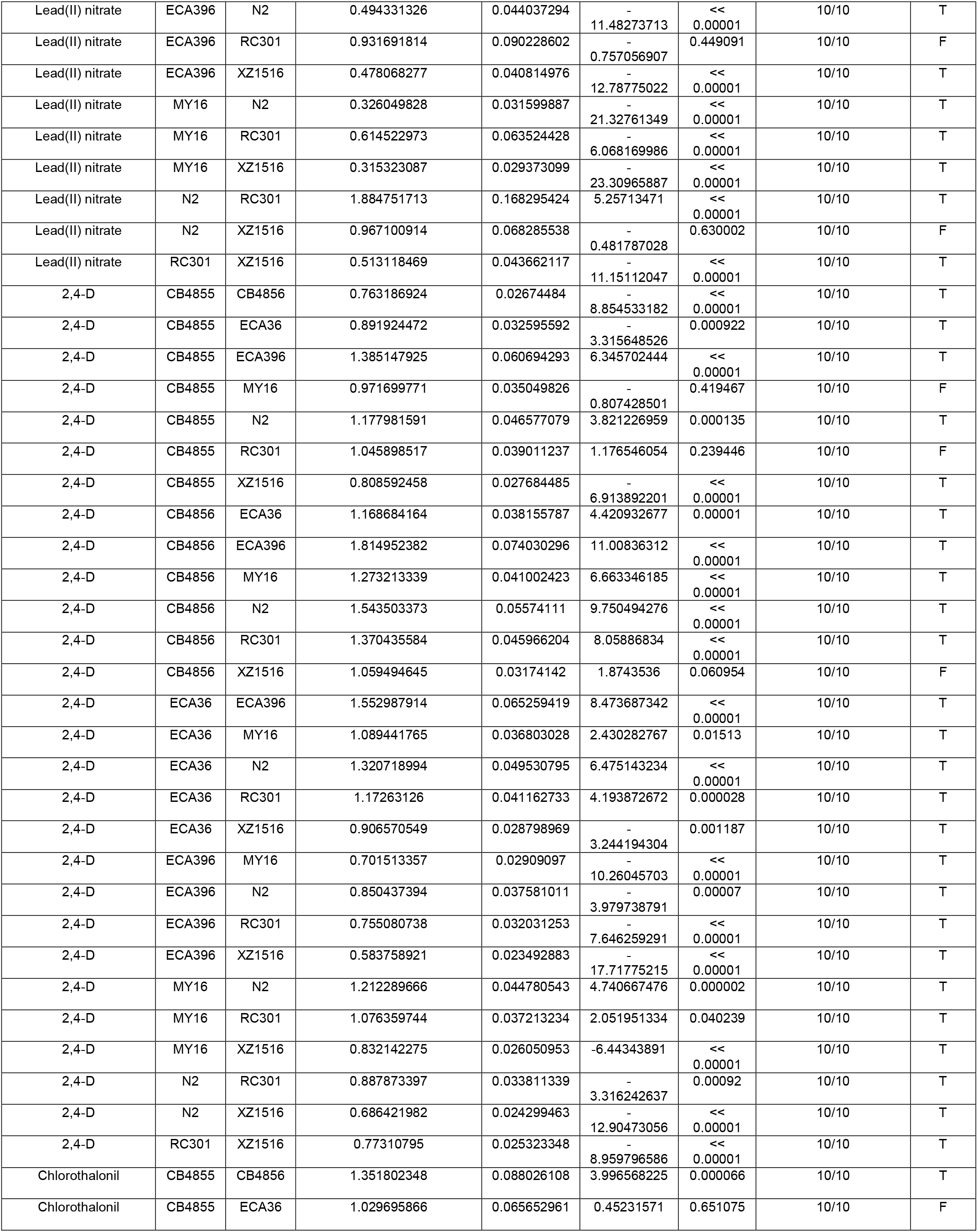
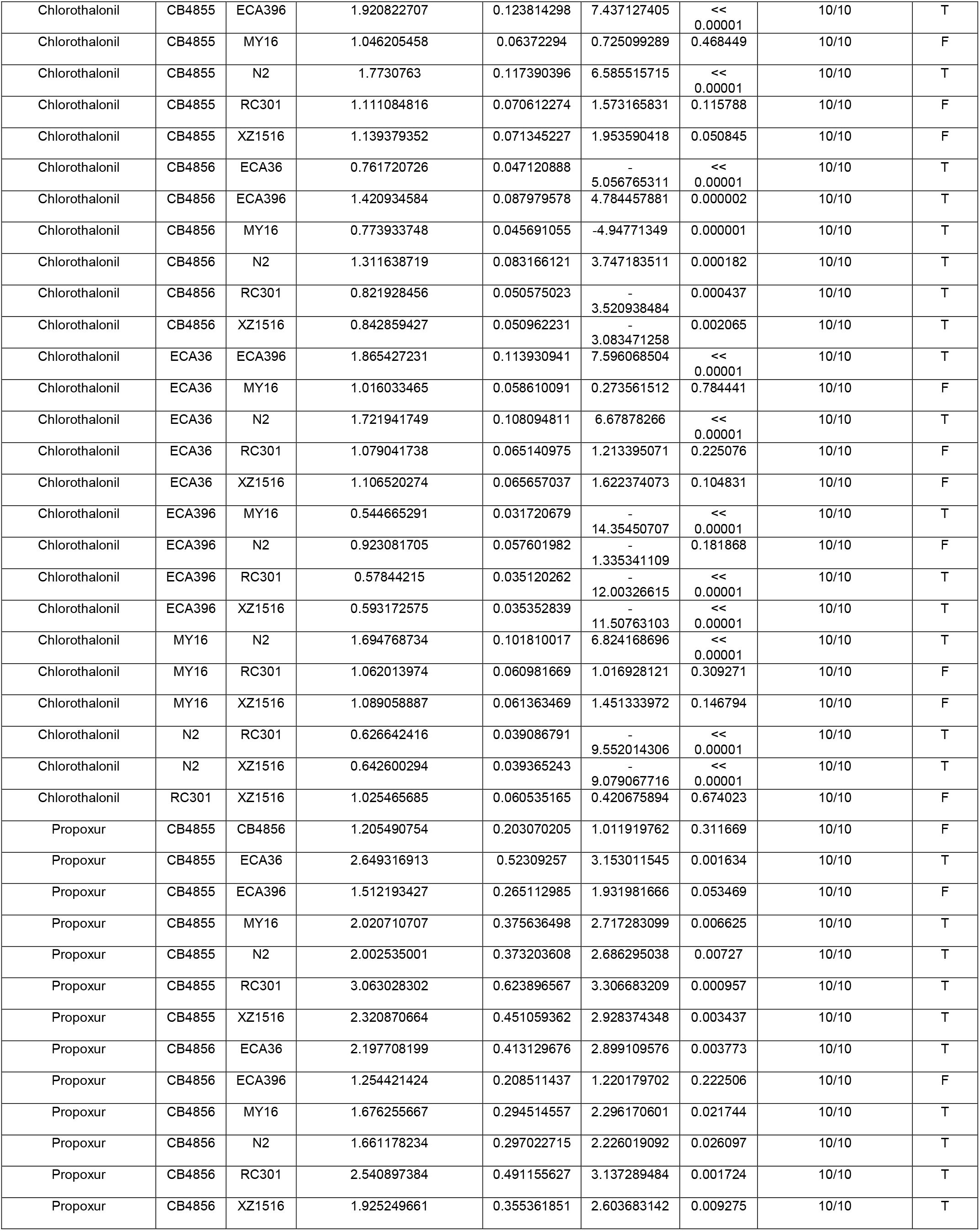
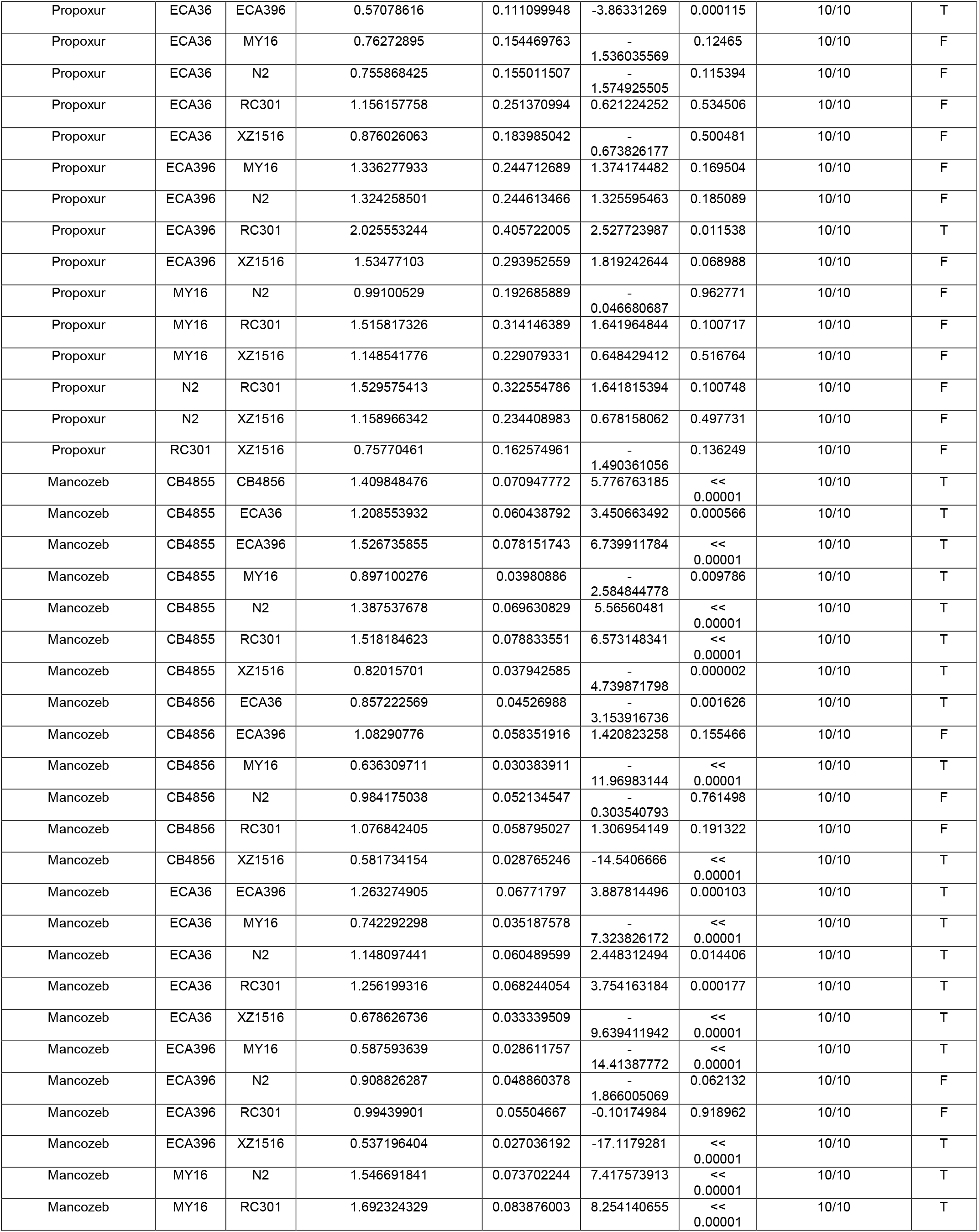
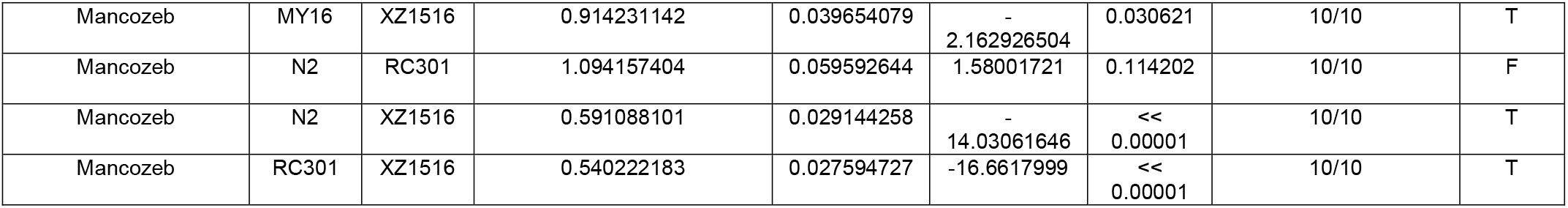
Relative potency estimates in pairwise comparisons of EC10 estimates among all strains for each toxicant.

**Supplemental Table 4.**
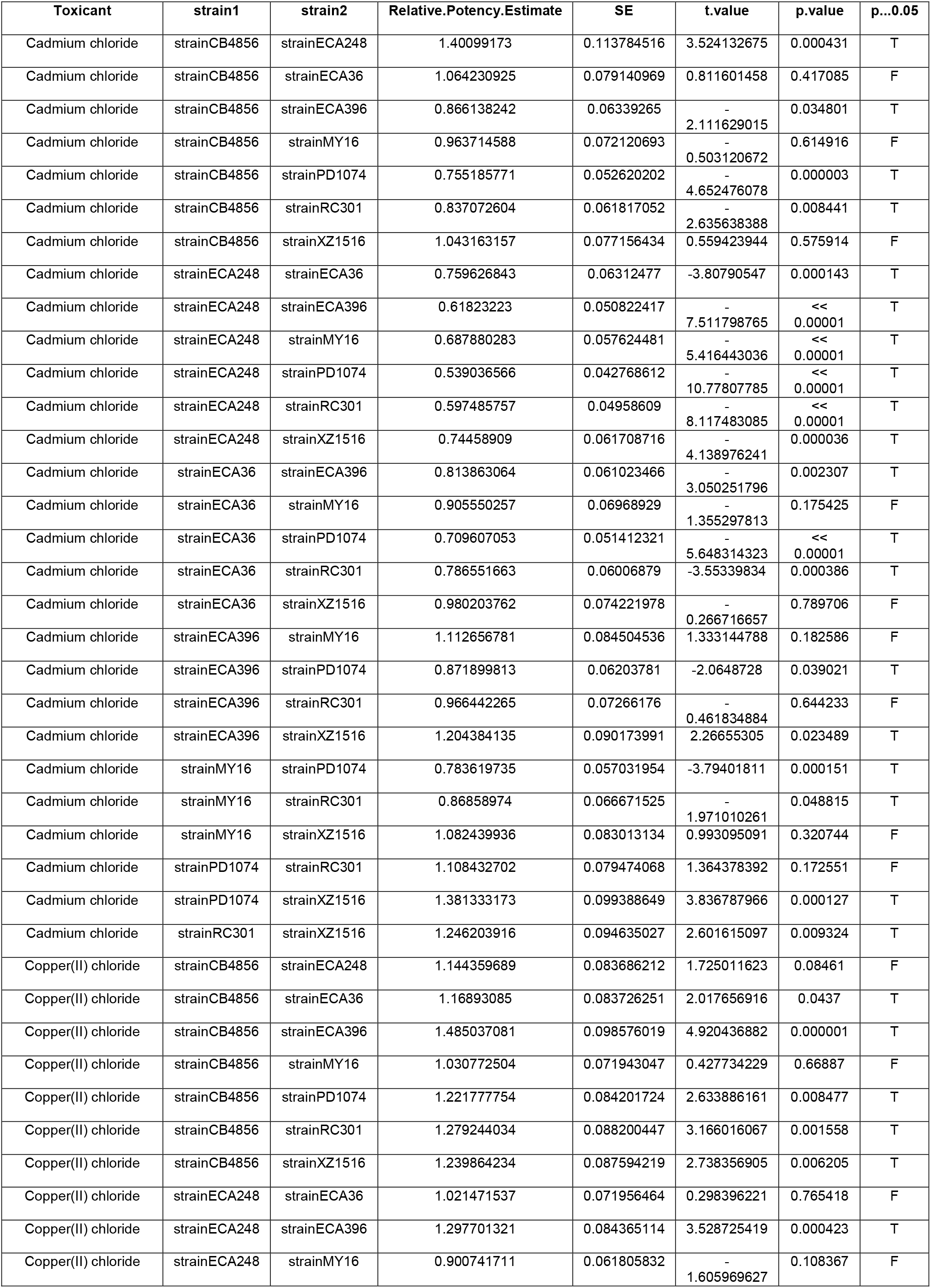
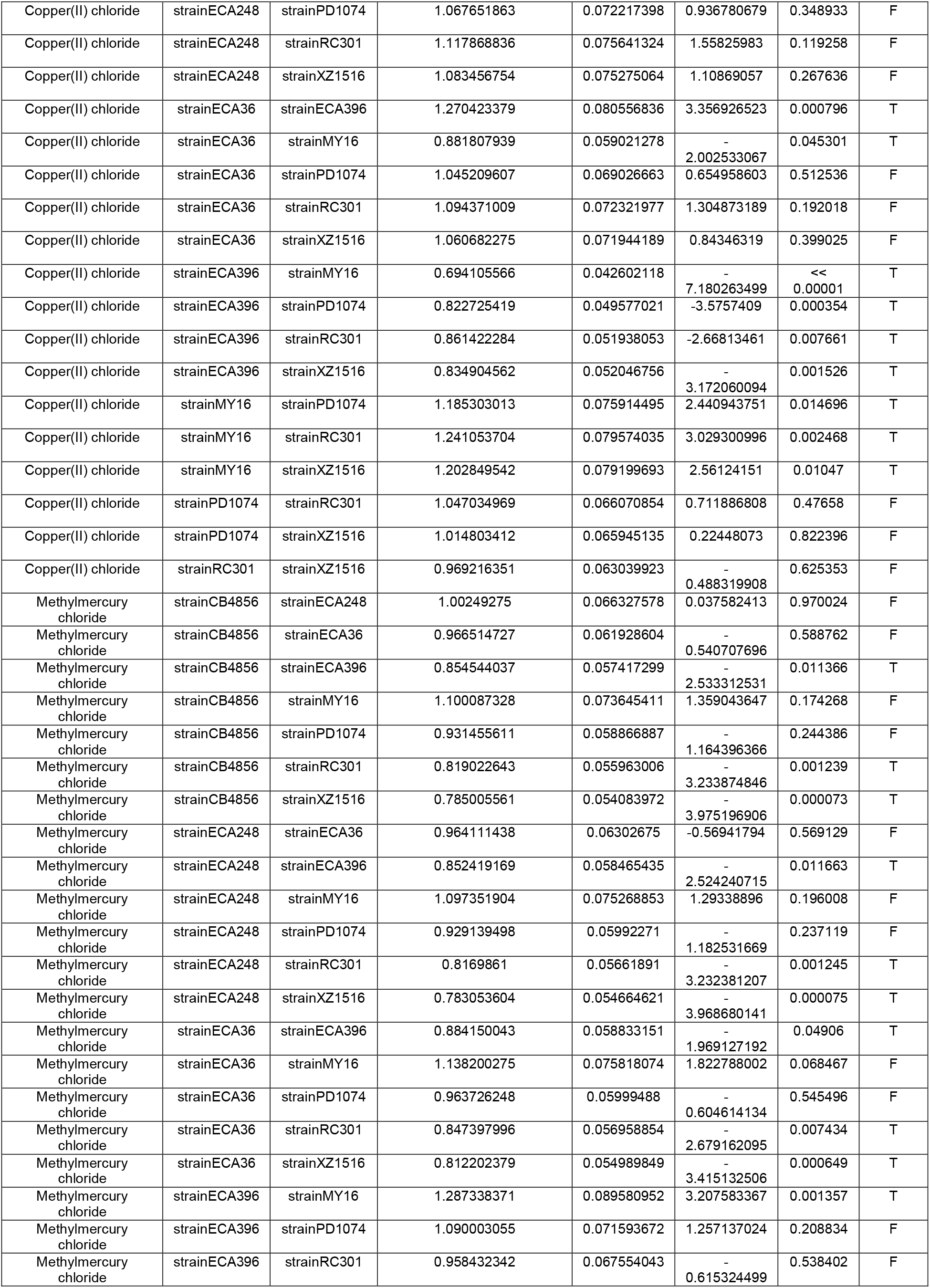
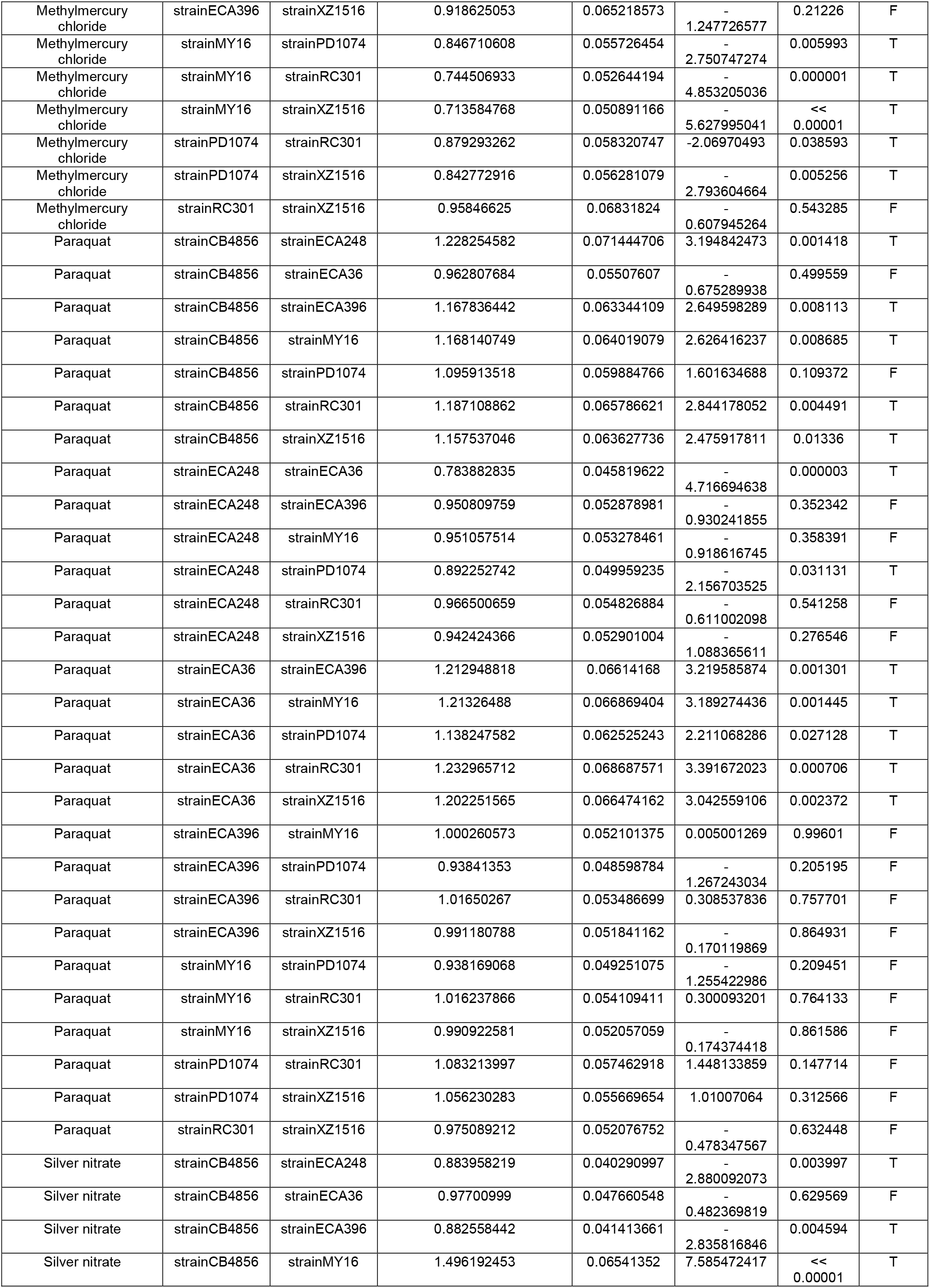
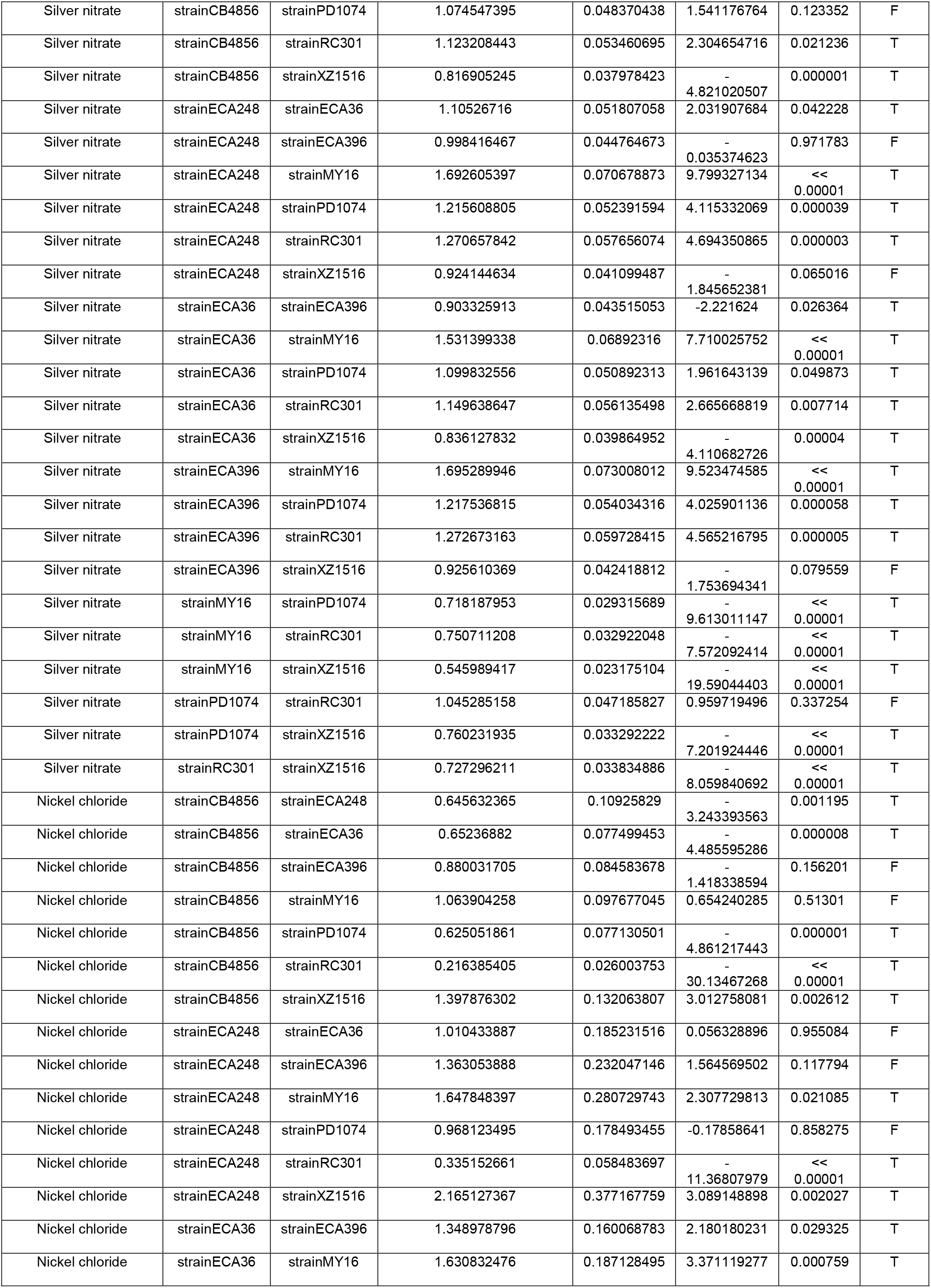
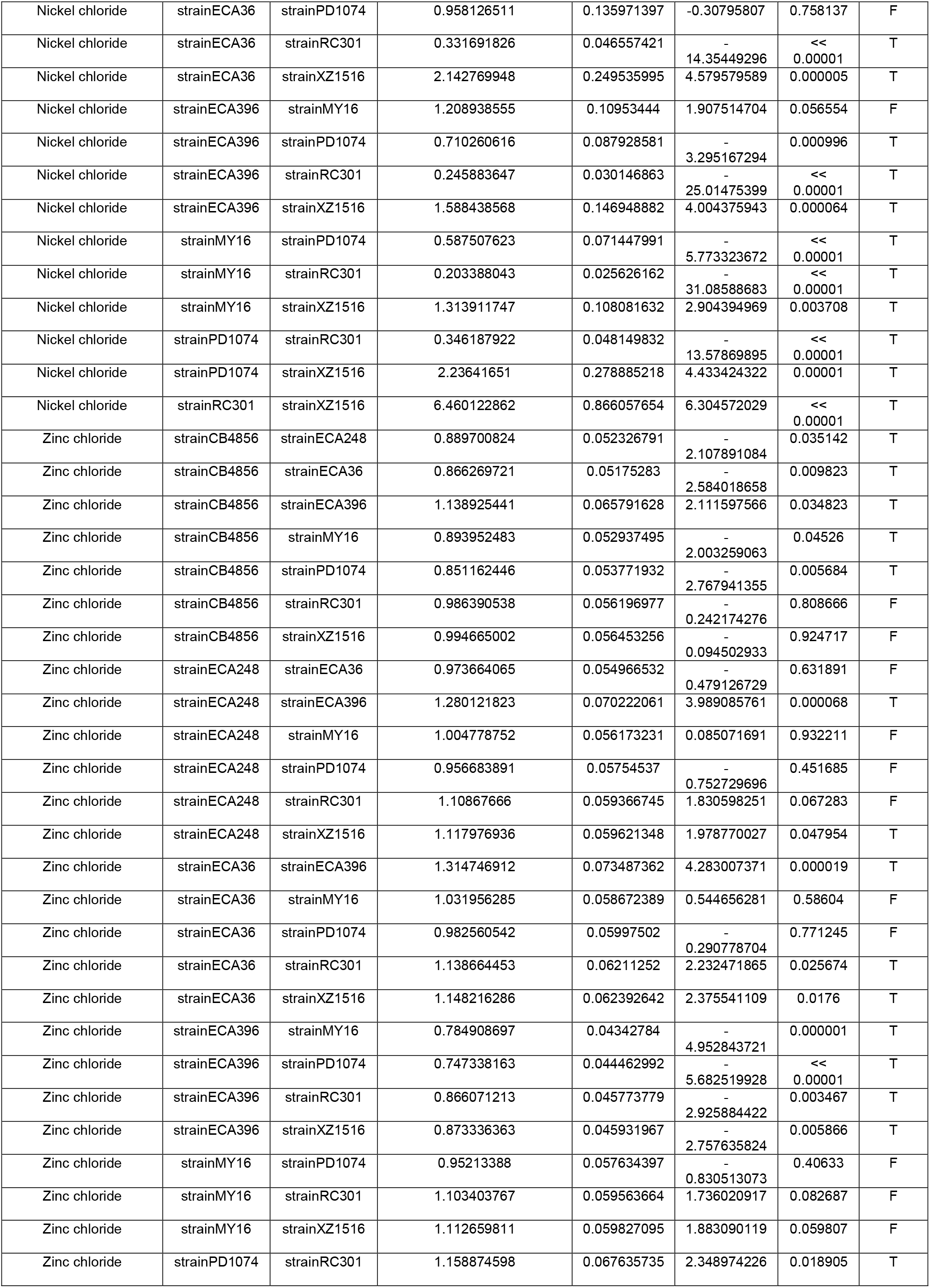
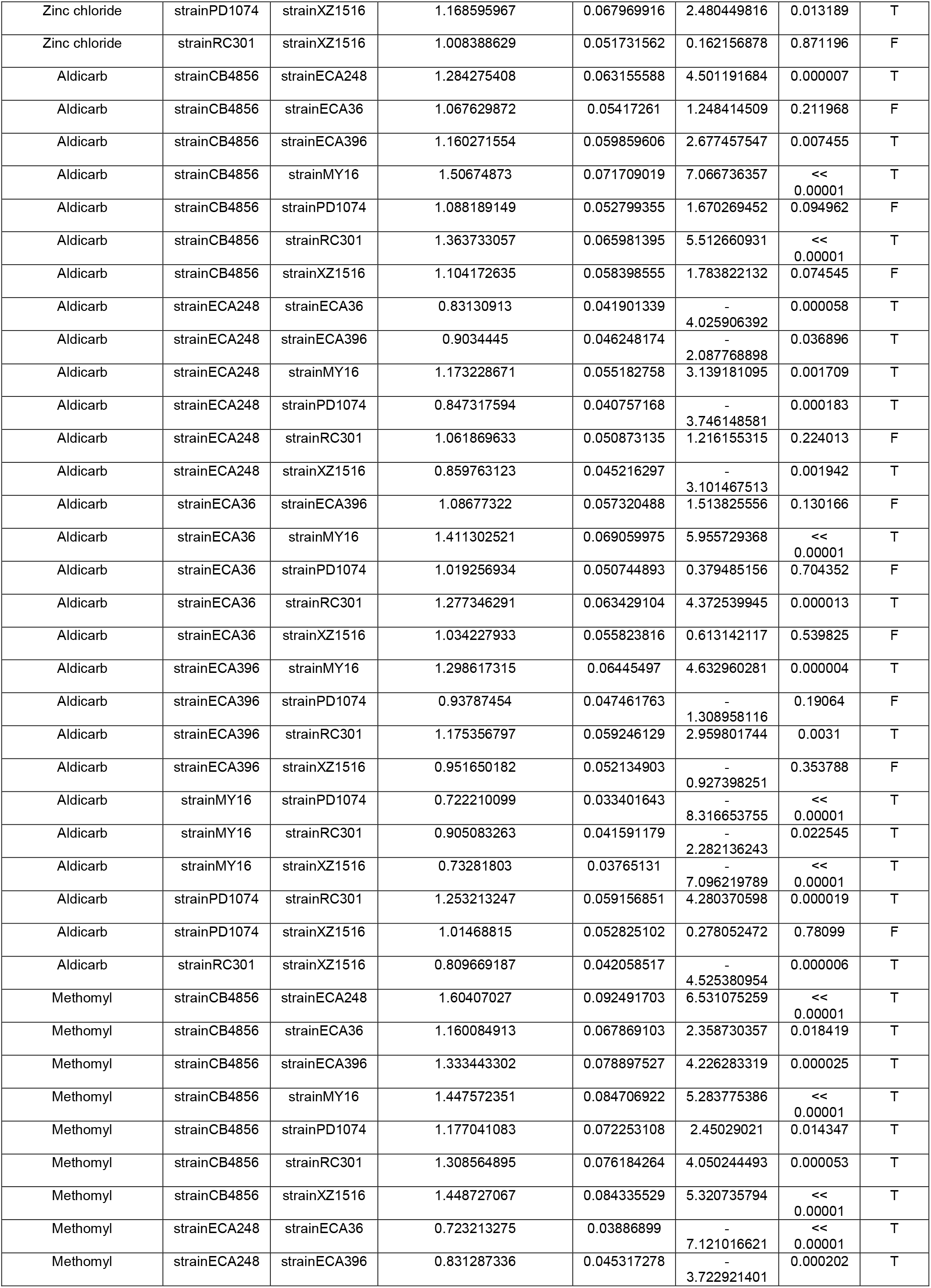
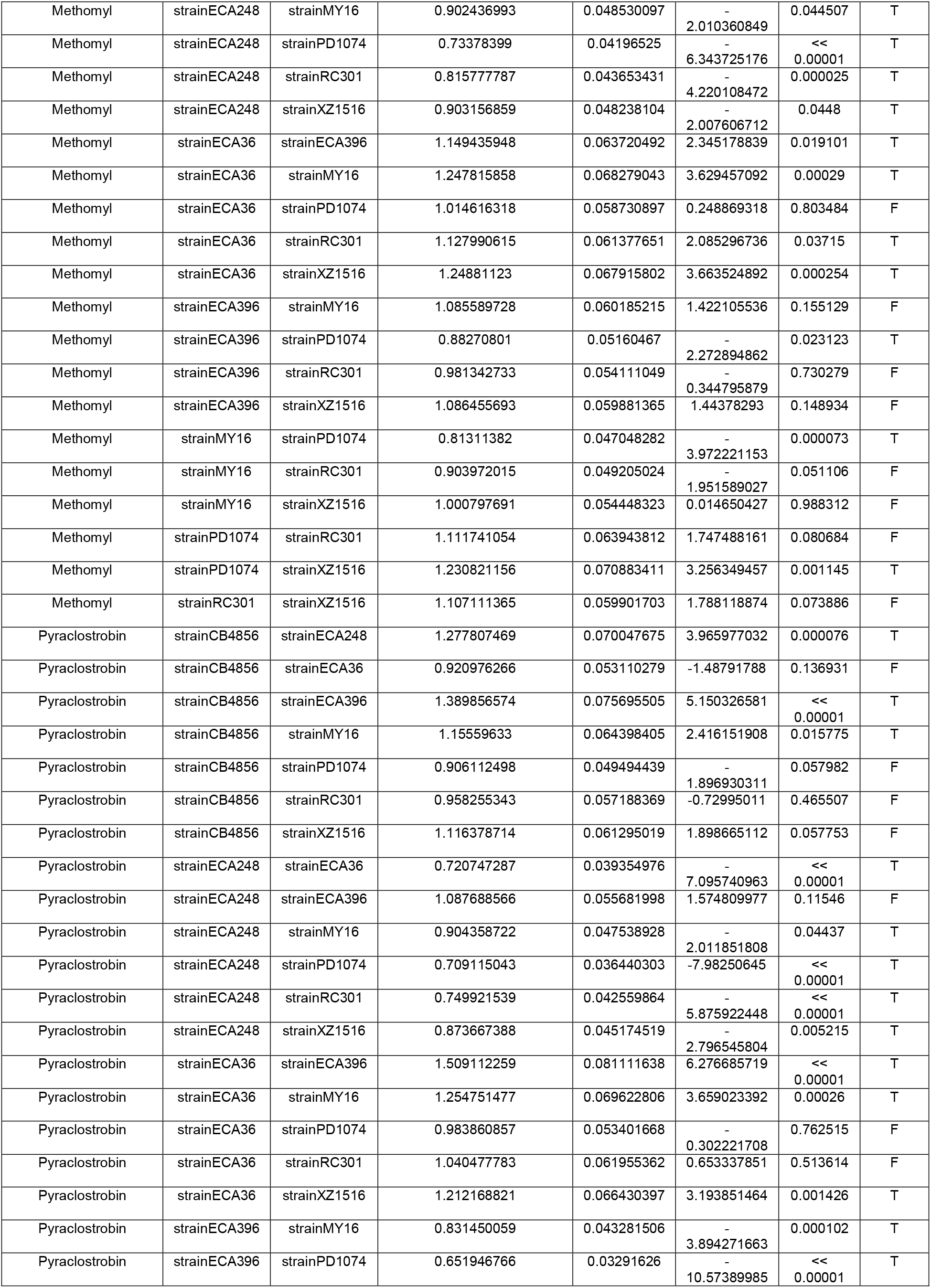
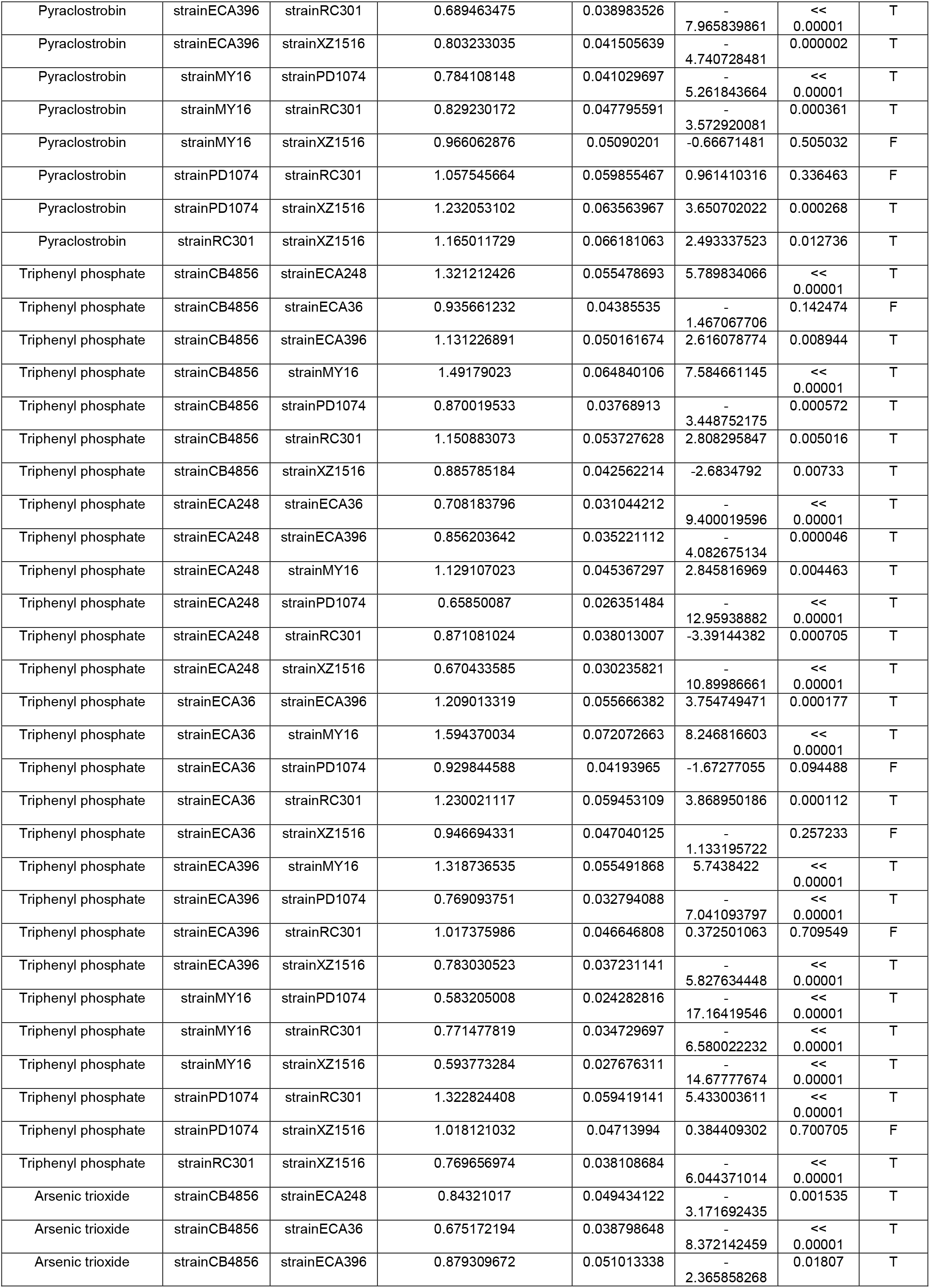
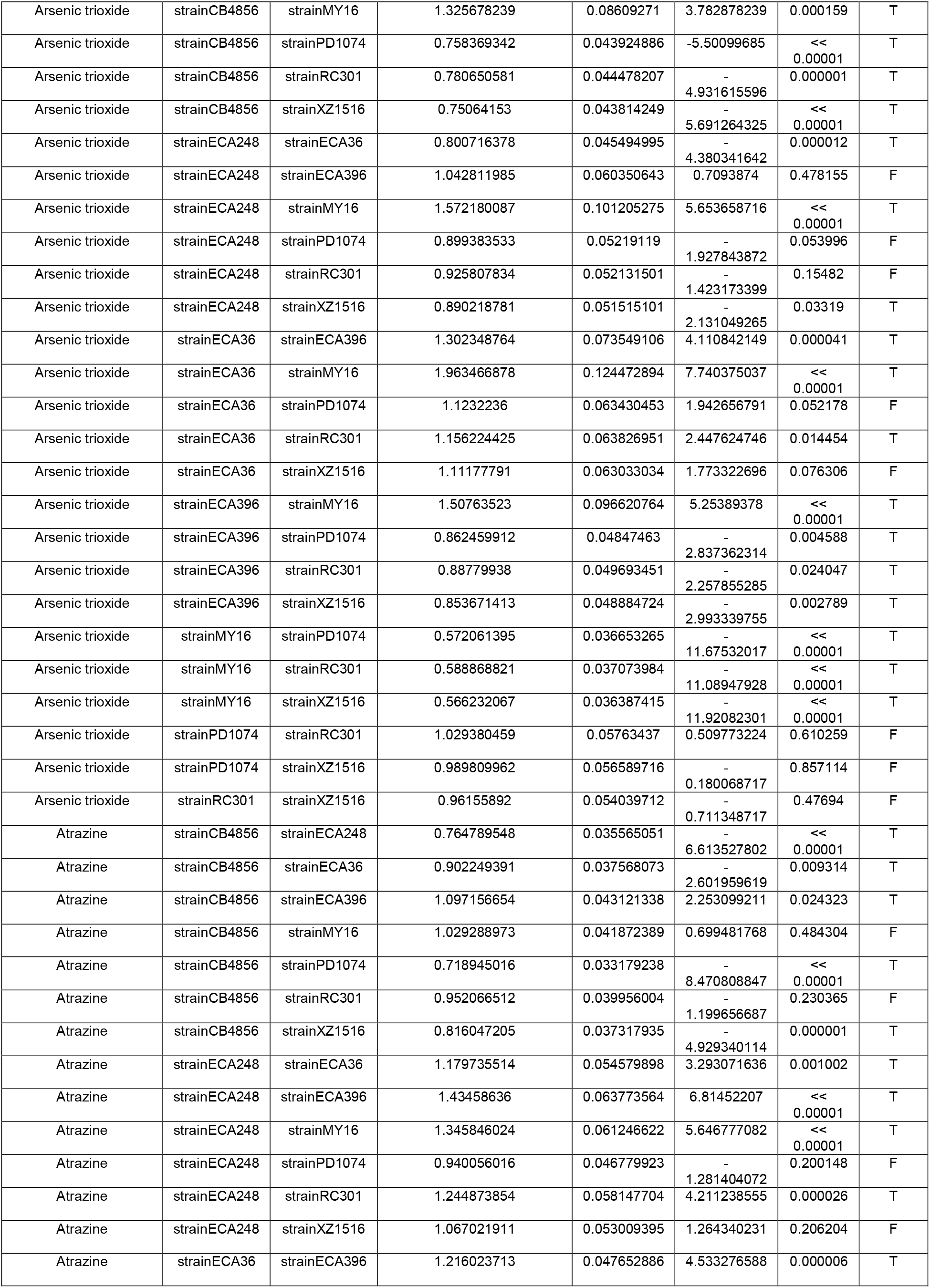
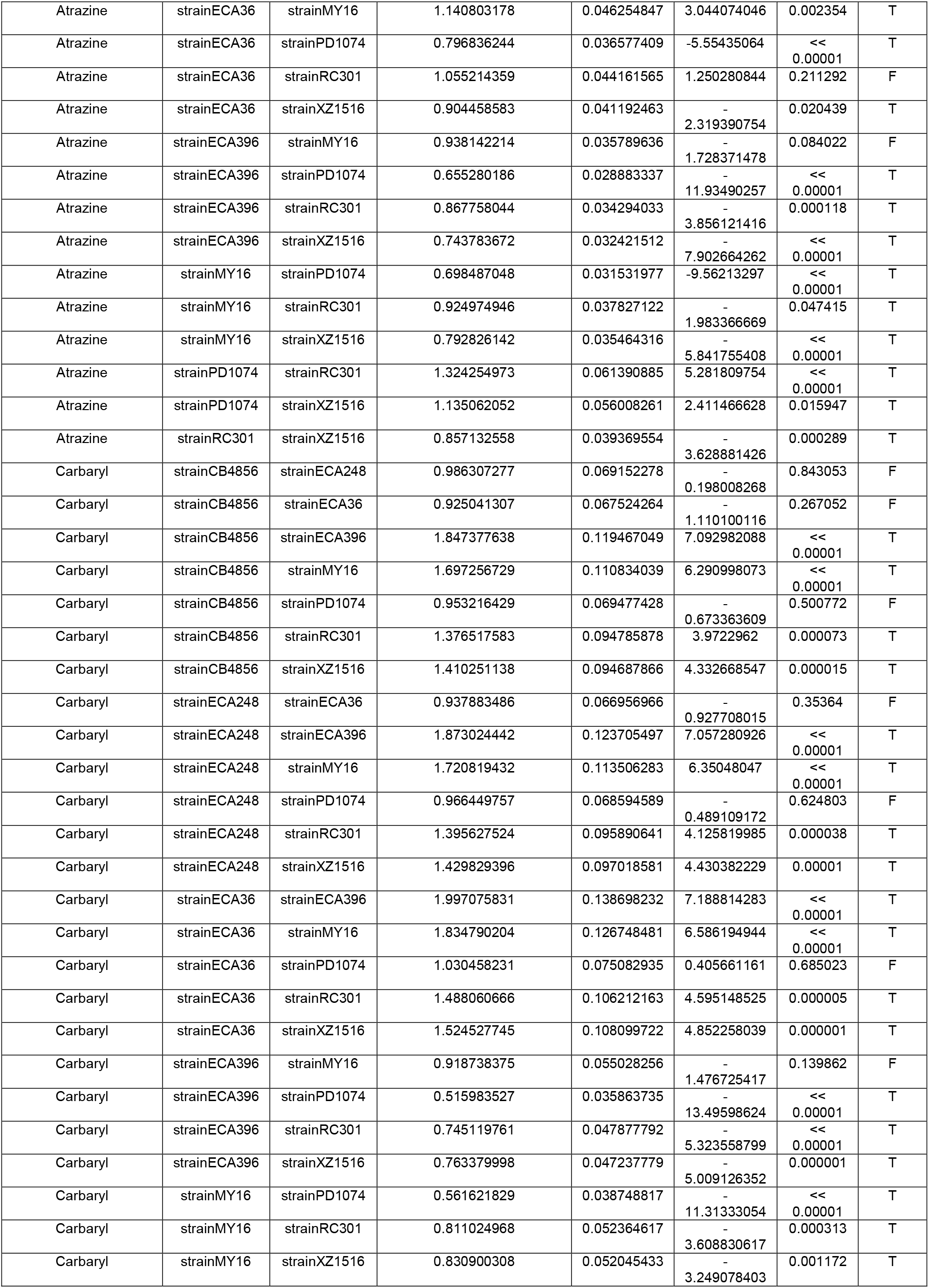
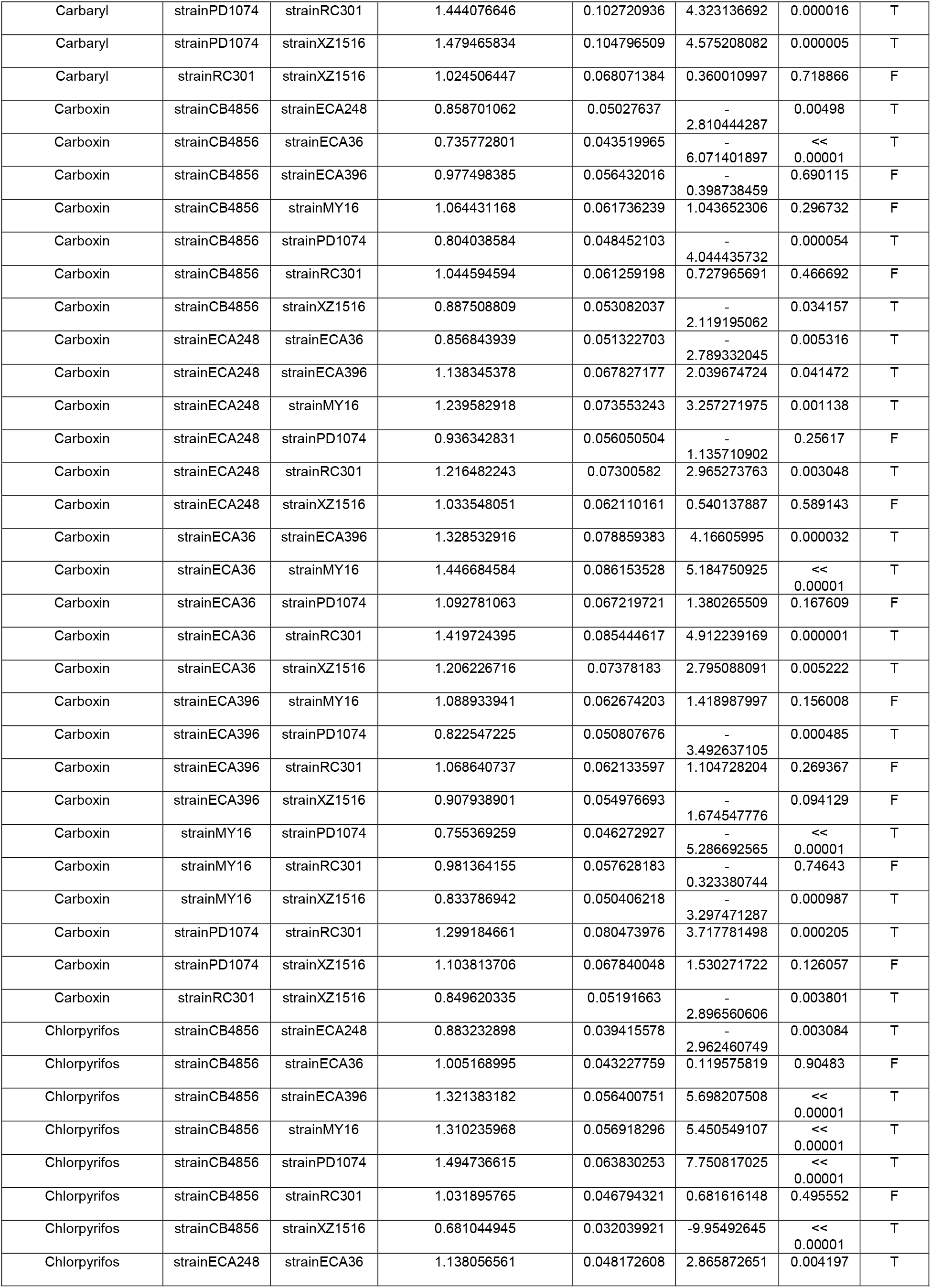
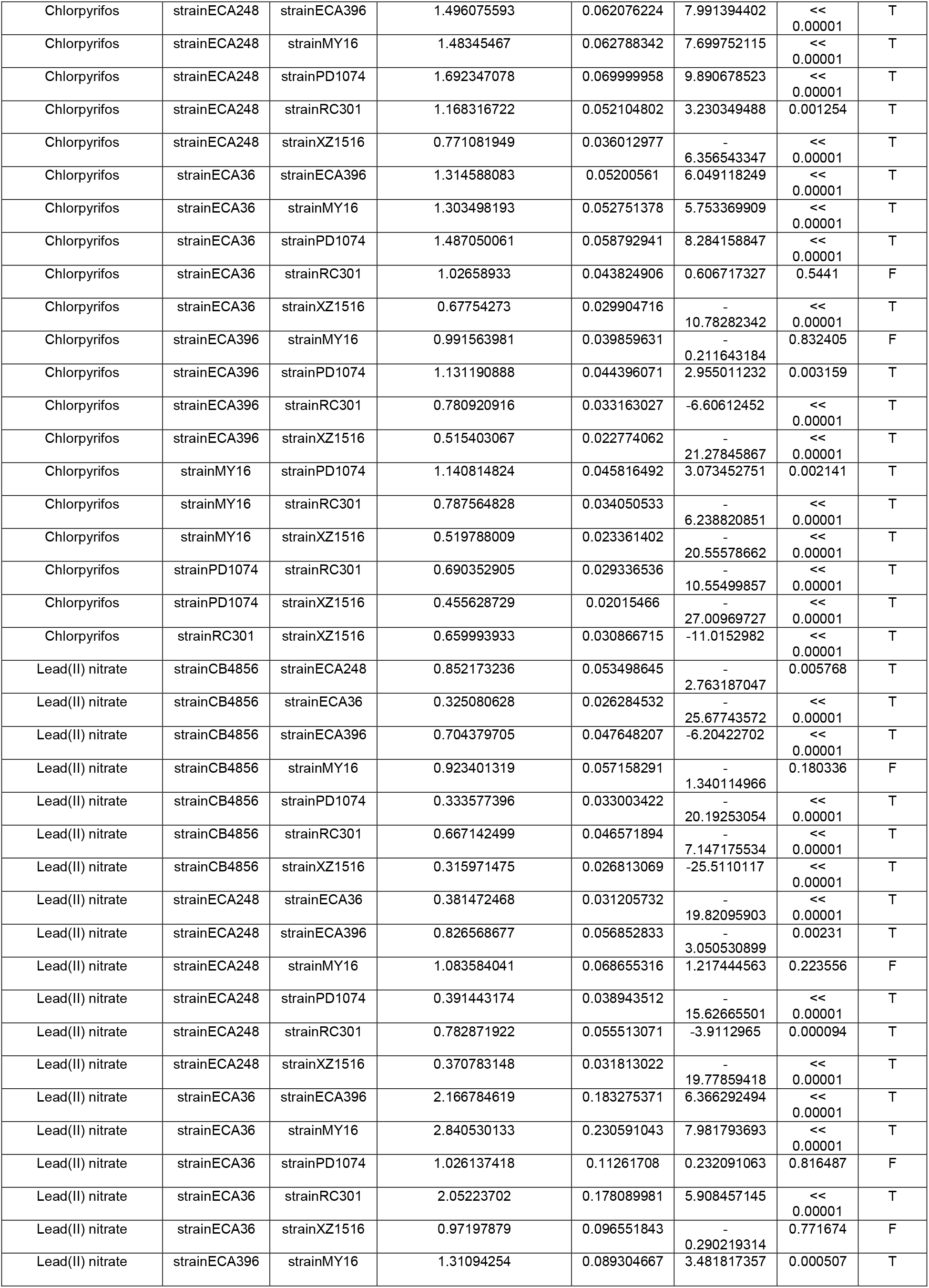
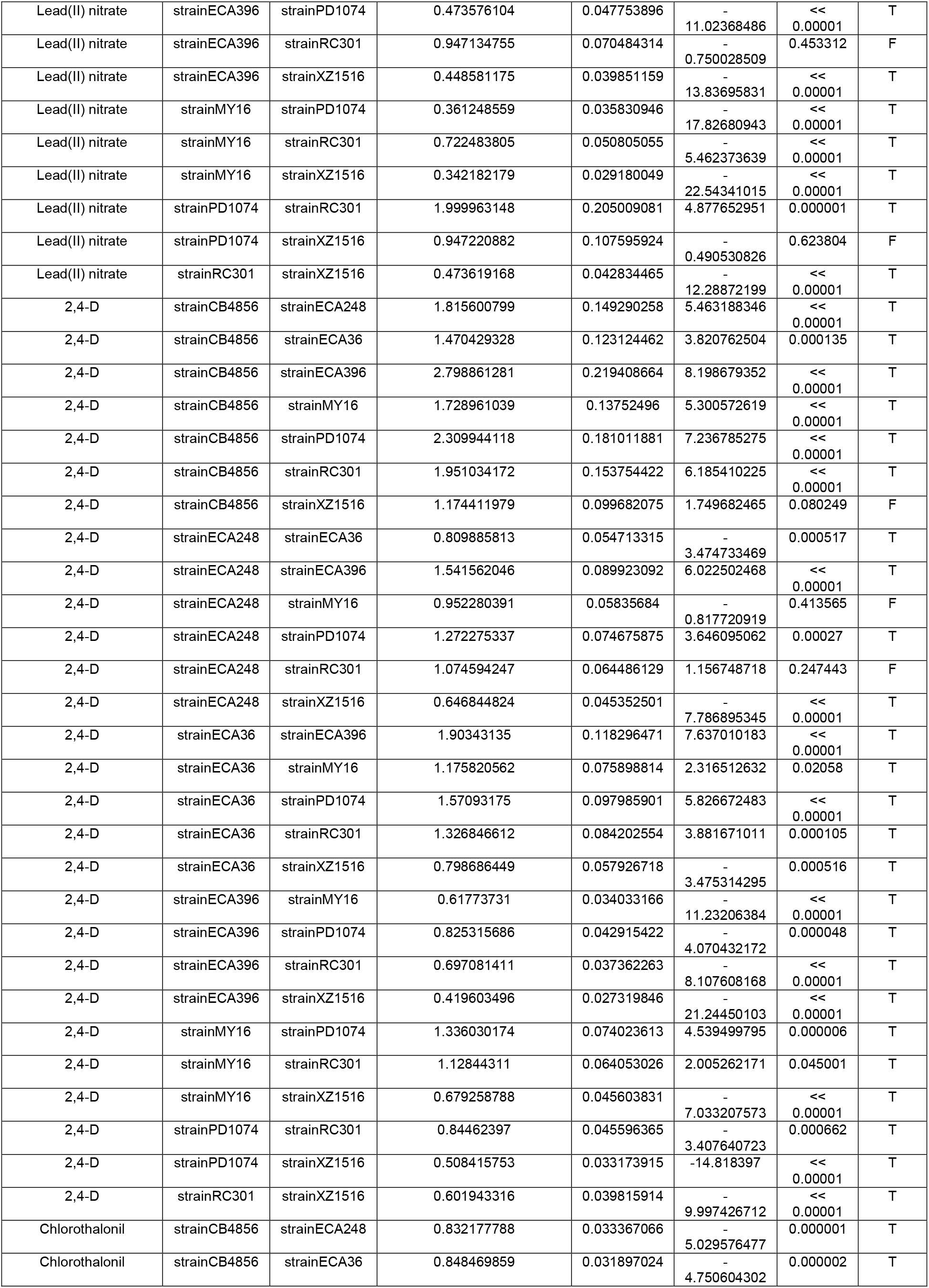
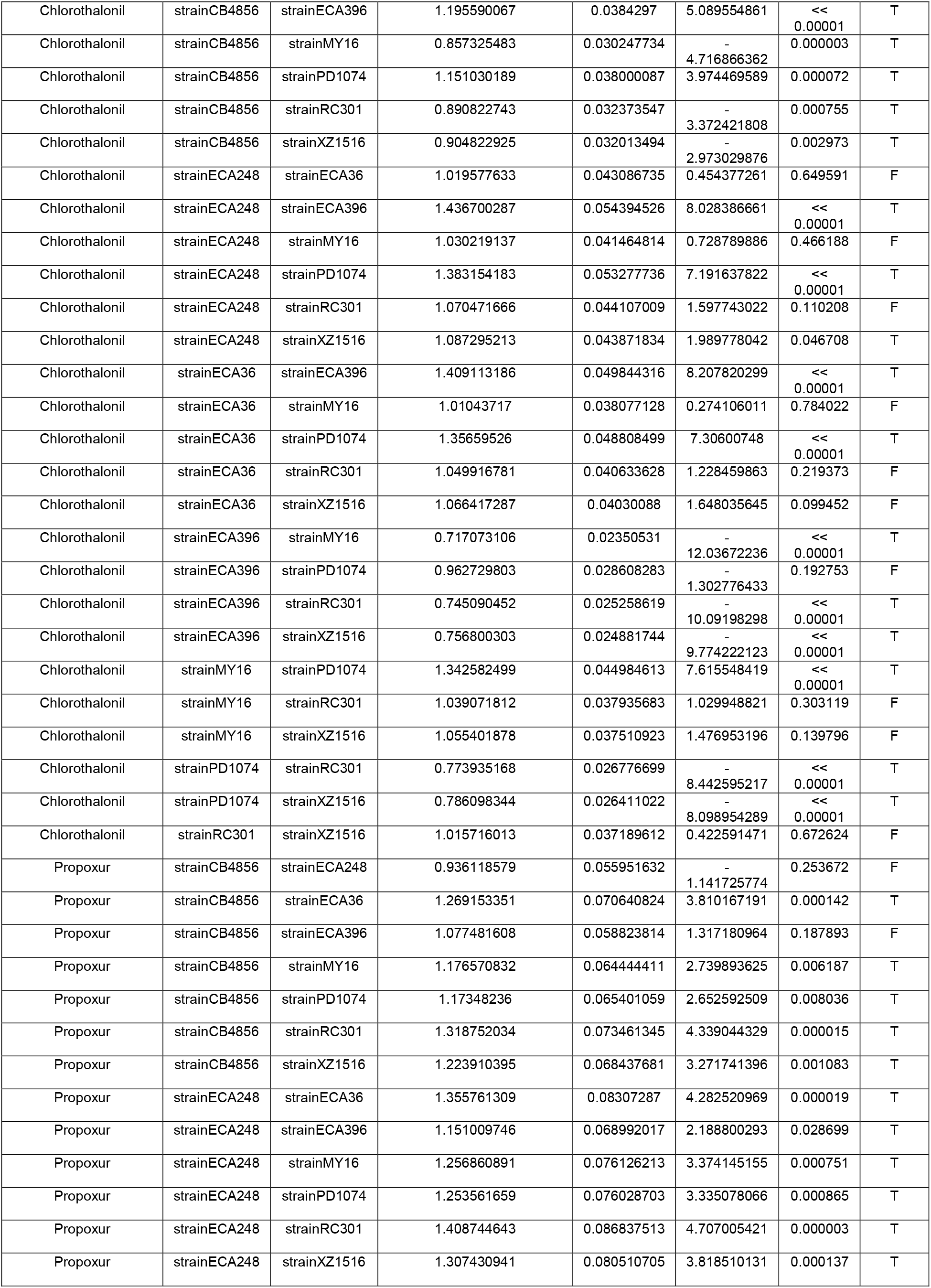
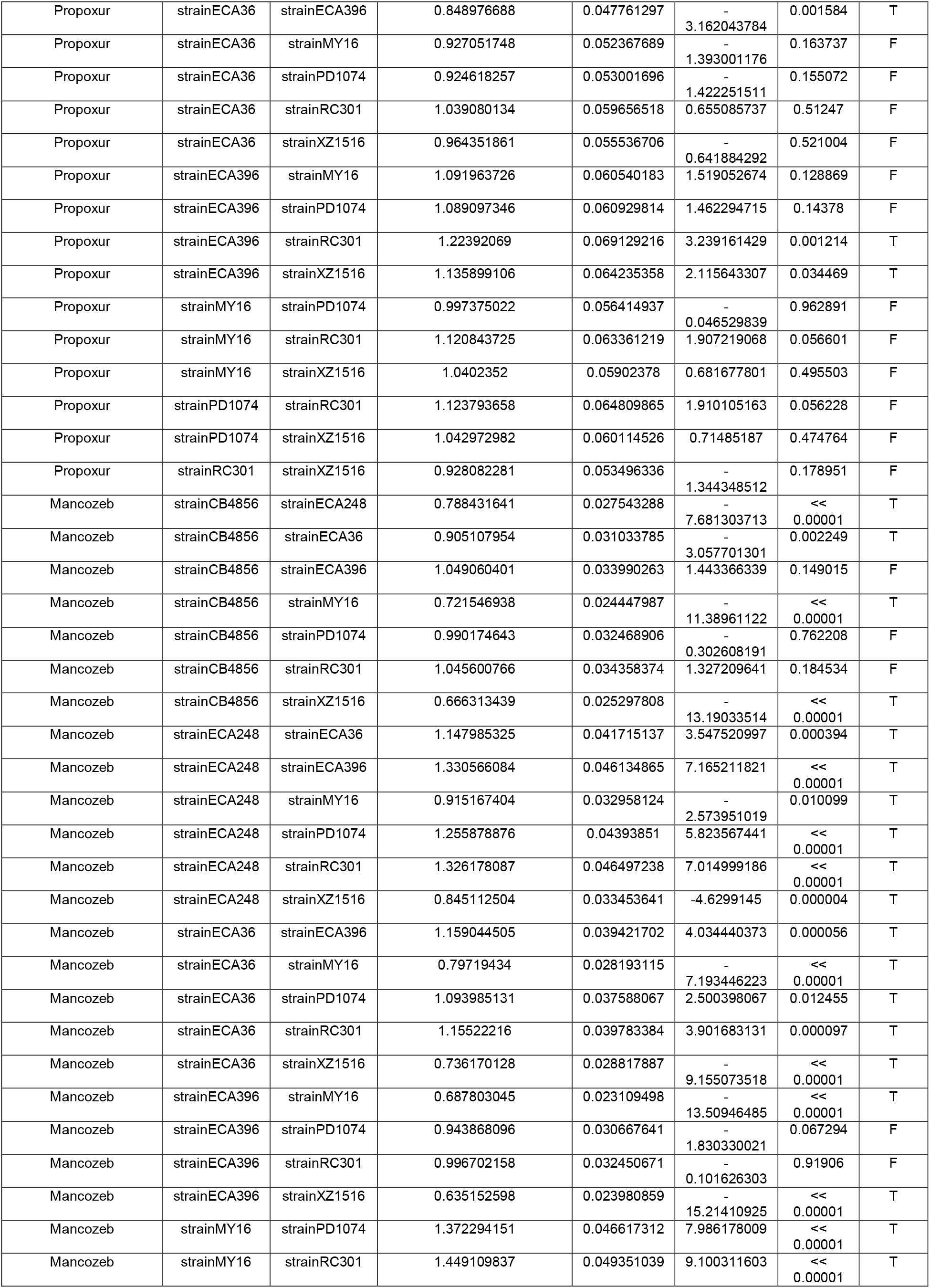
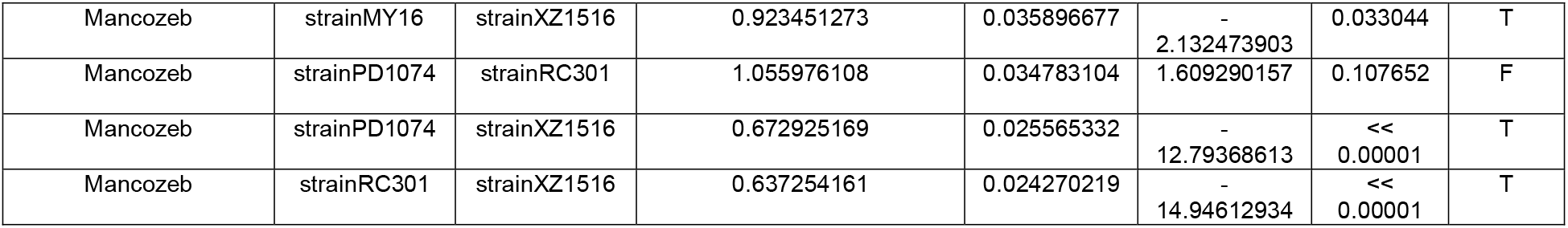
Relative potency estimates in pairwise comparisons of slope estimates among all strains for each toxicant.

